# β-cyclocitral elicits rapid signaling distinct from the excess-light response in Arabidopsis

**DOI:** 10.64898/2026.10.06.757037

**Authors:** M. Carnà, E. Tosato, P. Korwin Krukowski, A.S. Parmagnani, G. Mannino, A. Costa, S. D’Alessandro

## Abstract

β-Cyclocitral (β-cc), a β-carotene oxidation product, has been proposed as an early retrograde signal that primes protection against photooxidative stress. Its temporal relationship with photosynthetic acclimation, however, remains unresolved. Exogenous β-cc elicited cytosolic Ca²⁺ changes within minutes and induced the β-cc-responsive pAER::LUC detoxification reporter within 40 min. We exposed *Arabidopsis thaliana* plants to 1500 μmol photons m⁻² s⁻¹ under ventilation creating a high-irradiance regime without concomitant heat stress and with a relatively mild photooxidative outcome (RT-EL). Non-photochemical quenching increased, whereas PSII maximum quantum yield progressively decreased during RT-EL, with both responses approaching a plateau by 16 h. Volatile β-cc pretreatment did not improve photosynthetic performance or NPQ dynamics, whereas endogenous β-cc and β-cyclocitric acid accumulated only after prolonged RT-EL exposure (16 h). By contrast, hormone profiling and RNA-seq revealed extensive β-cc–dependent reprogramming that was largely opposite to the RT-EL transcriptional response. Together, these results reveal a kinetic separation between rapid β-cc signaling and its endogenous production. Under RT-EL major photosynthetic adjustments precede detectable β-cc accumulation, whereas β-cc triggers cellular responses within minutes once present. We therefore propose that the timing of β-cc action during excess light is determined primarily by the onset of substantial β-carotene oxidation.

## 1 Introduction

β-cyclocitral (β-cc) is a bioactive apocarotenoid derived from the oxidative cleavage of β-carotene (Carnà *et al*., 2025). Together with other β-carotene–derived volatiles such as β-ionone and dihydroactinidiolide, β-cc has been reported to modulate plant physiology and has attracted interest as a potential biostimulant to prime tolerance to abiotic and biotic stress (Shumbe *et al*., 2014; Jia *et al*., 2022; Felemban *et al*., 2024).

The physiological impact of β-cc differs markedly across photosynthetic lineages: it proved toxic to cyanobacteria, whereas it had little to no effect in algae, and it has mainly been associated with enhanced stress tolerance in land plants (Tomita *et al*., 2016; D’Alessandro *et al*., 2018; Roach *et al*., 2022; Wang *et al*., 2024). Interestingly, in both cyanobacteria and algae, β-cc has been reported to repress chlorophyll biosynthesis, a pathway that is partially conserved in plants (Ramel *et al*., 2012). In addition, β-cc also affects photosynthetic performance in *Chlamydomonas reinhardtii*: the maximum quantum yield of PSII decreased after a volatile β-cc treatment (60 µmol) under very low light (2 μmol photons m^−2^ s^−1^), and non-photochemical quenching (NPQ, measured after 2 minutes at 750 μmol photons m^-2^ s^-1^) was dramatically reduced by treatment with as little as 3 µmol volatile β-cc treatment in a 30-mL airspace (Roach *et al*., 2022). In the cyanobacterium *Microcystis aeruginosa,* the strong toxicity of β-cc might mediate a quorum-sensing-like mechanism and trigger distinct cell-death pathways depending on concentration and delivery mode (Wang *et al*., 2024). In that study, β-cc diluted in DMSO was administered directly to the growth medium, at concentrations ranging from 600 to 1800 μmol L^-1^. The programmed cell death (PCD)-inducing behavior of β-cc (400 μmol L^-1^) and β-ionone (200 μmol L^-1^) in water has also been reported in algae (*Chlamydomonas reinhardtii)* and aquatic plants (*Lemna turionifera*) (Sun *et al*., 2020; Pan *et al*., 2024). Accordingly, β-cc has been suggested to participate in the ecological balance among cyanobacteria, algae, and plants in aquatic environments, where β-cc can be actively released into the surrounding medium (Lin *et al*., 2024).

In terrestrial environments, the situation seems to be quite the opposite: β-cc is predominantly reported as a product of photooxidative processes linked to excessive excitation of photosystem II (PSII), acting as a signaling molecule that primes acclimation and stress tolerance (Ramel *et al*., 2012; Lv *et al*., 2015; D’Alessandro *et al*., 2018; Deshpande *et al*., 2021; Mitra *et al*., 2021; Avesani *et al*., 2025). β-cc treatments at various concentrations have been shown to induce an extensive transcriptional and metabolic reprogramming in several species, including *Arabidopsis thaliana*, *Solanum lycopersicum*, *Oryza sativa* and *Vitis vinifera* (Deshpande *et al*., 2021; Taniguchi *et al*., 2023; Avesani *et al*., 2025; Tiwari *et al*., 2025).

Mechanistically, β-cc has been linked to increased tolerance to excess light (EL) under controlled low-temperature conditions through METHYLENE BLUE SENSITIVITY 1 (MBS1)–dependent regulation of singlet-oxygen–responsive genes (SORG), as well as through the SCARECROW-LIKE 14 (SCL14)–dependent xenobiotic detoxification pathway (Shumbe *et al*., 2017; D’Alessandro *et al*., 2018). The latter involves typical phase I (oxidation/reduction, aldo-keto reductases (AKR), alkenal reductases (AER), aldehyde dehydrogenase (ALDH), peroxidases and cytochrome P450 oxidases (CYP)) and phase II (conjugation; glycosyltransferases (UGT) and glutathione S-transferases (GST)) enzymes commonly recruited during stress responses (Fode *et al*., 2008; D’Alessandro *et al*., 2018, 2020; D’Alessandro and Havaux, 2019; Tiwari *et al*., 2025).

β-cc has also been reported to interact with hormone pathways: it is induced by jasmonic acid (JA) in rice, and treatment with volatile β-cc at 10 μmol per liter repressed abscisic acid (ABA) accumulation and synthesis and induced JA signaling without the accumulation of jasmonate itself (Taniguchi *et al*., 2023). Furthermore, a role for salicylic acid (SA) has been described in the β-cc-induced protection against excessive light at 18 °C in Arabidopsis (300 μmol volatile in 22 liters) (Lv *et al*., 2015). Finally, volatile β-cc treatment (1500 μmol in 3 liter) has been proposed to inhibit 1-deoxy-D-xylulose 5-phosphate synthase (DXS), the rate-limiting enzyme of the MEP pathway upstream of isoprenoid biosynthesis, in Arabidopsis (Mitra *et al*., 2021).

Despite these advances, little is known about the kinetics and specificity of β-cc signaling, or about the pathways underlying its beneficial effect in land plants. In this work, we established a high-irradiance regime in well-watered, ventilated adult Arabidopsis plants in which leaf temperature remained stable and the photooxidative outcome was mild. We combined time-resolved chlorophyll-fluorescence imaging with β-cc quantification, reporter assays, Ca²⁺ imaging, hormone measurements, and RNA sequencing. This design allowed us to compare the kinetics of photosynthetic acclimation with endogenous apocarotenoid accumulation and with the response to externally supplied β-cc.

Our results reveal a temporal hierarchy. Photosynthetic adjustments were established during the first hours of excess light, whereas endogenous β-cc and β-cyclocitric acid accumulated later. Exogenous β-cc nevertheless triggered detectable responses within minutes. We therefore propose that β-cc becomes increasingly relevant when sustained excitation produces stronger β-carotene oxidation and a higher photooxidative load.

## 2 Material and Methods

### 2.1 Plant Growth conditions and treatments

*Arabidopsis thaliana* ecotype Col-0 (N70000) was used as wild type and was obtained from The Nottingham Arabidopsis Stock Centre (NASC). The pAER::LUC reporter line was kindly provided by Michel Havaux (Rac *et al*., 2021). In each experiment, the seeds for each line were derived from the same batch of plants grown together. For growth on plates, seeds were surface-sterilized with 70% ethanol and 0.01% Triton X-100, rinsed with 100% ethanol, and allowed to dry and then placed on square culture plates containing 45 mL of 0.5× Murashige and Skoog salts (Duchefa), 0.5 g L^−1^ MES, and 0.8% plant agar (Duchefa), adjusted to pH 5.7 with KOH, containing 50 μM Luciferin (Clinisciences). Plates were placed at 4°C for 2 days in darkness and then transferred to controlled growth conditions with 12 h/12 h light/dark photoperiod at 24°C day/20°C night. For nonsterile growth, Arabidopsis plants were grown in soil in a controlled environment at 130 μmol m^−2^ s^−1^ illumination with a 12-hour/12-hour photoperiod at 24°C day/20°C night. Both growth light and excess light were provided by 4000 K LED lamps.

Plants were grown for 4 or 5 weeks before treatments and exposed to excess light stress by transferring them to a system in which PPFD was set at 1500 µmol m^−2^ s^−1^, with water available in the saucer, and constant air flow at room temperature (26 °C, leading to an average leaf temperature of 25 °C).

90% β-cc was obtained from Thermo Fisher Scientific (B25700.09, lot. 10237632). 96% βI was obtained from Thermo Fisher Scientific (297131000, lot. A0461319). 99% meSA was obtained from Sigma-Aldrich. Classical β-cc pretreatment was performed by exposing the plants for 4 h to volatile β-cc in a transparent desiccator (10 L) in light (60 μmol photons m^−2^ s^−1^) at RT. 50 μL of β-cc was deposited on a cotton wick to favor β-cc volatilization in the airtight box. For the control treatment, β-cc was replaced by distilled water. Treatments for specific purposes are described in the corresponding subsection.

### 2.2 Luminescence quantification

Luminescence analyses were performed by treating approximately 40 12-day-old Arabidopsis seedlings in a square 12×12cm Petri dish in a sealed box (3L). 20 μL of β-cc, βI or meSA or water, as control, were deposited on a cotton wick to favor volatilization in the airtight box. The whole box was placed in a Fusion FX7 EDGE system equipped with a cooled camera (–65°C) and luminescence was acquired. Chlorophyll luminescence was acquired for 1 minute, immediately after dark, and used as a mask for successive quantifications. pAER::LUC luminescence was acquired for 5 minutes throughout the experiment. Data are reported as percentages of the value at t_0_.

For the quantification of luminescence induced by RT-EL, plates were exposed to stress conditions for 2, 3, 4 or 24 h and then transferred to the imaging system. In this case, chlorophyll luminescence was acquired for 1 minute, immediately after dark, and pAER::LUC luminescence was acquired for 5 minutes for 1 image.

### 2.3 PAM Imaging and data elaboration

Arabidopsis adult plants were dark-adapted for 20 min, and chlorophyll fluorescence was measured in a Fluorcam FC 800-O/1010 imaging fluorometer (Photon System Instruments). Chlorophyll fluorescence was acquired in a modified light curve protocol, in which plants were exposed for 1 min at each of the following actinic white-light intensities (10, 55, 150, 330, 600, 900 PPFD). F0 was measured 1 second after dark adaptation and a 6-s far-red light pulse. Far-red light has been used after every step of actinic light to obtain a measure of F0’. Fm has been measured after a 2500 µmol m^−2^ s^−1^ white light saturating pulse. PSII maximum quantum yield (Fv/Fm) was calculated as (Fm – F0)/Fm. NPQ_(T)_ has been calculated as the mean (Fm/F0-1) of control conditions divided by the (Fm’/F0’-1) of the sample at each light intensity –1 (Tietz *et al*., 2017). Y(NO) has been calculated as 1/(NPQ_(T)_’+1+qL’ x the mean (Fm/F0-1) of control conditions). qL’ has been calculated as qP’ x F0’/Ft’. Ft’ is the chlorophyll fluorescence measured at the end of the actinic period before the saturating pulse and the far-red light. qP has been calculated as Fq’/Fv’. Fq’ has been calculated as Fm’-Ft’. qE% has been calculated as the residual (1-NPQ_(T)_ after 1 min of darkness / the maximal NPQ_(T)_ of the sample) x 100.

### 2.4 GC-MS quantification of apocarotenoids

Arabidopsis adult plants were exposed to RT-EL conditions. Pools of two whole rosettes were collected and immediately frozen in liquid nitrogen. Samples were ground to a fine powder in liquid nitrogen and 200-500 mg of powder was transferred to the extraction buffer composed of: 1 mL HCl 1mM, 2 mL methyl *tert*-butyl ether (MTBE), 100 ng/ml 97% linalool (Sigma Aldrich). Samples were vortexed and sonicated for 10 minutes in ice-cold water. After centrifugation, the supernatant was transferred and evaporated under a stream of nitrogen on ice. 200 µL samples were stored for analyses.

Agilent 6890N gas chromatograph (GC) equipped with the Agilent 5973 Mass Selective Detector (MS), Column CPS CC-5MS-Xil, was used for analyses. Standard molecules were analyzed to determine retention time and confirm mass spectra. The principal degradation mass has been selected for SIM analyses (137 for β-cc, 153 for β-cca and 93 for Linalool). The analytical protocol was as follows: SIM, Flow rate 1 mL/min, starting temperature 60°C, 3°C/min until 150°C and an 8-min column cleaning at 300°C. Detection of the m/z 93 ion started at 75°C until 111°C, then detection of the m/z 137 until 126°C, then detection of the 153 m/z ion until 144°C.

The area of the peak of interest was normalized to the peak area of linalool and on the efficiency-of-detection factor, calculated as the ratio between the peak areas obtained from equal amounts of linalool and the analyte of interest. Data are expressed as nanograms of analyte per gram of fresh tissue.

### 2.5 Spectrometric quantification of anthocyanins

Arabidopsis adult plants, pretreated with β-cc or with water, were exposed to RT-EL conditions. Pools of two whole rosettes were sampled and immediately frozen in liquid nitrogen. Samples were ground to a fine powder in liquid nitrogen and 100 mg of powder was transferred to the extraction buffer (70% v/v ethanol) as described in the anthocyanin quantification method by pH differential (Lee *et al*., 2005). Anthocyanin pigment concentration was expressed as cyanidin-3-glucoside equivalents.

### 2.6 Extraction and quantification of phytohormones

To determine the hormonal profile, plant tissue was extracted with 90% (v/v) methanol containing 1% (v/v) formic acid, pre-cooled to −20 °C, at a tissue-to-solvent ratio of 1:10 (w/v). with respect to the fresh plant material. One mL of extraction solvent was added to 100 mg of each sample, followed by vortexing. The samples were subsequently treated for 15 minutes in an ice-cooled ultrasonic bath to promote efficient extraction of plant hormones. At the end of the extraction process, the samples were centrifuged at 10,000 × g for 10 minutes at 4 °C and filtered through a 0.2 µm membrane filter. Samples were stored at –80 °C until chemical analysis using an HPLC system (LC40 XR, Shimadzu, Milan, Italy) coupled with a triple quadrupole mass spectrometer (LCMS-8045, Shimadzu, Milan, Italy).

Chromatographic separation was performed on a reversed-phase C18 column (Luna C18, Phenomenex, USA; 150 × 3.0 mm, 3.0 µm), maintained at 40 °C. The mobile phase consisted of a binary system composed of 0.1% (v/v) formic acid in water (eluent A) and methanol containing 0.1% (v/v) formic acid (eluent B). The elution program was as follows: 62% (v/v) eluent A for 13 minutes; gradual increase of eluent B to 45% (v/v) between 13 and 14 minutes; decrease in eluent A to 10% (v/v) between 14 and 15 minutes, maintained until 16 minutes; return to initial conditions within 19 minutes, followed by a column rebalancing phase up to 20 minutes. The flow rate was maintained constant at 0.2 mL min^-1^. The autosampler temperature was set to 4 °C and the injection volume was 20 µL.

Hormone quantification was performed using external calibration curves obtained by dilution in 90% (v/v) MetOH at six different concentrations (1000, 100, 10, 1, 0.1, and 0.01 ng mL^-1^). The curves were constructed by plotting the areas of the chromatographic peaks as a function of concentration and applying linear regression. For each experimental treatment, the analyses were performed in technical triplicate on independent biological replicates.

### 2.7 Time-lapse *in vivo* Ca^2+^ dynamics triggered by β-cc

3-week-old *Arabidopsis thaliana* plants expressing the R-GECO1 Ca²⁺ indicator (Keinath *et al*., 2015) were imaged *in vivo* using a Nikon SMZ18 fluorescence stereomicroscope (Nikon, Japan, https://www.nikon.com/) with a SHR PLAN APO 0.5X WD:71 to image the rosette leaves. Excitation light was provided by a CoolLED pE-300 ultra illumination system at 580nm. Images were collected with a Hamamatsu C11440 CMOS camera (Hamamatsu, https://www.hamamatsu.com/). The exposure time was set to 1 s, with an 8-bit resolution of 1440×1024 pixels. Images were acquired every 10 s. For time-course experiments, fluorescence intensity was determined over regions of interest (ROIs), encompassing entire leaves. Fluorescence intensities measured within the analyzed ROIs were used for the fluorescence (F) calculation and normalized to the initial fluorescence (F0) and plotted versus time (ΔF/F0). Background subtraction was performed in each channel before fluorescence calculation by selecting a ROI outside the sample.

For imaging experiments, pots containing Arabidopsis plants were placed inside a Magenta box and covered with cellophane during microscope acquisition. at 120 seconds from the start of the experiment, 5 µL of β-cc was applied to a piece of paper placed on the soil. The Magenta box was then immediately resealed with cellophane to prevent β-cc dispersion. For control treatments, at 120 s after the start of the experiment, the cellophane cover was briefly opened and a mock application was performed by pipetting 5 µL of air to mimic the molecule administration.

### 2.8 RNA sequencing and analyses

Arabidopsis adult plants, pretreated with β-cc or with water, were exposed to RT-EL conditions. Three pools of two whole rosettes were collected and immediately frozen in liquid nitrogen. Samples were ground to a fine powder in liquid nitrogen and RNA was extracted from 100 mg of powder with TriFast (30-2010, VWR) according to the manufacturer’s instructions. RNA-seq libraries were prepared and sequenced by the DiBio sequencing service (University of Padova). Sequencing was performed in paired-end mode, targeting 20 million paired-end reads per sample. Adapter trimming and read quality filtering were performed by the sequencing service prior to downstream analyses.

Transcript abundance was quantified using Salmon against the index built from the Arabidopsis transcriptome (Araport11_cdna_20220914) (Patro *et al*., 2017). Quantification results were imported into R and summarized at the gene level for differential expression analysis accounting for transcript length and library size effects in downstream modeling (Soneson *et al*., 2016). Differential expression was tested in R using DESeq2 (Love *et al*., 2014). Gene-level count matrices were modeled with a negative binomial generalized linear model as implemented in DESeq2, using the default size-factor normalization and dispersion estimation procedures. Differential expression was assessed with Wald tests within the DESeq2 framework. P values were adjusted for multiple testing using the Benjamini–Hochberg method to control the false discovery rate (FDR). Genes were considered differentially expressed using the following criteria: minimum 10 reads, p-adj < 0.01, log2 fold changes (LFC) <-1 or >1 (Supplemental data 1). To obtain stabilized effect sizes, LFCs were shrunk using the ashr method. Shrunken LFCs were used for fold-change reporting and ranking (Stephens, 2017).

Downstream analyses and visualization were performed in R using tidyverse-compatible workflows (readr, dplyr, tidyr, purrr, fs) and base stats. Plots were generated using ggplot2, with label optimization via ggrepel (Wickham, 2016). Sample-to-sample relationships and expression patterns were visualized using heatmaps generated with pheatmap. Overlaps among gene sets were visualized using VennDiagram. Functional enrichment analyses were performed with clusterProfiler and the org.At.tair.db and AnnotationDbi packages for Arabidopsis gene annotation (Yu, 2024). In addition, enrichment and functional profiling were corroborated using gprofiler2. When needed, metadata or supplementary tables were imported using readxl. Parallel computation for selected steps was enabled via BiocParallel.

Functional enrichment was also analyzed using MapMan X4_Araport gene annotations in combination with the enricher function. In addition, the “MapMan average” algorithm was developed to directly average gene-level log_2_ fold-change values in a subclass of biological functions obtained by taking the first three levels of the BINCODE. Scripts are available in the supplementary data files.

For the rank-overlap analysis, genes were ranked by shrunken log2 fold change for the β-cc vs CTRL and RT-EL vs CTRL contrasts. At increasing rank depths (1–50% from both tails), concordant overlaps (up/up and down/down) and discordant overlaps (up/down and down/up) were tested using one-sided hypergeometric tests. Direction-specific P-values were combined using Fisher’s method and reported as –log_10_(P). The analysis was performed after restricting to genes with padj < 0.01 in at least one contrast.

Public Arabidopsis high-light RNA-seq datasets were selected from AraLightMeta (Bobrovskikh *et al*., 2025), retaining only adult leaf/rosette experiments. Public datasets were reanalyzed using a uniform differential-expression workflow, and genes with adjusted P < 0.01 were considered significant. PCA was performed on log2 fold-change profiles for genes quantified across all retained datasets and significantly regulated in at least one condition, together with the RT-EL and β-cc contrasts generated in this study.

### 2.9 Cis-regulatory motif enrichment analysis

Promoter regions spanning −1000 to +100 bp relative to the annotated transcription start site were extracted for β-cc-upregulated and background genes. Motif occurrences were identified with FIMO using P < 1 × 10^-4 (Grant *et al*., 2011) with the maximum number of stored scores increased to 1000000. JASPAR plant motifs were obtained from the JASPAR 2024 CORE collection (Rauluseviciute *et al*., 2024) and grouped into transcription factor families for visualization. Enrichment was assessed using two-sided Fisher’s exact tests with Benjamini–Hochberg correction. Motifs with FDR < 0.05 were considered significant. For strongly induced genes, genes with log2FC ≥ 1.5 were compared with the remaining β-cc-upregulated genes and motifs with FDR < 0.05 were retained.

### 2.10 Data analysis and availability

Graphs were generated and statistical analyses performed in Python (Python Software Foundation, https://python.org/) using the pandas, Matplotlib, and seaborn libraries (Hunter, 2007; McKinney, 2010; Waskom, 2021).

Statistical tests were performed using the Pingouin and SciPy libraries (Vallat, 2018; Virtanen *et al*., 2020). A Mann-Whitney post hoc test was adopted for nonparametric data comparisons and a Games-Howell for multiple comparisons of data with normal distributions.

NPQ_(T)_ images were calculated by a custom python script implementing the calculation presented in section 4.3 on the images, using the NumPy and PIL libraries (Harris *et al*., 2020).

Scripts are available in the supplementary data files.

## 3 Results

### 3.1 A buffered excess-light regime maintains stable leaf temperature

Chlorophylls and carotenoids are the main pigments responsible for light absorption in plants within the 400-700 nm range (photosynthetically active radiation) but incident irradiance does not by itself define photooxidative stress severity (McCree, 1972; Wu *et al*., 2025). Absorbed excitation can support photochemistry (biochemically active radiation), be dissipated through regulated NPQ, or contribute to non-regulated energy loss and reactive oxygen production (NO) (Zuo, 2025). These processes dissipate approximately 50% of the absorbed light even under physiological growth-light conditions (150 μmol photons m^−2^ s^−1^) and their contribution increases with light intensity (Figure 1).

**Figure 1:**
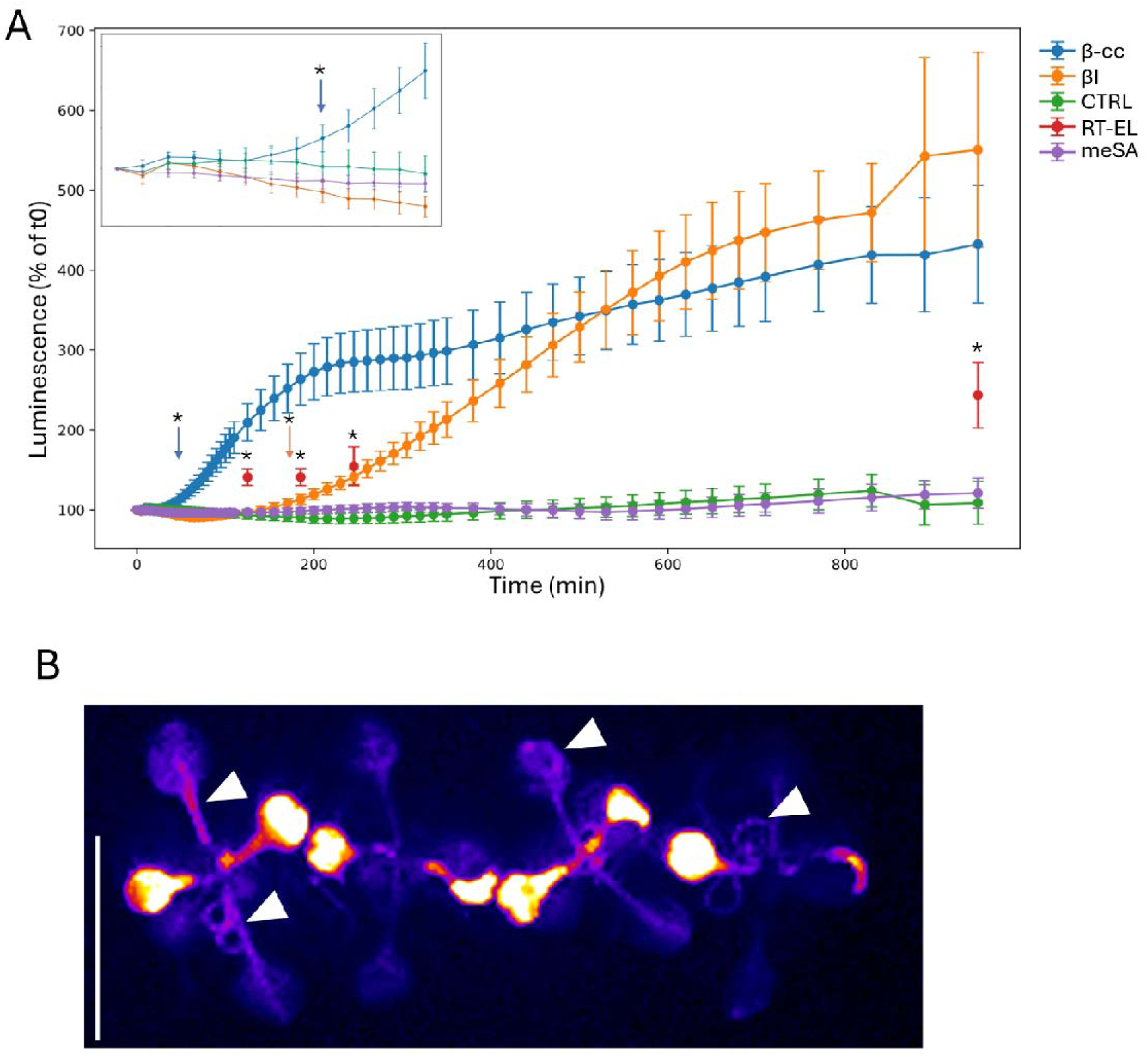
Kinetics and spatial features of β-cc signaling. (A) Time course of pAER::LUC luminescence after exposure to volatile β-cc or βI (n=120; Games-Howell pairwise tests in each timepoint); MeSA and control are shown for comparison (n=80), and RT-EL-induced luminescence is also shown (n=40) (A). The inset shows a magnification of the first 60 min (A). Representative pAER::LUC image after 16 h β-cc exposure. Reporter induction was observed in cotyledons, true leaves (particularly along the vasculature) and roots (white arrows) (B). The white bar represents 1 cm. Asterisks indicate statistically significant differences, p < 0.05. All time points following an arrow are significantly different from the control; arrow colors correspond to the respective treatments. Error bars are CI95.

Because thermal dissipation increases with irradiance, excess light would substantially elevate leaf temperature if not for evapotranspiration, provided that soil water availability allows sustained stomatal opening. Conversely, reduced soil water potential promotes stomatal closure, which limits both evaporative cooling and CO₂ uptake and may exacerbate acceptor-side limitations at PSI, thereby enhancing reactive oxygen species formation (D’Alessandro *et al*., 2020). Consistently, previous work showed that leaf temperature can increase by 4°C at 600 μmol photons m^−2^ s^−1^ or by 13°C (from 22°C to 35°C) at 1200 μmol photons m^−2^ s^−1^, when air cooling is insufficient (Balfagón *et al*., 2019).

We exposed four-week-old adult plants to 1500 μmol photons m⁻² s⁻¹ at an air temperature of approximately 26 °C. Plants remained well watered and received gentle airflow. Before excess-light exposure, plants were treated for 4 h with volatile β-cc (300 μmol in a 10-L chamber) or water. Infrared imaging showed that leaves remained within 23–27 °C throughout the treatment, with no detectable difference between β-cc-pretreated and control plants (Figure S1). We therefore refer to this regime as room-temperature excess light (RT-EL).

Leaf temperature did not differ between β-cc–treated and control plants, indicating no major β-cc effect on evapotranspiration under these well-watered conditions.

**Figure S1:**
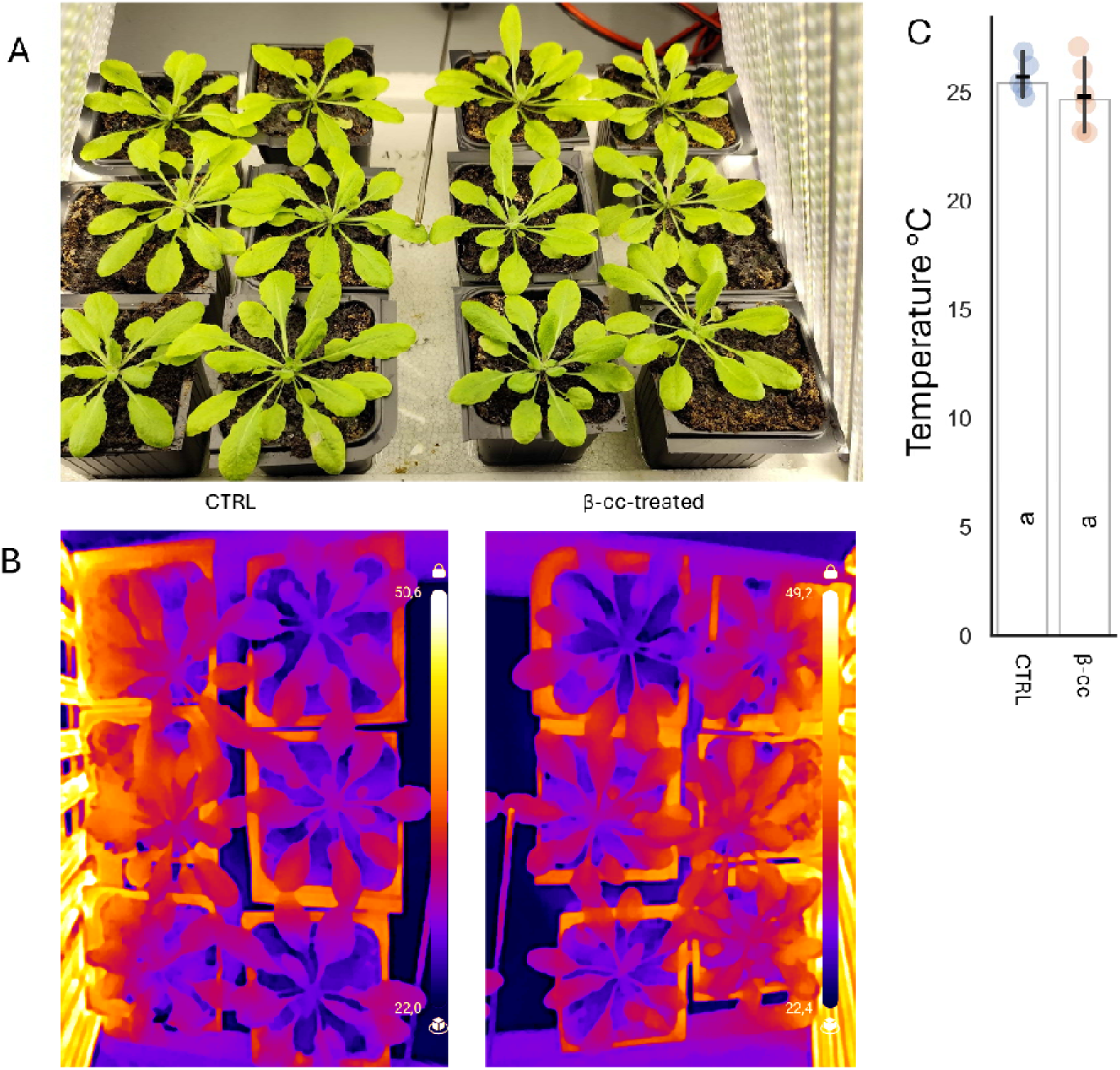
Visible-light (A) and infrared (B) images of 4-week-old adult Arabidopsis plants exposed to 1500 μmol photons m^-2^ s^-1^. Temperature scale from black to white (22°C to 50°C). Leaf temperature distributions were quantified across plants and showed similar mean temperatures in control (CTRL) and β-cc–treated plants (n=6; t-test), remaining within 23–27 °C throughout the treatment (C). Letters represent statistically homogeneous groups.

### 3.2 Exogenous β-cyclocitral elicits rapid responses

Four-hour β-cc treatments have been widely used to characterize β-cc-dependent transcriptional and physiological responses in terrestrial plants (Lv *et al*., 2015; D’Alessandro *et al*., 2018; Tiwari *et al*., 2025). However, this commonly used endpoint provides little information on how rapidly β-cc is perceived. We therefore asked whether β-cc signaling is initiated substantially earlier than the established 4-h response.

To resolve the timing of β-cc signaling, we used a luciferase reporter line (pAER::LUC) (Rac *et al*., 2021). Volatile β-cc (100 µmol in a 3-L headspace) was applied to 12-day-old seedlings in a sealed transparent chamber, and luminescence was monitored over time. *AER* (*At5G16970*) encodes an alkenal reductase, a component of the SCL14-dependent detoxification program, which is robustly but not specifically induced by β-cc (D’Alessandro *et al*., 2018). Although reporter luminescence occurs downstream of a complex transcriptional activation (SCL14-TGAII and ANAC102-ANAC081/002/032), pAER::LUC-dependent luminescence increased within 40 min after β-cc exposure (Figure 1A, blue arrow), indicating rapid activation of the β-cc response. Spatially, reporter induction was strongest in cotyledons, but was also evident in true leaves (particularly along the vasculature) and, unexpectedly, in roots (Figure 1B).

We next compared β-cc to β-ionone (βI; 100 µmol in a 3-L headspace), another β-carotene–derived apocarotenoid that does not increase plant tolerance to excessive light, while it has recently been reported to improve plant responses to biotic stress (Ramel *et al*., 2012; Felemban *et al*., 2024). β-cc induced pAER::LUC earlier than βI, whereas βI elicited a comparable response only at later time points (Figure 1A, orange arrow at 155 min). Because βI can oxidize to dihydroactinidiolide (dha), which shares activity with β-cc (Shumbe *et al*., 2014), the delayed induction of the pAER::LUC signal by βI may reflect the conversion of βI to dha and/or a lower specificity for the receptor.

Methyl salicylate (MeSA; 150 µmol in a 3-L headspace) was included because SCL14-dependent detoxification can be responsive to salicylic acid–related signaling (Fode *et al*., 2008). Under our assay conditions, MeSA did not induce pAER::LUC luminescence, suggesting that AER induction is not a general response to volatile defense-related cues in this setup (Figure 1A, purple line).

Finally, we also analyzed the induction of the luciferase reporter under RT-EL conditions: plates containing around 40 seedlings were exposed to the stress conditions and analyzed after 2, 3, 4 and 24 h of treatment. The induction of the pAER::LUC reporter after only 2 h of treatment, may indicate an early production of β-cc, readily metabolized by cellular mechanisms, or that the SCL14-dependent detoxification pathway could be activated in a β-cc-independent manner, likely due to other reactive species, which are known to induce the pathway (Figure 1A, red points).

Because β-cc triggered the transcriptional reporter activation on a sub-hour timescale, we next investigated whether it also elicited rapid cellular signals such as calcium signaling. Cytosolic calcium concentration ([Ca^2+^]_cyt_) dynamics were monitored in adult Arabidopsis leaves expressing the fluorescent R-GECO1 indicator (Figure 2A-E) (Keinath *et al*., 2015). Exposure to volatile β-cc (25 μmol volatile in 0.5 L) induced a rapid increase in [Ca^2+^]_cyt_ that became detectable within the first few minutes after treatment. The signal increased progressively and approached a plateau after approximately 20 min (Figure 2F). Notably, comparable calcium responses were observed both in leaves proximal to the β-cc application site and in distant leaves, whereas control-treated plants showed only minimal fluctuations in fluorescence. Consistently, the maximal ΔF_max_/F₀ response was significantly higher in both proximal and distant β-cc-treated leaves than in the corresponding controls, with no significant difference between proximal and distant leaves (Figure 2G). These results indicate that volatile β-cc triggers a rapid and spatially distributed [Ca^2+^]_cyt_ response that is not restricted to tissues located near the site of application.

**Figure 2:**
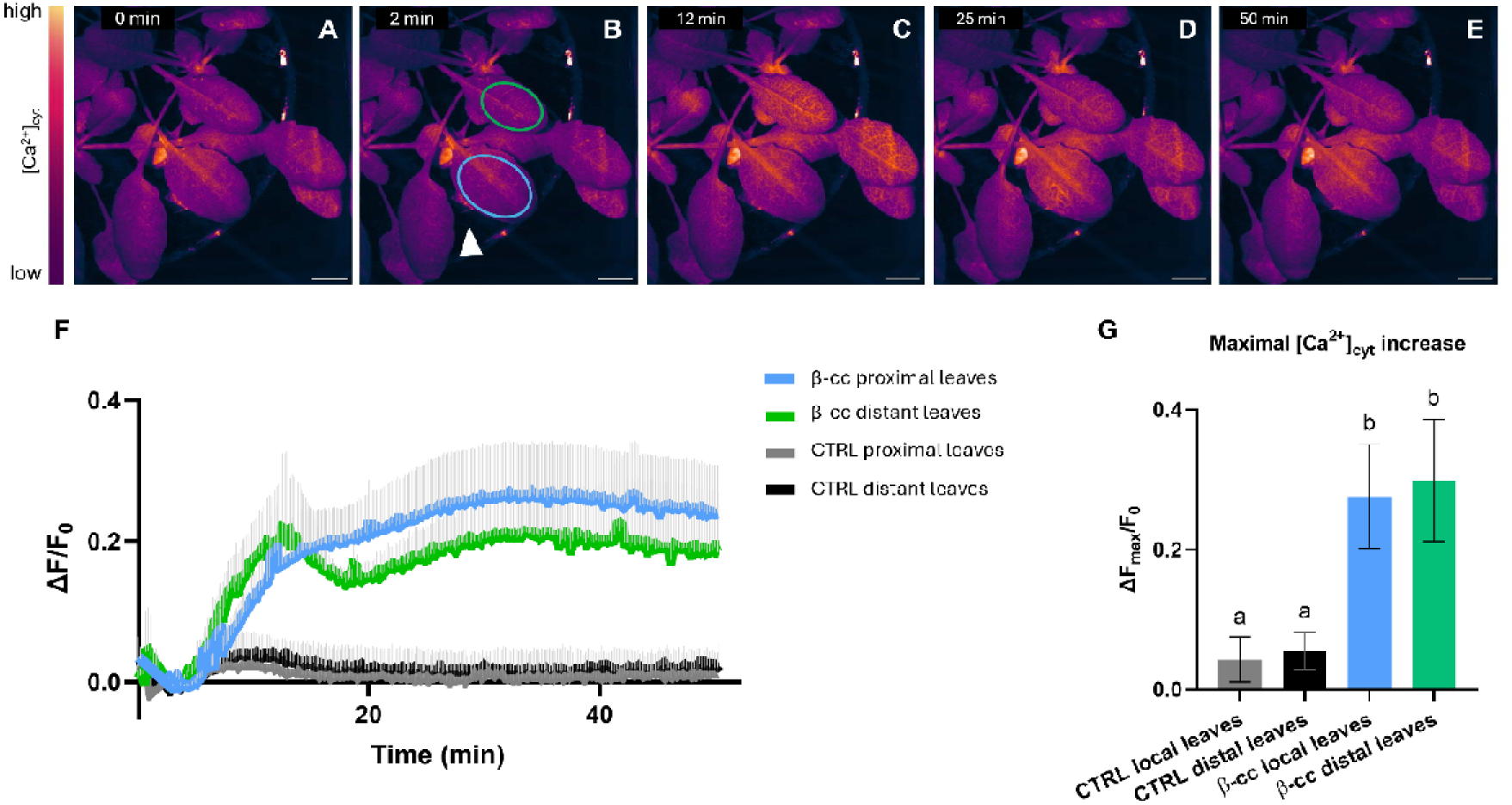
[Ca^2+^]_cyt_ transients induced by β-cc. (A–E) False-color time-lapse imaging of β-cc administration to plants expressing the cytosolic Ca²⁺ indicator R-GECO1. (F) β-cc was applied at 2 min. (B) The white arrow indicates the site of β-cc application on a piece of paper on the soil. The light blue circle marks the proximal leaf, defined as the analyzed Region of Interest (ROI) on the leaf near the β-cc application site. The green circle marks the distant leaf, defined as the ROI located on a plant distant from the application site. (F) Average traces showing cytosolic Ca²⁺ increases triggered by β-cc or control treatments. (G) Maximal cytosolic Ca²⁺ increase induced by β-cc or control treatments. Scale bar = 5000 µm. n ≥ 5 leaves analyzed per experiment. One-way ANOVA, alpha = 0,05. Letters represent statistically homogeneous groups. Error bars are SD.

### 3.3 Photosynthetic adjustments of β-cc-treated plants under RT-EL

We assessed the photosynthetic performance of β-cc-pretreated and control plants by pulse-amplitude-modulated (PAM) imaging under control conditions and during exposure to RT-EL. Under control conditions, volatile β-cc pretreatment did not affect the maximum quantum yield of PSII (Fv/Fm) or NPQ_(T)_ measured at growth irradiance (150 μmol photons m^-2^ s^-1^) or high irradiance (900 μmol photons m^-2^ s^-1^) (Figure 3 A, B). This contrasts with previous observations in aquatic photosynthetic organisms, where β-cc can impair PSII efficiency and NPQ (Tietz *et al*., 2017; Mitra *et al*., 2021; Roach *et al*., 2022). NPQ_(T)_ was calculated as NPQ normalized to the mean F_m_/F_0_ of control samples, and therefore includes sustained and photoinhibitory quenching (Tietz *et al*., 2017).

**Figure 3:**
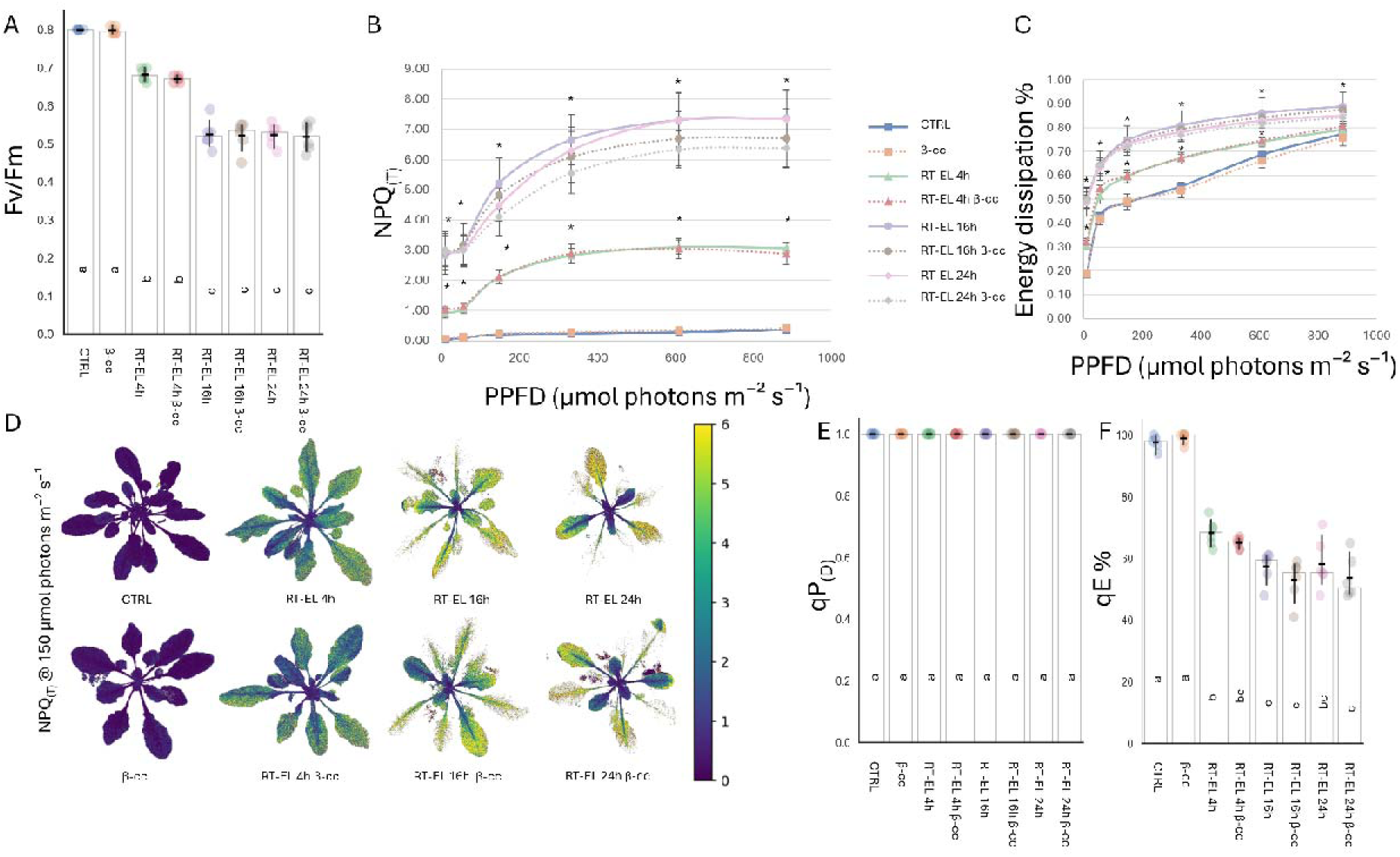
Photosynthetic responses of Arabidopsis to room-temperature excess light (RT-EL) with or without β-cyclocitral (β-cc) pretreatment. (A) Maximum quantum yield of PSII (Fv/Fm). (B) NPQ_(T)_ light response curves, expressed as NPQ normalized to the mean Fm/F0 of control plants. (C) Fraction of light dissipated under increasing actinic light irradiance. (D) Representative false-color images of NPQ measured at growth irradiance (150 μmol photons m^-2^ s^-1^). (E) qP_(D)_, measured after 1 min of darkness, used as an estimate of rapid dark recovery of the PSII acceptor side. (F) qE%, calculated from NPQ relaxation after 1 min of darkness and representing the rapidly reversible fraction of NPQ. Data are mean ± CI95 (n = 6). Statistical analysis: Mann-Whitney pairwise tests at each light intensity, alpha = 0,05. Letters represent statistically homogeneous groups. Error bars indicate 95% CI in panels A, E and F and SDs in panels B and C. RT-EL (1500 μmol photons m^-2^ s^-1^).

Exposure to RT-EL strongly altered all photosynthetic parameters (Figure 3). Fv/Fm progressively decreased from 0.8 under control conditions to 0.68 after 4 h of RT-EL, and reached 0.52 after 16 h, remaining at a similar level at 24 h (Figure 3A). Thus, most of the decrease in PSII maximal efficiency occurred within the first 16 h of treatment, after which values appeared to stabilize. In parallel, NPQ_(T)_ increased markedly and approached a plateau by 16 h, when assessed at both growth and high irradiance (Figure 3B). At growth irradiance, NPQ_(T)_ increased from 0.2 under control conditions to 2 after 4 h and approximately 5 after 16 h of RT-EL. While, at high irradiance, it increased from 0.4 to 3 after 4 h and to 7 after 16 h (Figure 3B). The stabilization of both Fv/Fm and NPQ_(T)_ after 16 h suggests that plants were able to enter a new photosynthetic steady state under RT-EL, characterized by sustained dissipation of excess excitation energy and partial maintenance of PSII functionality (Figure 3C). Indeed, Fv/Fm values under RT-EL remained higher than those previously reported for excess-light treatments, consistent with a milder degree of photoinhibition limiting the development of severe photooxidative damage (Lv *et al*., 2015; Shumbe *et al*., 2017; D’Alessandro *et al*., 2018).

The increase in quenching under RT-EL was also evident spatially. Representative NPQ_(T)_ images acquired at growth irradiance showed a clear increase in dissipative activity in stressed plants relative to controls, with the strongest signal generally observed in older leaves, whereas the shoot apex and younger leaves appeared less affected (Figure 3D). This spatial heterogeneity is consistent with leaf age-dependent differences in sensitivity to prolonged excess light.

RT-EL did not affect the rapid reopening of PSII reaction centers in the dark (qP_(D)_ after 1 minute of darkness), a measure of the PSII status at the acceptor side (Figure 3E). qP(D) remained close to 1 under all conditions, indicating rapid re-oxidation of Q_A_ after transfer to darkness and suggesting that the photosynthetic electron transport chain retained substantial functionality. We also estimated the rapidly reversible fraction of NPQ after 1 min of darkness (qE%). In control plants, qE% was high, indicating that most quenching relaxed rapidly in the dark (Figure 3F). By contrast, RT-EL reduced qE%, after 4 h, showing that a progressively larger fraction of total quenching was due to a sustained quenching component. The sustained quenching (qS) was also appreciable at low irradiance (NPQ_(T)_ at 10 μmol photons m^-2^ s^-1^), as NPQ increased from 0.05 under control conditions to 1 after 4 h of RT-EL, and then to 3 at 16 and 24 h, consistent with an increasing contribution of photoinhibitory components of NPQ under prolonged stress.

Overall, these data indicate that RT-EL induced an acclimated photosynthetic state in adult Arabidopsis plants, which limited the build-up of uncontrolled excitation pressure. Importantly, none of these RT-EL–induced changes differed between β-cc–pretreated and control plants (Figure 3A–F), indicating that β-cc signaling did not modify the photosynthetic response to mild stress conditions. Thus, although β-cc signaling occurs on a timescale comparable to the onset of photosynthetic adjustment, it does not appear to contribute substantially to the establishment of this acclimated state.

### 3.4 Endogenous β-cyclocitral accumulates after prolonged RT-EL

In Arabidopsis, β-cc is primarily generated through oxidative degradation of β-carotene associated with photosystem II (D’Alessandro and Havaux, 2019; Carnà *et al*., 2025). This process happens in the presence of overexcitation even though only a minimal fraction of β-carotene is chemically degraded during the quenching. Low-level oxidative turnover of β-carotene is expected to occur even under non-stress conditions, consistent with the detectable basal β-cc pool. We therefore asked when sustained RT-EL caused β-carotene oxidation products to accumulate above this basal level.

Endogenous β-cc remained near the control level at 4 h and increased only after prolonged RT-EL exposure (16 to 48 h of stress) (Figure 4A). β-cc levels increased from 5 ng/g fresh weight (FW) under control conditions to 25, 55 and 95 ng/g(FW) after 16, 24 and 48 h of RT-EL, respectively. These amounts are considerably lower than values commonly reported for excess-light treatments in Arabidopsis, which often reach 250 ng/g FW, with higher baseline levels as well. The discrepancy may reflect differences in growth and irradiation regimes and is consistent with lower β-cc amounts reported in other studies (Taniguchi *et al*., 2023).

**Figure 4:**
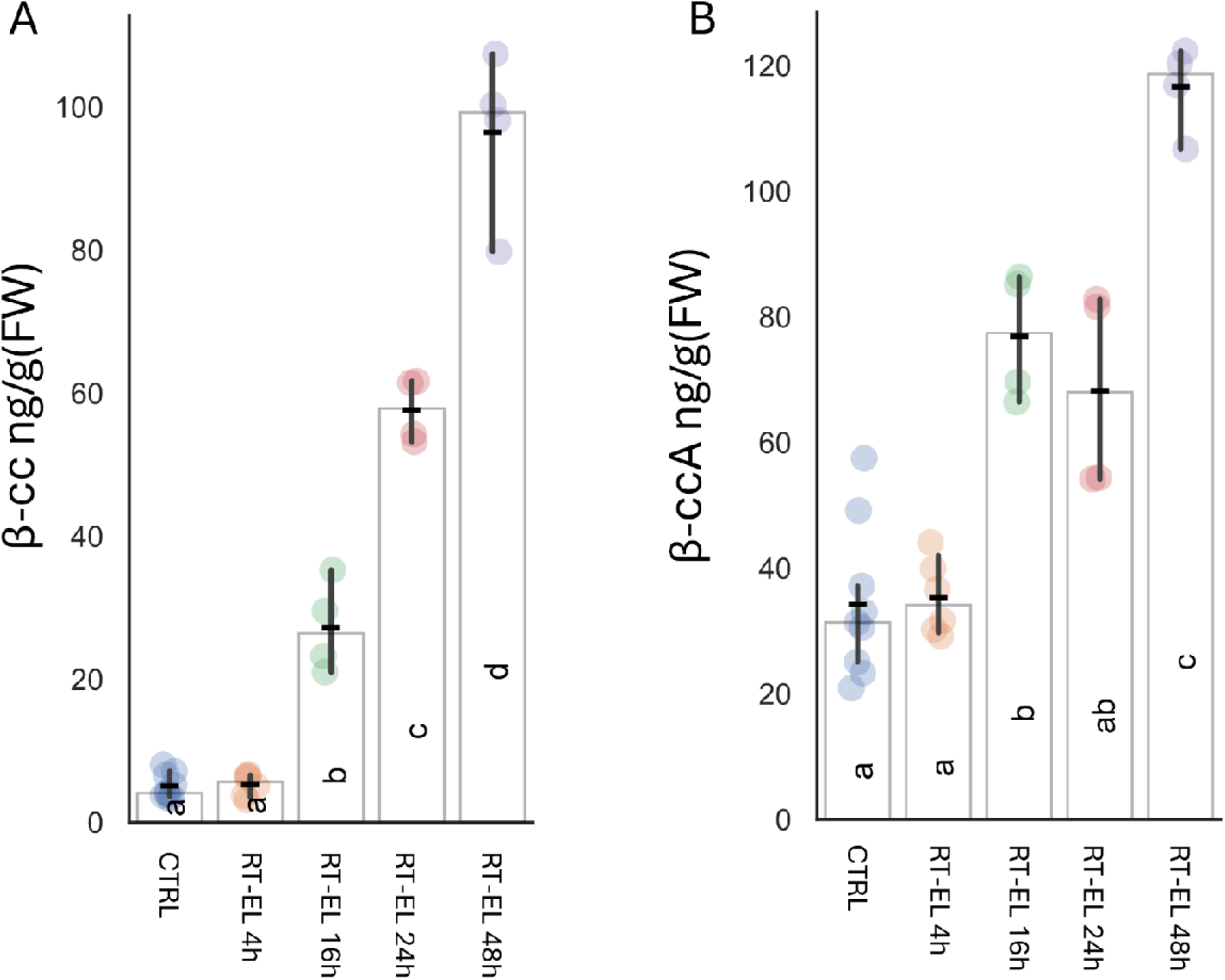
Endogenous β-cyclocitral (β-cc) and β-cyclocitric acid (β-cca) accumulation during RT-EL. (A) β-cc and (B) β-cca quantified by GC–MS in control plants and during RT-EL. n = 8 (4 biological replicates × 2 technical replicates); statistics: Games–Howell test, α = 0.05. Letters represent statistically homogeneous groups. Error bars are 95% CI. RT-EL (1500 μmol photons m^-2^ s^-1^).

We also quantified β-cyclocitric acid (β-cca), the main product of β-cc oxidation (D’Alessandro *et al*., 2019). β-cca increased from 30 ng/g (FW) under control conditions, to 80 ng/g(FW) after 16 h and 120 ng/g(FW) after 48 h of RT-EL (Figure 4B). Notably, the β-cca:β-cc ratio shifted from 6:1 in plants grown under control conditions to approximately 1:1 in plants subjected to 48 h RT-EL, suggesting that RT-EL alters the balance between β-cc formation and β-cca conversion and/or turnover.

Although β-cc is often considered a rapid marker of photosystem overexcitation (D’Alessandro and Havaux, 2019; D’Alessandro *et al*., 2020), neither β-cc nor β-cca showed a significant increase after 4 h RT-EL (Figure 4A,B). The absence of a detectable increase at 4 h does not necessarily imply that β-cc formation was absent, because the measured pool reflects the balance between production, volatilization, oxidation to β-cca, and further metabolism. Nevertheless, it demonstrates that β-carotene oxidation products did not accumulate above their basal steady-state levels during the photosynthetic adjustment phase. Notably, previous studies showed that β-cc was already 50% higher after 1 h of stress at controlled air-temperature (7°C) (Ramel *et al*., 2012). However, the β-cc treatment caused a strong accumulation in the plants, similar to what observed previously (D’Alessandro *et al*., 2019).

### 3.5 β-cyclocitral crosstalk with phytohormones

To characterize β-cc effects, we quantified anthocyanin accumulation and plant hormones, which have been linked to its signaling in literature. RT-EL induced anthocyanin accumulation from 16 h onward and the β-cc pretreatment did not measurably alter RT-EL–induced anthocyanin levels, in contrast to prior reports under low-temperature excess-light regimes (Figure 5A) (Lv *et al*., 2015).

**Figure 5:**
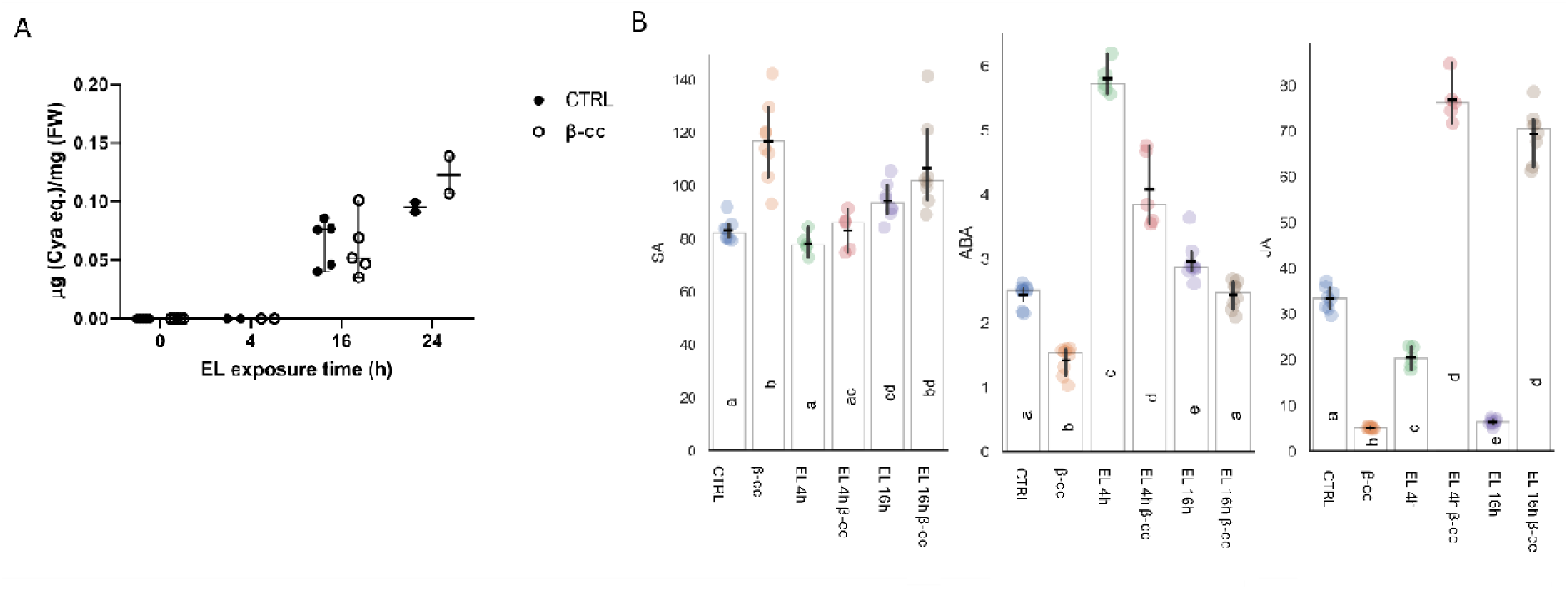
Physiological and hormonal responses to β-cc and RT-EL. Anthocyanin content (n=5, two-way ANOVA) (A). Salicylic acid (SA), abscisic acid (ABA) and jasmonic acid (JA) levels (ng/g (FW)) in control plants, β-cc–treated plants under normal light, and plants exposed to RT-EL with or without β-cc pretreatment (n=6, Mann-Whitney pairwise for ABA, Games-Howell pairwise for JA and SA) (B). Letters represent statistically homogeneous groups. Error bars represent 95% CI.

We next measured SA, ABA, and JA (Figure 5B). Under control conditions, β-cc modestly increased SA (30%) and reduced ABA (40%), consistent with earlier observations for ABA regulation by β-cc (Taniguchi *et al*., 2023). Under RT-EL, SA did not show a pronounced additional increase in β-cc–pretreated plants relative to RT-EL controls, consistent with the absence of a strong SA burst, supporting an acclimated state of the plants under these conditions rather than activation of PCD (Lv *et al*., 2015; D’Alessandro *et al*., 2020). ABA increased transiently after 4 h of RT-EL (approximately twofold) and returned toward baseline levels by 16 h and β-cc partially attenuated ABA accumulation at 4 h (Figure 5B).

JA displayed the strongest modulation. Under control conditions, β-cc reduced JA to 15% of control levels, whereas under RT-EL β-cc enhanced JA accumulation relative to RT-EL alone (Figure 5B). Together, these data indicate that β-cc can substantially reshape stress-hormone homeostasis, with effects that depend on the physiological context (control vs RT-EL).

### 3.6 β-cyclocitral signaling under RT-EL

To characterize β-cc effects and their interaction with the RT-EL response, we performed RNA-sequencing on adult Arabidopsis plants after β-cc pretreatment and 16 h of RT-EL. This time point coincided with the plateau in the major fluorescence changes and the first clear increase in endogenous β-cc.

Principal component analysis (PCA) separated β-cc-treated samples from controls along PC1, which explained 79% of the variance, while both RT-EL groups occupied the opposite side of the same axis (Figure 6A). Consistent with previous reports, β-cc alone triggered extensive transcriptional reprogramming, with approximately 4,000 differentially expressed genes (DEGs) relative to controls (Figure 6B) (Del Boccio *et al*., 2025; Tiwari *et al*., 2025). RT-EL also elicited a comparably large response, with ∼4,000 DEGs relative to controls (Figure 6C).

**Figure 6:**
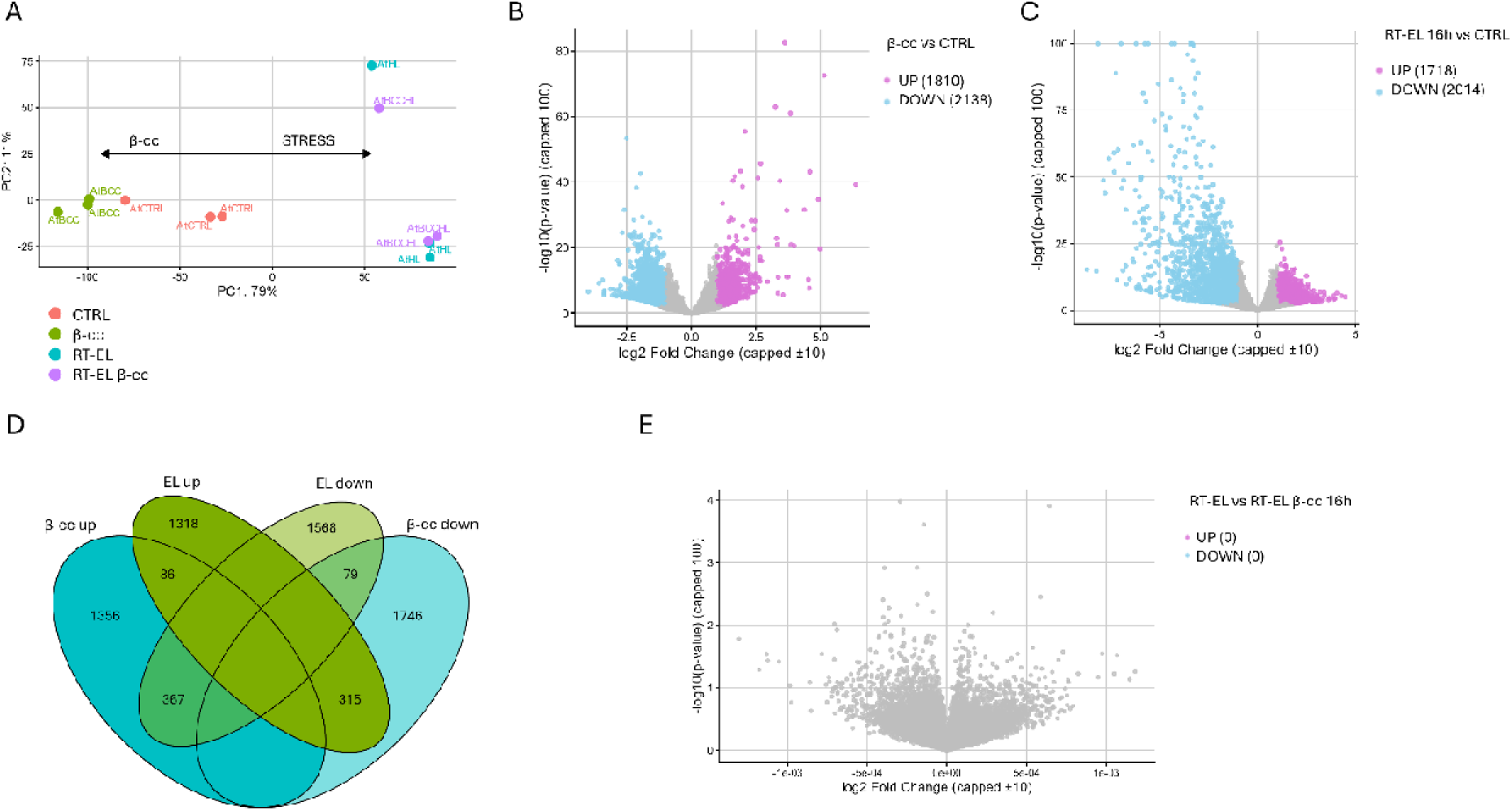
Transcriptomic responses to β-cc and RT-EL in adult Arabidopsis. PCA plot based on the top 4000 variable genes (A). Volcano plot of β-cc-treated samples versus control samples (B). Volcano plot of samples exposed to excessive light for 16 h versus control samples (C). Venn diagram comparing DEGs from β-cc vs control and RT-EL vs control, separated by direction of regulation (D). Volcano Plot of samples exposed to excessive light for 16 h treated with β-cc versus untreated samples (E). Data were obtained from RNA-seq analyses on n = 3 biological replicates.

The β-cc and RT-EL programs showed limited concordance. Only 166 DEGs were regulated in the same direction in both comparisons (87 upregulated and 79 downregulated) (Figure 6D). In contrast, a larger set of genes displayed opposing regulation: 367 genes were upregulated by β-cc but downregulated by RT-EL, and 313 genes were downregulated by β-cc but upregulated by RT-EL (Figure 6D). These results indicate that β-cc and RT-EL induce largely distinct and frequently contrasting, transcriptional responses. Importantly, the direct comparison within RT-EL (RT-EL + β-cc vs RT-EL) revealed the absence of a residual β-cc signature (Figure 6E), indicating that RT-EL largely overrides β-cc-dependent transcriptional changes under these conditions, in agreement with the absence of β-cc-mediated physiological priming. To further characterize the interaction between β-cc and RT-EL, we tested the interaction between the two stimuli (β-cc x RT-EL). The interaction term identified almost all the β-cc responsive genes as significantly affected by the interaction (Figure S2A), indicating that the transcriptional output of the combined treatment cannot be explained simply by the sum of the responses to β-cc and RT-EL applied individually. This widespread interaction confirms the observation that the β-cc transcriptional signature is largely suppressed or reshaped under RT-EL conditions.

We also performed a rank-overlap analysis using genome-wide rankings based on the expression changes for the β-cc versus CTRL and RT-EL versus CTRL contrasts. Significantly altered genes (P-adj value < 0.01) in at least one condition were progressively sampled from both extremes of the ranked distributions. Concordant overlaps remained limited and confined to the most strongly regulated genes, corresponding mainly to the SCL14-dependent detoxification response. By contrast, discordant overlaps, corresponding to genes regulated in opposite directions by β-cc and RT-EL, showed a marked and progressively increasing enrichment (Figure S2B). These results demonstrate that the antagonistic relationship between the β-cc– and RT-EL-induced transcriptional programs represents a broad genome-wide feature of the two responses.

**Figure S2:**
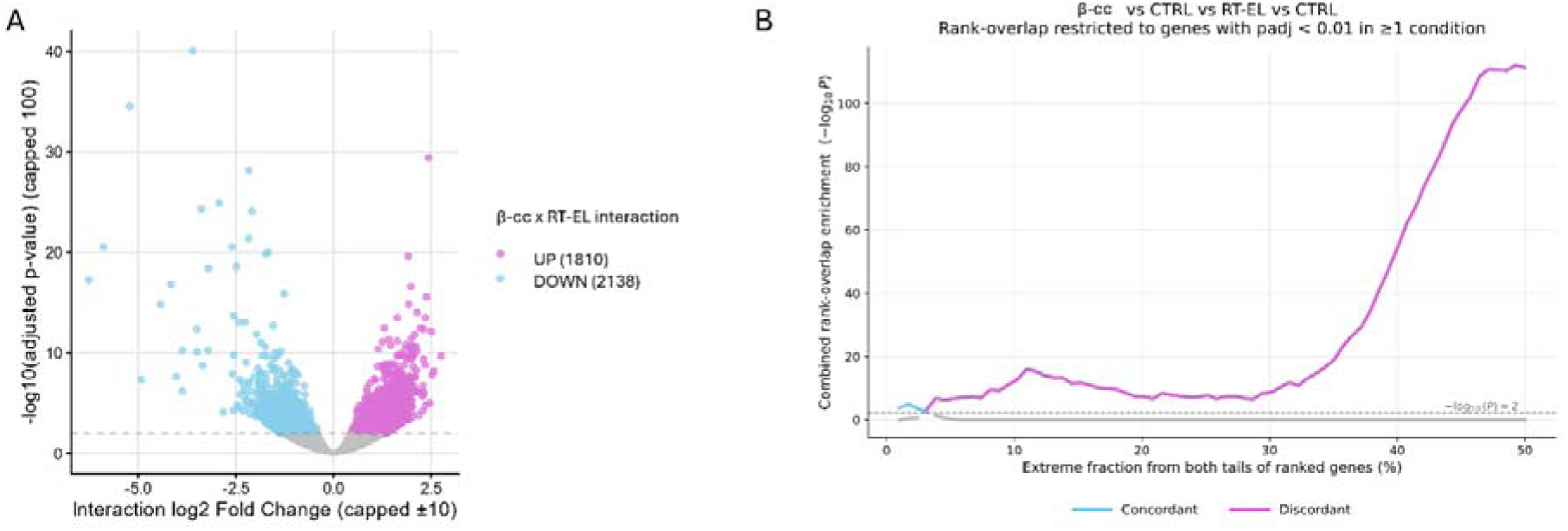
Interaction and rank-overlap analysis of β-cc and RT-EL transcriptional responses. Volcano plot of the β-cc x RT-EL interaction, showing genes with significant positive and negative interaction effects (A). Rank-overlap analysis comparing β-cc and RT-EL responses, restricted to genes with padj < 0.01 in at least one condition; concordant and discordant overlaps are shown across increasing fractions of the ranked gene lists (B).

We then investigated whether specific transcription-factor families were preferentially associated with the β-cc response. Motif-family enrichment analysis revealed several enrichments depended both on the direction and magnitude of gene regulation (Figure 7). Among β-cc-upregulated genes, enrichment became particularly evident at higher induction thresholds for several transcription-factor families. TGA-, bZIP/ABRE-, and bHLH/MYC/PIF-related motifs showed the most consistent enrichment represented by multiple significantly enriched motifs, particularly among strongly β-cc-induced genes. WRKY motifs were also enriched among the most strongly induced genes, whereas RVE/CCA1/LHY, TCP, ARF/Auxin, AP2/ERF and TALE families were detected among moderately induced genes. A different pattern was observed among β-cc-downregulated genes. MYB-related motifs were the most prominent, showing significant enrichment among the most strongly repressed genes, as well as a strong depletion from induced genes. RVE/CCA1/LHY and WRKY showed odds ratios below one in the downregulated gene set, indicating relative depletion rather than enrichment, in opposition to the enrichment in the induced genes. LBD and E2F/DP motifs were enriched among genes with more moderate negative fold changes. In contrast, CAMTA was depleted.

**Figure 7:**
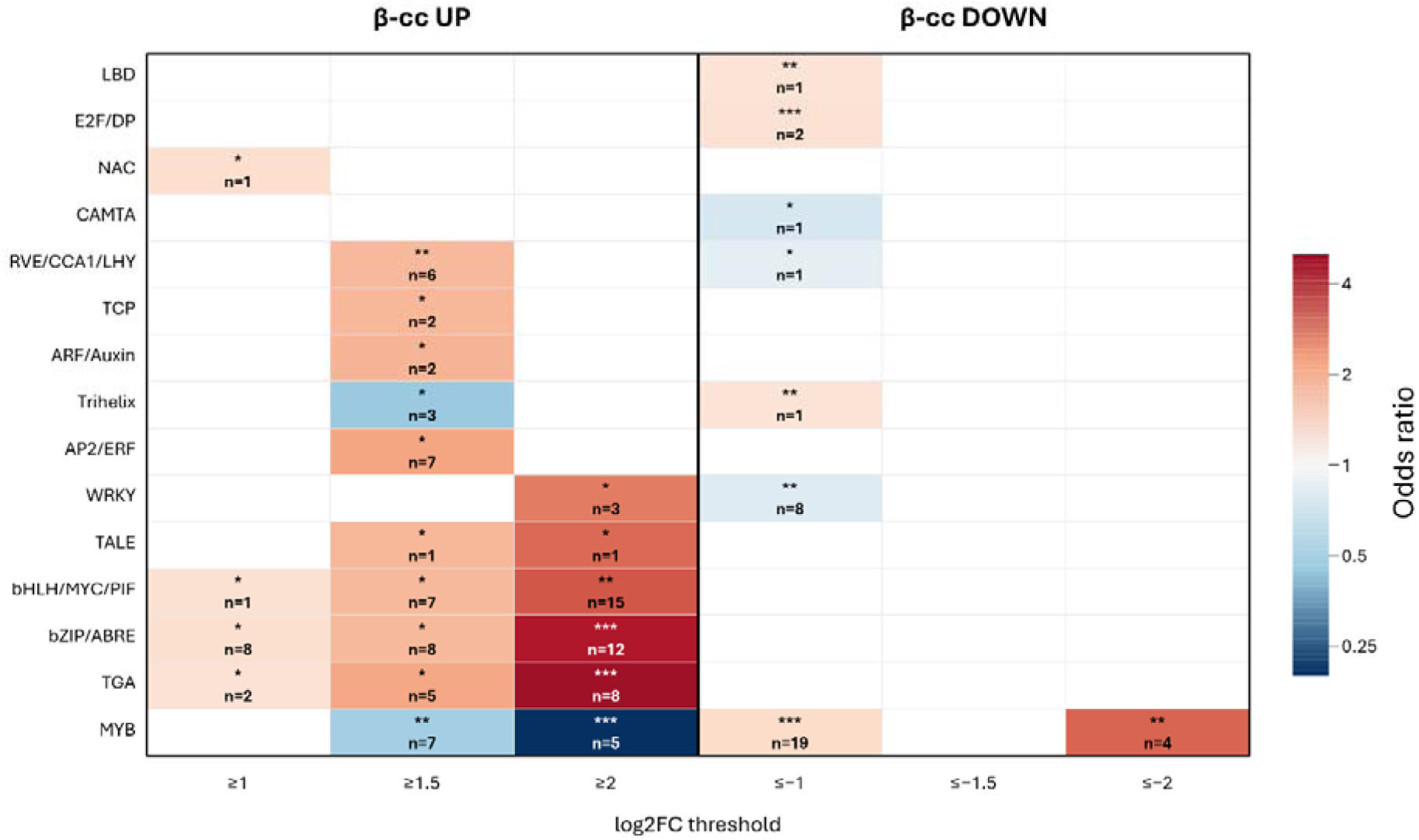
Transcription-factor-family motif enrichment across the β-cc response. Motif enrichment was summarized separately for genes upregulated or downregulated by β-cc across log2 fold-change thresholds. Tile color indicates the odds ratio of family representative enrichment, asterisks indicate significance, and n reports the number of enriched representatives (*p<0.01, **p<0.001, ***p<0.0001).

Overall, these data suggest that β-cc-dependent transcriptional reprogramming is associated with a defined set of transcription-factor families, with particularly strong representation of MYB-, TGA-, bZIP/ABRE– and bHLH/MYC/PIF– and WRKY-related regulatory signatures. This pattern is consistent with the broad β-cc-dependent transcriptional reprogramming, supports the established involvement of TGA transcription factors in β-cc signaling, and identifies additional candidate regulatory factors that may contribute to this response.

Subcellular-location enrichment further distinguished the responses. Using SUBA annotations, we tested whether predicted subcellular localizations were overrepresented among up– or downregulated genes (Hooper *et al*., 2017). β-cc-regulated genes were enriched for proteins assigned to the vacuole, cytosol, and secretory pathway, with depletion of nuclear elements. RT-EL preferentially affected genes encoding mitochondrial and plastid proteins (Supplemental Data 1).

Gene Ontology, KEGG, MapMan, and gene-set enrichment analyses identified the same broad separation (Ashburner *et al*., 2000; Thimm *et al*., 2004; Subramanian *et al*., 2005; Kanehisa *et al*., 2010). β-cc positively enriched mitochondrial electron transport and oxidative phosphorylation and affected the ICE-CBF cold-response module, while negatively enriching several categories related to nuclear regulation, RNA metabolism, and defense (Table 1). RT-EL positively enriched RNA processing, protein folding, proteostasis, and mitochondrial import, and negatively enriched light-harvesting and hormone-signaling categories (Table 2). Complete ranked outputs are provided in Supplemental Data 1.

**Table 1:** selected functional enrichments induced by β-cc.

| GO code | GO term | Score | Gene count | Algorithm |
| --- | --- | --- | --- | --- |
| GO:0019646 | aerobic electron transport chain | 7.8 | 26 | enrichGO |
| GO:0010629 | negative regulation of gene expression | -12.5 | 165 | enrichGO |
| GO:0140719 | constitutive heterochromatin formation | -7.3 | 31 | enrichGO |
| ath00190 | Oxidative phosphorylation | 6.6 | 54 | enrichKEGG |
| ath03013 | Nucleocytoplasmic transport | -9.3 | 59 | enrichKEGG |
| ath03018 | RNA degradation | -6.5 | 50 | enrichKEGG |
| GO:0042775 | mitochondrial ATP synthesis coupled electron transport | 3.8 | 19 | gprofiler2 |
| GO:0051028 | mRNA transport | -3.4 | 34 | gprofiler2 |
| 15.7.31 | PHD finger transcription factor | -6.9 | 12 | Enricher:MapMan |
| 26.6.2.1 | NLR effector receptor | -6.3 | 35 | Enricher:MapMan |
| 15.7.30 | JUMONJI transcription factor | -5.6 | 12 | Enricher:MapMan |
| 7.4.6 | coenzyme A kinase | 1.58 |  | MapMan average |
| 26.3.4 | ICE-CBF cold acclimation transcriptional cascade | 1.21 |  | MapMan average |

**Table 2:** selected functional enrichments induced by RT-EL.

| GO code | GO term | Score | Gene count | Algorithm |
| --- | --- | --- | --- | --- |
| GO:0009451 | RNA modification | 12.5 | 138 | enrichGO |
| GO:0006457 | protein folding | 10.9 | 82 | enrichGO |
| GO:0170036 | import into the mitochondrion | 4.3 | 17 | enrichGO |
| GO:0050821 | protein stabilization | 4.3 | 13 | enrichGO |
| GO:0009768 | photosynthesis, light harvesting in photosystem I | -7.4 | 16 | enrichGO |
| GO:0009733 | response to auxin | -7.1 | 100 | enrichGO |
| ath04075 | Plant hormone signal transduction | -8.3 | 114 | enrichKEGG |
| ath00196 | Photosynthesis - antenna proteins | -6.5 | 14 | enrichKEGG |
| GO:0009538 | photosystem I reaction center | -4.2 | 8 | gprofiler2 |
| 15.7.33 | bHLH transcription factor | -4 | 39 | Enricher:MapMan |
| 18.8.1.30 | MAP3K-RAF kinase | -3.6 | 16 | Enricher:MapMan |
| 15.7.32 | TAZ transcription factors | -2 |  | MapMan average |
| 27.2.2 | MRF TOR-dependent protein translation regulators | -1.9 |  | MapMan average |

MapMan pathway views clarified which components drove the enrichments. β-cc and RT-EL both induced parts of mitochondrial oxidative phosphorylation, but alternative oxidase genes responded mainly to RT-EL (Figures S3A and S4A). β-cc modestly increased several light-reaction transcripts, whereas RT-EL broadly repressed the same modules (Figures S3B and S4B). Both treatments affected JA biosynthesis, with a remarkable difference at the level of lipoxygenase (LOX) genes, which were downregulated by β-cc (Figures S3C, S4C). This regulation may account for the reduced JA levels in β-cc-treated plants under control conditions (Figure 5B).

Analysis of selected marker genes further emphasized the contrast between treatments (Supplemental Data 1). β-cc increased the expression of the transcription factor *GLK2*, while decreasing *GUN1*, whereas RT-EL strongly repressed several *LHCII* antenna genes and *GLK2*. β-cc induced the oxidative-stress-responsive genes *ZAT12* and *FER1*, reporters of H_2_O_2_, while RT-EL also induced *APX1* and *BBX1*, and increased expression of the kinase *AGC2* OXI1, consistent with activation of oxidative stress signaling often linked to lesion– and cell-death associated pathways (Beaugelin *et al*., 2019; D’Alessandro *et al*., 2020). Notably, β-cc induced 38 singlet oxygen marker genes (eSORG), under growth irradiance, different from the 34 induced by RT-EL, highlighting the differences between the two stimuli (Dogra *et al*., 2017).

The strongest shared component was SCL14-dependent detoxification. CYP81D11, AER, GRX480, GSTU7/19/24, and UGT73B2/B3 were induced by β-cc and contributed to the 87 commonly upregulated genes induced by RT-EL. Among these, AER expression increased sixfold with β-cc and twofold with RT-EL, consistent with the kinetics observed using the pAER::LUC reporter (Section 2.2). *CYP81D11* showed the strongest induction by β-cc (80-fold), whereas it increased only modestly under RT-EL (5-fold) and remained low in RT-EL plants regardless of β-cc pretreatment. However, strong induction alone of these genes by exogenous β-cc may not be sufficient to protect photosynthetic performance under RT-EL

**Figure S3:**
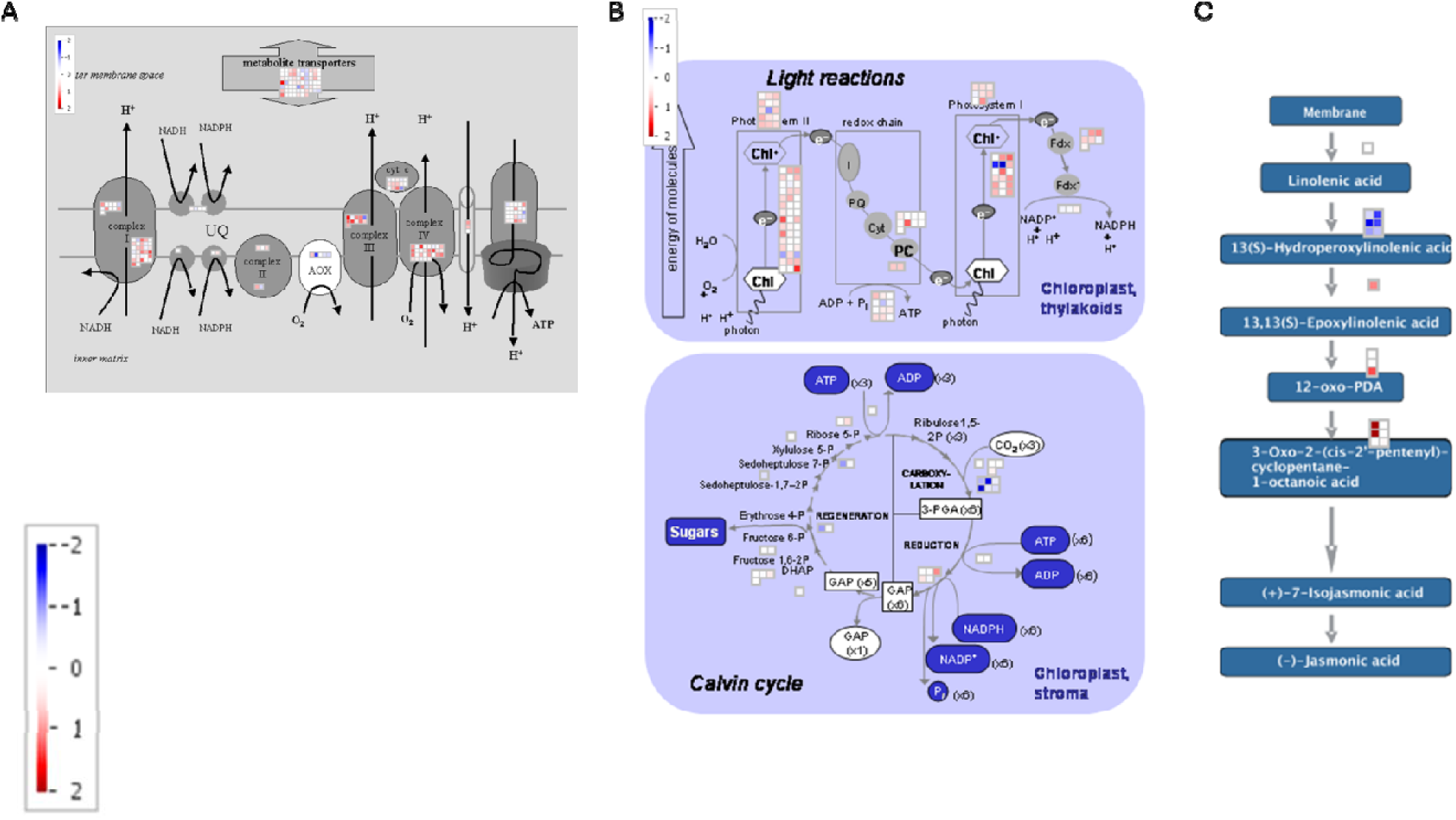
MapMan overview of pathways significantly affected by β-cc treatment. Differentially expressed genes (DEGs) from the β-cc vs control comparison were mapped onto MapMan functional bins. Panels highlight (A) mitochondrial oxidative phosphorylation, (B) photosynthetic light reactions, and (C) jasmonate biosynthesis. Color scale indicates direction and magnitude of expression changes (log2 fold change), with upregulated and downregulated genes shown accordingly.

**Figure S4:**
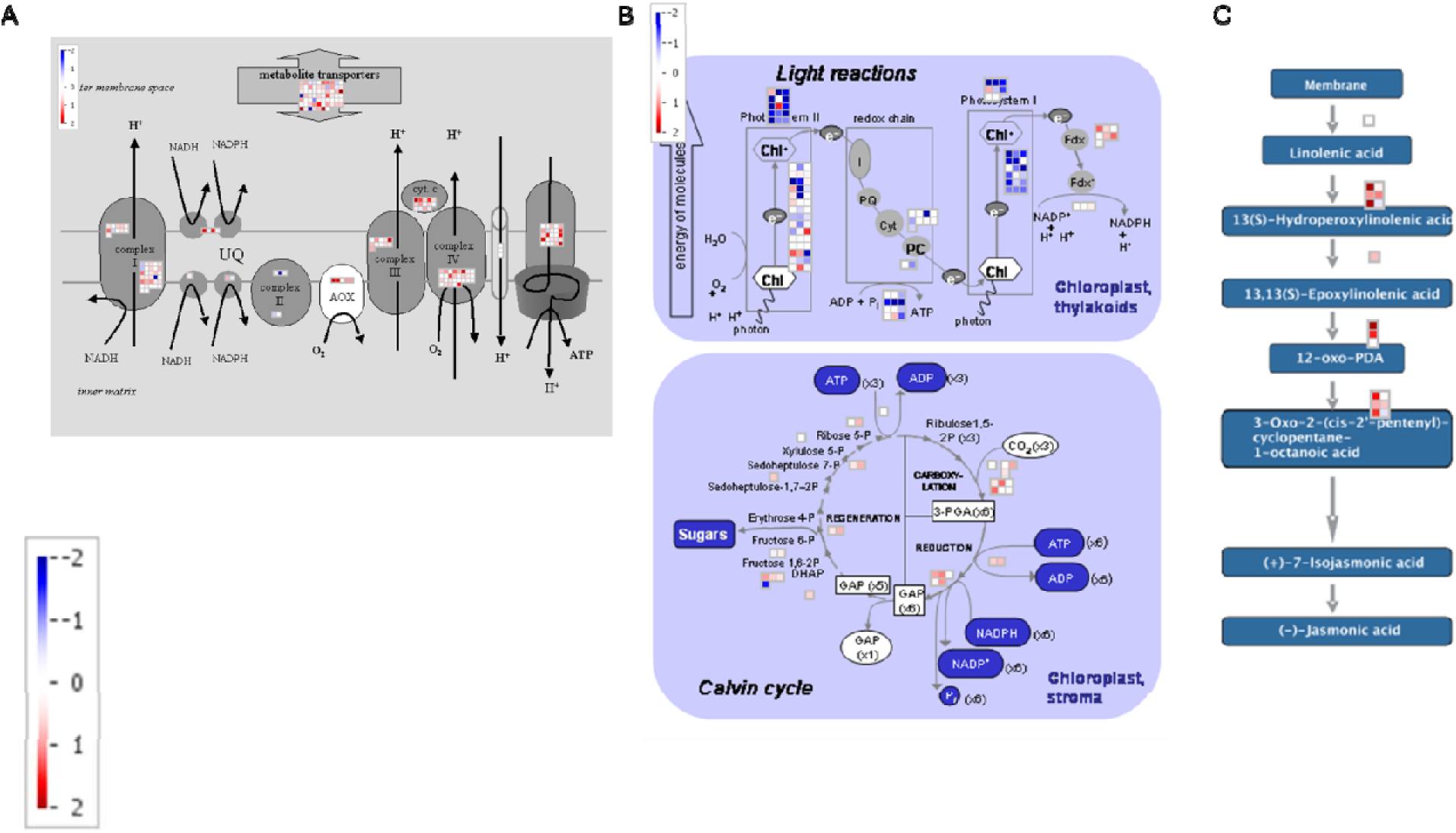
MapMan overview of pathways significantly affected by RT-EL treatment. Differentially expressed genes (DEGs) from the RT-EL vs control comparison were mapped onto MapMan functional bins. Panels highlight (A) mitochondrial oxidative phosphorylation, (B) photosynthetic light reactions, and (C) jasmonate biosynthesis. Color scale indicates direction and magnitude of expression changes (log2 fold change), with upregulated and downregulated genes shown accordingly.

Finally, we asked whether the transcriptional response induced by our RT-EL regime was consistent with high-light responses reported in independent Arabidopsis experiments. We therefore compared the RT-EL and β-cc transcriptional profiles with 12 published high-light conditions derived exclusively from adult leaf or rosette tissues and spanning different irradiances and exposure times (Bobrovskikh *et al*., 2025). PCA was performed on log2 fold-change profiles for genes tested across all datasets and significantly regulated (adjusted P < 0.01) in at least one condition. Despite the substantial differences in experimental setup among studies, the published high-light conditions occupied a relatively coherent region of the PCA space. Our RT-EL response localized within this distribution, indicating that the transcriptional program induced by RT-EL is broadly consistent with the range of responses independently observed in Arabidopsis leaves exposed to high irradiance (Figure S5). By contrast, the β-cc transcriptional response was clearly separated from the high-light cluster.

**Figure S5:**
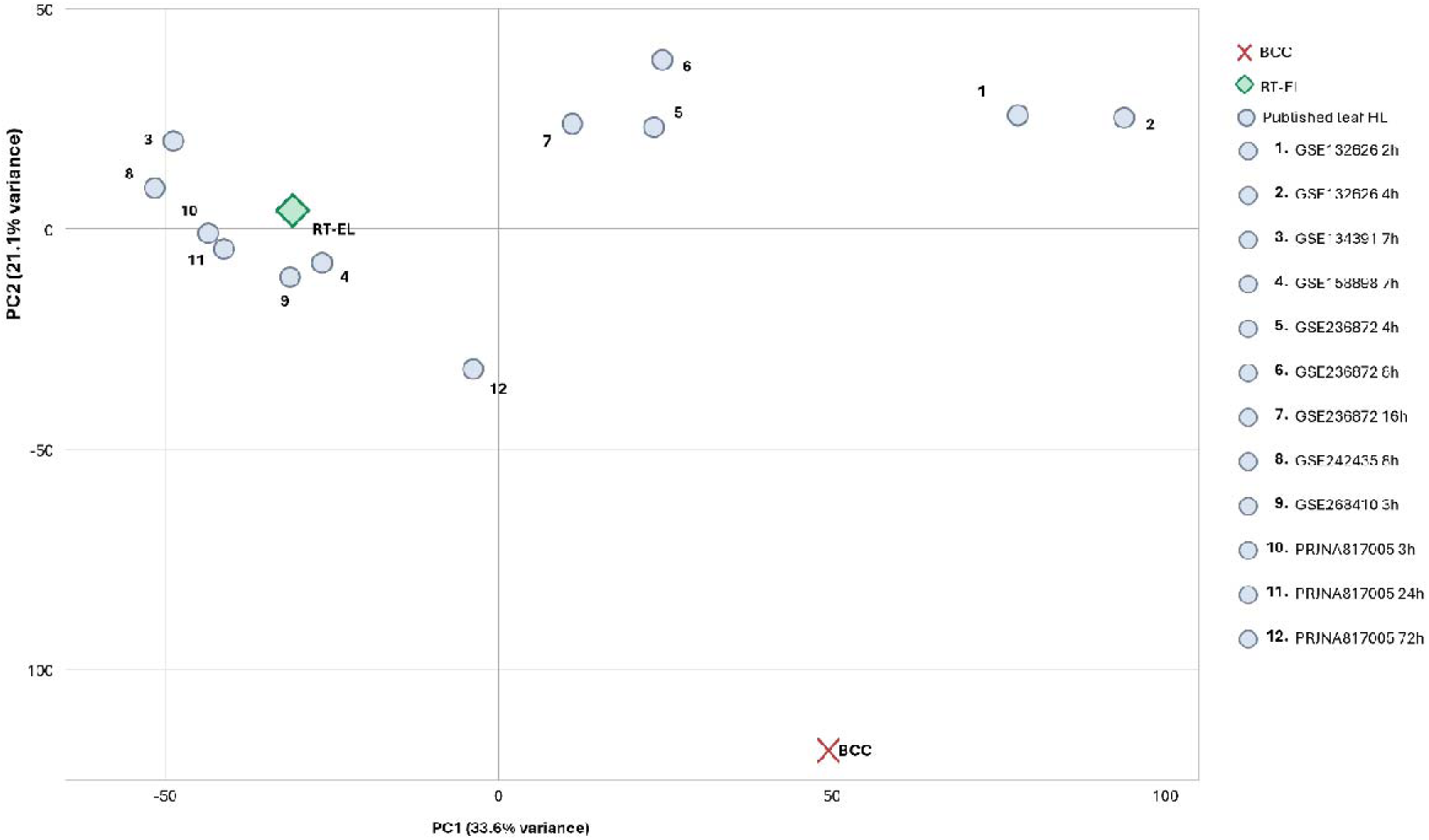
Comparison of the RT-EL and β-cyclocitral transcriptional responses with independent Arabidopsis leaf high-light transcriptomes. PCA was performed using log_2_ fold-change profiles from 12 published high-light conditions obtained from adult leaf or rosette tissues and the RT-EL and β-cc contrasts generated in this study. The analysis included 11,744 genes quantified across all datasets and significantly regulated (adjusted P < 0.01) in at least one condition.

## 4 Discussion

In this study, we investigated the kinetics of β-cc accumulation under RT-EL and of its signaling, and deepened the characterization of its effect on photosynthesis.

We discovered that exogenously supplied β-cc elicited cytosolic calcium changes within minutes and activated the pAER::LUC detoxification reporter within less than one hour, demonstrating that plant cells are able to perceive and respond to β-cc on a very short timescale. Thus, β-cc signaling itself is intrinsically rapid and is unlikely to represent a kinetic bottleneck in the response to photooxidative stress.

This rapid signaling contrasted with the kinetics of endogenous β-cc formation under room-temperature excess-light regime. Despite exposure to 1500 μmol photons m⁻² s⁻¹, plants maintained leaf temperatures close to ambient and established major photosynthetic adjustments before a detectable increase in β-cc or β-cyclocitric acid occurred. NPQ increased strongly within the first 4 h and approached a new steady state by 16 h, whereas endogenous β-cc and β-cca remained near control levels at 4 h and accumulated only after prolonged exposure. Under conditions in which excitation pressure can initially be accommodated by photochemical and non-photochemical energy dissipation, detectable β-carotene oxidation products accumulate only later. Accordingly, β-cc does not appear to be required for photosynthetic adaptation. Indeed, β-cc signaling is rather different from other chloroplastic retrograde signals as it does not reproduce the characteristic repression of nuclear-encoded photosynthetic genes observed during EL.

Once produced, however, β-cc is capable of initiating cellular signaling within minutes and exogenous β-cc induced extensive transcriptional and hormonal reprogramming. Interestingly, we did not see a strong induction of SA by β-cc under excessive light, indicating a marked difference from the response to excessive light at lower temperature.

Moreover, the β-cc and RT-EL transcriptional programs were largely distinct and frequently opposed, with the RT-EL response predominating when the two stimuli were combined. These observations suggest that β-cc signaling is not simply a signaling molecule produced under excess light, but rather represents a distinct response to β-carotene oxidation whose physiological relevance increases as the photooxidative component of the stress becomes more pronounced.

We therefore propose a kinetic model in which the earliest response to excess irradiance is dominated by photosynthetic regulation and energy dissipation. If high excitation persists and oxidative pressure increases, β-carotene oxidation progressively generates β-cc and related apocarotenoids. Because β-cc perception is rapid, their appearance can then rapidly engage calcium signaling, detoxification, and broader transcriptional responses. The relative timing of these processes is expected to depend strongly on the photooxidative severity of the environment, providing a possible explanation for the earlier β-cc accumulation and stronger β-cc-dependent protection previously observed when excess light is combined with low temperature.

## Author contributions: CRediT

Data acquisition and analyses: C.M., T.E., K.K.P., P.A.S., M.G.; Supervision and writing: D.S., A.C.; Funding acquisition: D.S.

## Acknowledgements

We thank Michel Havaux for kindly providing the pAER::LUC reporter line.

## Supplementary Material

The following supplementary data are available at JXB online.

Figure S1. Leaf temperature during RT-EL treatment.

Figure S2. Interaction and rank-overlap analyses of β-cc and RT-EL transcriptional responses.

Figure S3. MapMan overview of pathways affected by β-cc.

Figure S4. MapMan overview of pathways affected by RT-EL.

Figure S5. Comparison with independent Arabidopsis high-light transcriptomes.

Supplemental Data 1. Complete transcriptomic and functional-enrichment datasets (Figure 6 and S2).

Supplemental Data 2. Scripts for statistical analyses and plots (Figure 3, 4, 5 and S1)

Supplemental Data 3. Scripts and data for figure S5

Supplemental Data 4. Scripts and data for figure 7

## Data availability

The RNA-seq data generated in this study are available in [GEO/SRA/ENA] under accession XXXXX. All other data supporting the findings of this study are available within the article and its supplementary data.

## Use of Artificial Intelligence

ChatGPT (OpenAI), academic account, was used as a supporting tool to improve the clarity, grammar, and overall quality of the English language used in this work. It was also used to assist with the development, revision, and debugging of Python and R scripts. All AI-generated suggestions, text revisions, and code were critically reviewed, tested, and validated by the authors.

## Abbreviations

ABA: abscisic acid
β-cc: β-cyclocitral
β-cca: β-cyclocitric acid
EL: excess light
JA: jasmonic acid
NPQ: non-photochemical quenching
RT-EL: room-temperature excess light
SA: salicylic acid

