## Supplementary figures and images for "β-cyclocitral elicits rapid signaling distinct from the excess-light response in Arabidopsis"

### Figure_S1

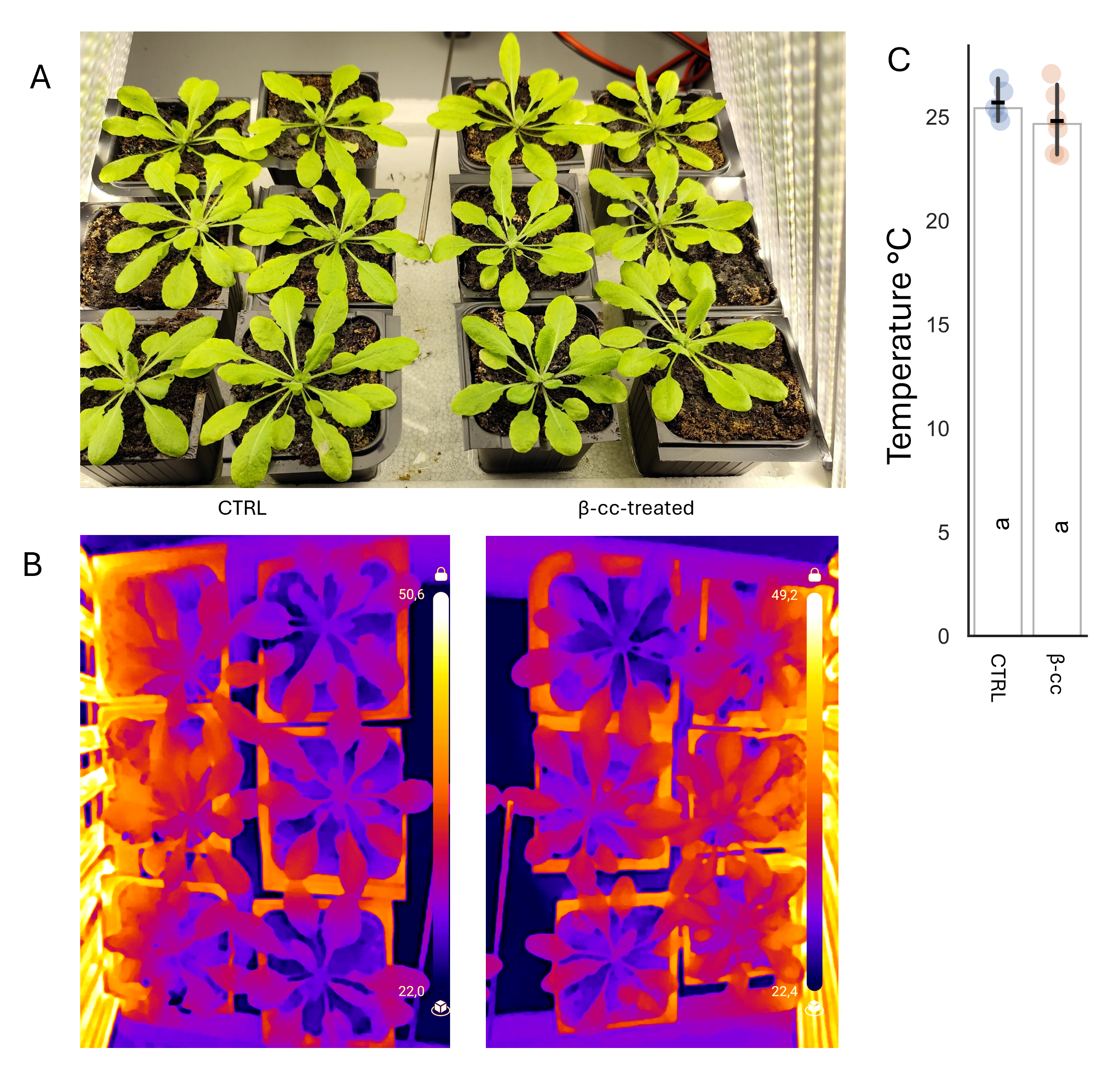

### Figure_S2

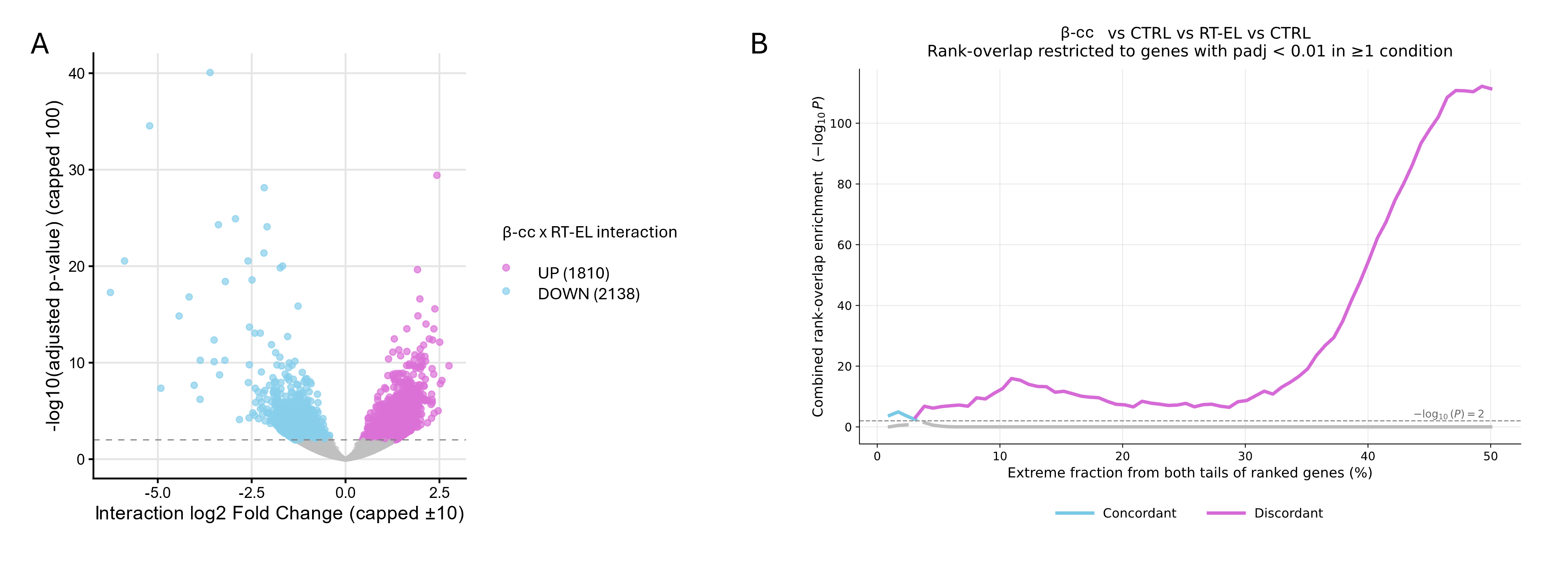

### Figure_S3

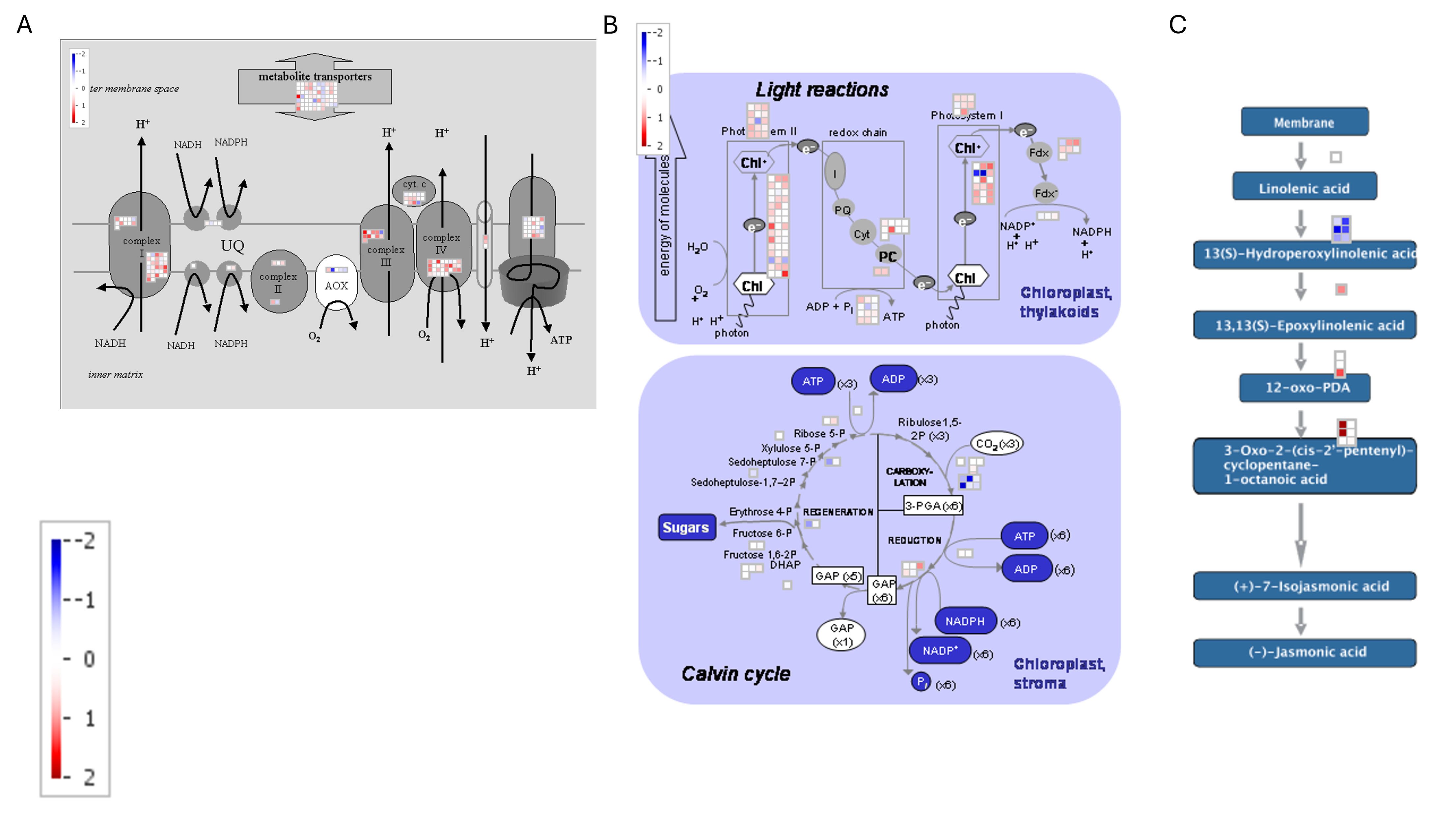

### Figure_S4

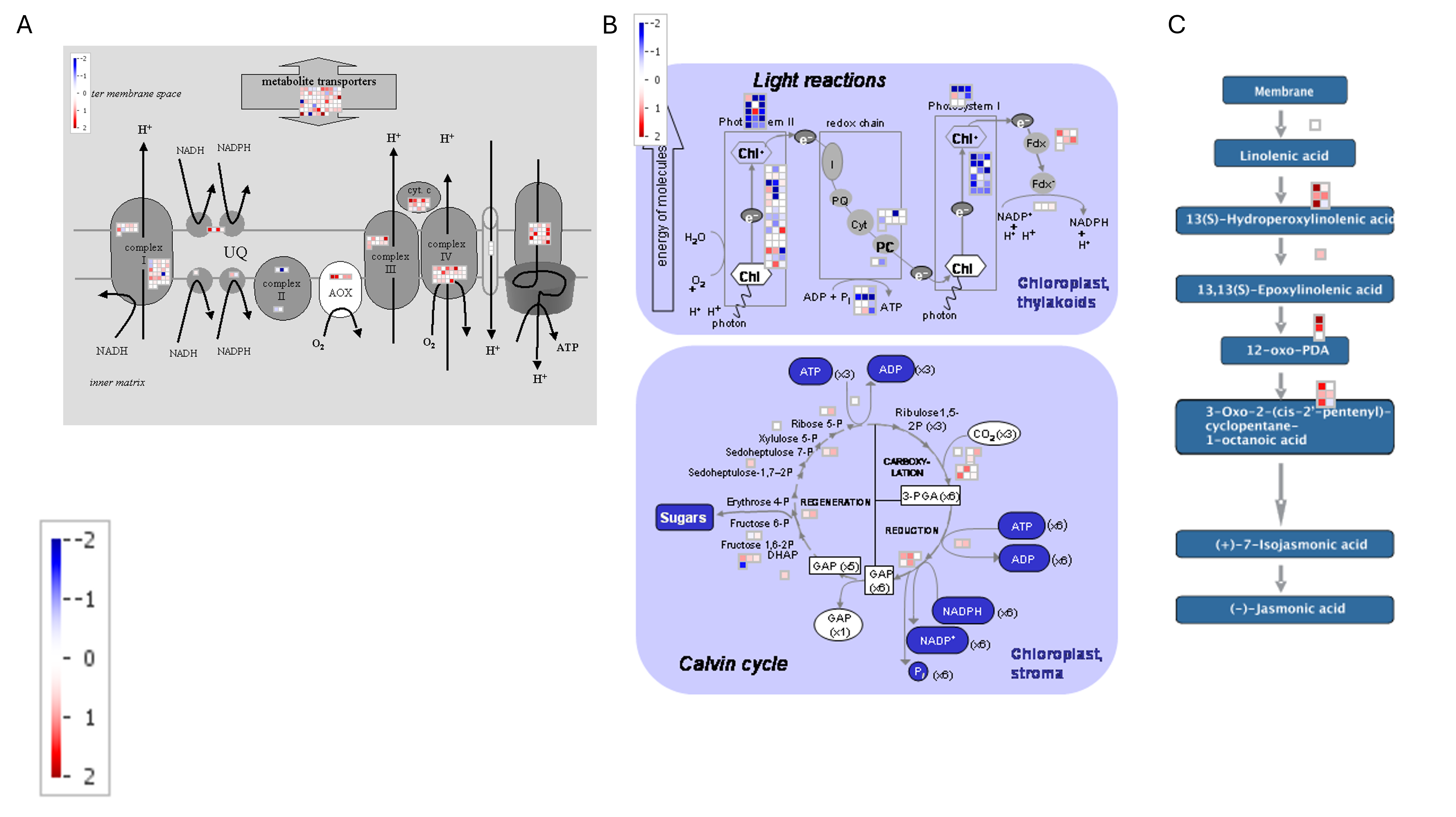

### Figure_S5

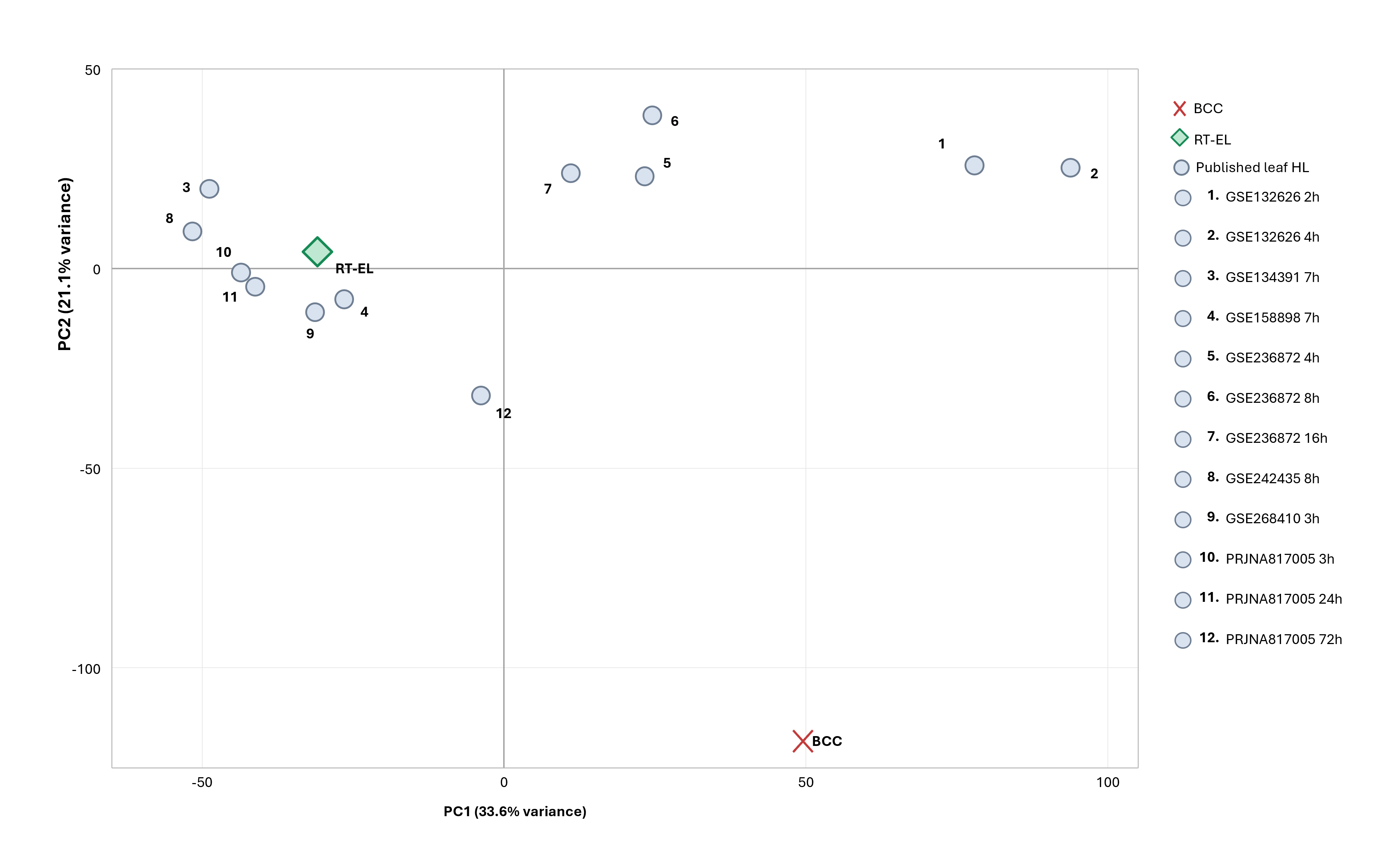
