## Supplemental_data_1 for "β-cyclocitral elicits rapid signaling distinct from the excess-light response in Arabidopsis": Supplemental_data_1.html

RNA-seq analysis based on Salmon and DESeq2


Code 

- Show All Code
- Hide All Code

### RNA-seq analysis based on Salmon and DESeq2

###### Stefano D’Alessandro

###### 29 settembre 2026

- 1 Abstract
- 2 Parameters
  - 2.1 Active modules
- 3 Data & Import
- 4 Input targets (Salmon
  quant.sf)
- 5 Differential expression (tximport +
  DESeq2)
- 6 Factorial model for the BCC by
  excess-light interaction
- 7
  Raw counts per sample
- 8 Total DEG vs Control
- 9 List of DEGs for table
- 10 PCA Plot
- 11 PCA Plot (Expressed Genes)
- 12 Heatmaps
- 13 Sample Distance Heatmap
- 14 Specific Markers
- 15 Organelles
- 16 MA Plots
- 17 Volcano Plot
- 18 Venn Diagram: look only at gene
  directions
- 19 GO Enrichment
- 20 GO Enrichment — *Physcomitrium
  patens*
- 21 KEGG ENRICHMENT
- 22 GSEA
- 23 GSEA GO — *Physcomitrium
  patens*
- 24 MAPMAN GO Enrichment
- 25 MAPMAP HEAT
- 26 TF enrichment based on
  PlantTFDB
- 27 Reproducibility

#### 1 Abstract

This report describes a reproducible RNA-seq differential expression
workflow based on **Salmon** transcript quantification and
**DESeq2** gene-level inference (via
**tximport**). It includes quality checks, differential
expression results, and functional enrichment analyses.

#### 2 Parameters

These values control filtering, significance thresholds, and
enrichment-term simplification. You can edit the defaults in the
**setup** chunk or in the launcher script.

- **Minimum filter**: 10
- **padj threshold**: 0.01
- **log2 fold-change threshold (+/-)**: 1
- **Cutoff for simplifying GO enrichment**: 0.3
- **Physcomitrium GO annotation**:
  physcomitrium\_GO.tsv

##### 2.1 Active modules

| Module | Status |
| --- | --- |
| Raw counts | ✅ |
| DEG bar chart | ✅ |
| DEG table | ✅ |
| Factorial BCC x EL | ✅ |
| PCA + Scree | ✅ |
| Heatmap variable | ✅ |
| Heatmap markers | ✅ |
| Organelles (SUBA) | ✅ |
| Volcano + MA | ✅ |
| Venn diagrams | ✅ |
| GO enrichment (TAIR) | ✅ |
| GO enrichment (Physcomitrium) | ⬜ |
| GSEA | ✅ |
| MapMan enrichment | ✅ |
| MapMan bin heatmap | ✅ |
| TF enrichment | ✅ |

#### 3 Data & Import

```
suppressPackageStartupMessages({
  library(tximport)
  library(readr)
  library(DESeq2)
  library(ggplot2)
  library(ggrepel)
  library(pheatmap)
  library(clusterProfiler)
  library(org.At.tair.db)        
  library(AnnotationDbi)
  library(tidyr)
  library(BiocParallel)
  library(fs)
  library(purrr)
  library(dplyr)
  library(stats)
  library(VennDiagram)
  library(readxl)
  library(gprofiler2)
  library(scales)
  library(svglite)
})

#cores
cores <- max(1, parallel::detectCores()-1)
BPPARAM <- SnowParam(workers = cores, type = "SOCK")
#register(BPPARAM)
```

#### 4 Input targets (Salmon quant.sf)

```
sample_dirs <- list.dirs(quant_dir, full.names = TRUE, recursive = FALSE)
files <- file.path(sample_dirs, "quant.sf")
names(files) <- basename(sample_dirs)

if (!file.exists("target_file.txt")) {
  sample_table <- data.frame(sample = basename(sample_dirs), condition = rep("control", length(sample_dirs)))
  write.table(sample_table, "target_file.txt", sep = "\t", quote = FALSE, row.names = FALSE)
  stop("File target_file.txt generated. Modify and relaunch the analysis.")
}
```

#### 5 Differential expression (tximport + DESeq2)

```
# ------------------------------------------------------------
# tximport + DESeq2 all-vs-all contrasts with shrinkage
# ------------------------------------------------------------

# Read sample sheet
samples <- read.delim("target_file.txt", stringsAsFactors = FALSE)

# Basic hygiene: trim sample names, ensure required columns exist, no duplicates
stopifnot(all(c("sample", "condition") %in% colnames(samples)))
samples$sample <- trimws(samples$sample)
stopifnot(!anyDuplicated(samples$sample))

# Build quant.sf paths and keep only existing ones
files_all <- file.path(quant_dir, samples$sample, "quant.sf")
names(files_all) <- samples$sample

exists_sf <- file.exists(files_all)
if (any(!exists_sf)) {
  warning("quant.sf missing for: ", paste(names(files_all)[!exists_sf], collapse = ", "))
}

files <- files_all[exists_sf]
if (length(files) == 0) stop("No quant.sf files found. Check quant_dir and sample names.")

# Reorder 'samples' to match 'files' exactly
idx <- match(names(files), samples$sample)
if (anyNA(idx)) stop("Match failed: some file names were not found in samples$sample. Check naming/whitespace.")
samples <- samples[idx, , drop = FALSE]

# Set rownames for DESeq2 and final safety check
rownames(samples) <- samples$sample
stopifnot(identical(rownames(samples), names(files)))

# Factorize condition (keep input order, but you can set explicit levels if you prefer)
samples$condition <- factor(samples$condition, levels = unique(samples$condition))

# Encode the biological 2x2 design explicitly.
# If BCC and EL columns are present in target_file.txt they are used and
# validated. Otherwise they are derived from the four condition labels.
factorial_key <- data.frame(
  condition = c("AtCTRL", "AtBCC", "AtHL", "AtBCCHL"),
  BCC = c("No", "Yes", "No", "Yes"),
  EL  = c("No", "No", "Yes", "Yes"),
  stringsAsFactors = FALSE
)

if (!all(c("BCC", "EL") %in% colnames(samples))) {
  samples <- samples %>%
    select(-any_of(c("BCC", "EL"))) %>%
    left_join(factorial_key, by = "condition")
} else {
  expected_design <- factorial_key[match(samples$condition, factorial_key$condition), ]
  mismatch <- is.na(expected_design$condition) |
    samples$BCC != expected_design$BCC |
    samples$EL != expected_design$EL
  if (any(mismatch)) {
    stop(
      "BCC/EL columns do not match condition labels for: ",
      paste(samples$sample[mismatch], collapse = ", ")
    )
  }
}

if (anyNA(samples[, c("BCC", "EL")])) {
  stop(
    "Cannot derive the factorial design. Expected conditions: ",
    paste(factorial_key$condition, collapse = ", ")
  )
}

samples$BCC <- factor(samples$BCC, levels = c("No", "Yes"))
samples$EL  <- factor(samples$EL,  levels = c("No", "Yes"))

# dplyr joins do not preserve custom row names. Restore the sample IDs because
# DESeq2 requires colData row names to match the tximport matrix columns.
rownames(samples) <- samples$sample
stopifnot(identical(rownames(samples), names(files)))

# All four BCC x EL combinations must be represented.
factorial_cells <- table(samples$BCC, samples$EL)
if (any(factorial_cells == 0L)) {
  stop("The factorial design is incomplete: at least one BCC x EL cell has no samples.")
}

#Names for successive analyses
control_cond <- levels(samples$condition)[1]
samples$condition <- relevel(samples$condition, ref = control_cond)
samples$condition <- factor(samples$condition, levels = c(control_cond, setdiff(levels(samples$condition), control_cond)))
conds <- levels(samples$condition)
other_conds <- conds[-1]

# Read tx2gene and import Salmon
tx2gene <- read.delim(
  file.path(db_dir, "tx2gene_folder/tx2gene.tsv"),
  header = FALSE,
  stringsAsFactors = FALSE
)
colnames(tx2gene) <- c("TranscriptID", "GeneID_X", "GeneID", "E", "Description", "Name")

txi <- tximport(files, type = "salmon", tx2gene = tx2gene)

# Extra safety: tximport columns must match colData rownames
stopifnot(identical(colnames(txi$counts), rownames(samples)))

# Build DESeq2 dataset and prefilter
dds <- DESeqDataSetFromTximport(txi, colData = samples, design = ~ condition)
dds <- dds[rowSums(counts(dds)) > filtro, ]
dds$condition <- droplevels(dds$condition)

# Fit ONCE
suppressWarnings(
  dds <- DESeq(dds, parallel = FALSE)
)

gene_names <- rownames(dds)
conds <- levels(dds$condition)

# ------------------------------------------------------------
# All-vs-all contrasts
# Use 'ashr' shrinkage for uniform support across arbitrary contrasts
# ------------------------------------------------------------

pairs <- combn(conds, 2, simplify = FALSE)

get_pair_res <- function(dds, treat, ctrl, tx2gene, alpha = 0.05, shrink_type = "ashr") {
  gene_names <- rownames(dds)
  
  # "treat vs ctrl": log2FC > 0 means higher in treat
  res <- DESeq2::results(dds, contrast = c("condition", treat, ctrl), alpha = alpha)

  # Shrink LFC while keeping p-values/padj from 'res'
  res_sh <- DESeq2::lfcShrink(dds, contrast = c("condition", treat, ctrl), res = res, type = shrink_type)

  res_df <- as.data.frame(res_sh)
  res_df$stat_unshrunk <- res$stat
  res_df <- res_df[gene_names, , drop = FALSE]
  idx <- match(rownames(res_df), tx2gene$GeneID_X)
  res_df$GeneID <- tx2gene$GeneID[idx]
  res_df
}

# Parallel computation of all contrasts 
res_pair_list <- bplapply(pairs, function(p) {
  a <- p[1]
  b <- p[2]
  name <- paste0(b, "_vs_", a)  # convention: second vs first
  df <- get_pair_res(dds, treat = b, ctrl = a, tx2gene = tx2gene, shrink_type = "ashr")
  list(name = name, df = df)
}, BPPARAM = BPPARAM)

res_list <- setNames(
  lapply(res_pair_list, `[[`, "df"),
  vapply(res_pair_list, `[[`, character(1), "name")
)

# Sanity check: number of pairwise contrasts
expected <- length(conds) * (length(conds) - 1) / 2
stopifnot(length(res_list) == expected)

# ------------------------------------------------------------
# Ensure each result includes ALL genes (fill missing rows with NA)
# Do it in a type-safe way (avoid matrix(NA) coercion issues)
# ------------------------------------------------------------

all_genes <- unique(tx2gene$GeneID_X)  # gene IDs (2nd column)

res_list <- lapply(res_list, function(res_df) {
  missing_genes <- setdiff(all_genes, rownames(res_df))

  if (length(missing_genes) > 0) {
    # Create an NA "template" with the same columns/classes as res_df
    na_df <- res_df[rep(1, length(missing_genes)), , drop = FALSE]
    na_df[,] <- NA
    rownames(na_df) <- missing_genes

    # Keep GeneID consistent if present
    if ("GeneID" %in% colnames(na_df)) na_df$GeneID <- missing_genes

    res_df <- rbind(res_df, na_df)
  }

  # Reorder rows to match all_genes
  res_df <- res_df[all_genes, , drop = FALSE]
  res_df
})

# Save
dir_create("Report")
saveRDS(res_list, file = "Report/res_list_all_genes.rds")
```

#### 6 Factorial model for the BCC by excess-light interaction

The four-level `condition` model above remains useful for
ordinary pairwise comparisons. The factorial model below fits the same
four group means but tests the biological interaction explicitly:

\[
(AtBCCHL - AtHL) - (AtBCC - AtCTRL)
\]

A positive interaction means that the BCC effect becomes more
positive under EL. A negative interaction means that the BCC effect
becomes less positive, or more negative, under EL. The interaction is
therefore not another version of the `AtBCCHL versus AtHL`
comparison; it tests whether that BCC effect differs from
`AtBCC versus AtCTRL`.

```
if (run_factorial_model) {

  factorial_dir <- file.path("Report", "factorial_BCC_x_EL")
  dir_create(factorial_dir)

  # Fit the full 2x2 model. With all four cells represented, this model and
  # ~ condition have the same fitted group means; the parameterization and
  # biological contrasts differ.
  dds_factorial <- DESeqDataSetFromTximport(
    txi,
    colData = samples,
    design = ~ BCC + EL + BCC:EL
  )
  dds_factorial <- dds_factorial[rowSums(counts(dds_factorial)) > filtro, ]
  dds_factorial$BCC <- relevel(droplevels(dds_factorial$BCC), ref = "No")
  dds_factorial$EL  <- relevel(droplevels(dds_factorial$EL),  ref = "No")
  design(dds_factorial) <- ~ BCC + EL + BCC:EL

  suppressWarnings(
    dds_factorial <- DESeq(dds_factorial, parallel = FALSE)
  )

  stopifnot(identical(rownames(dds), rownames(dds_factorial)))

  factorial_names <- resultsNames(dds_factorial)
  bcc_coef <- grep("^BCC_.*_vs_", factorial_names, value = TRUE)
  el_coef  <- grep("^EL_.*_vs_",  factorial_names, value = TRUE)
  interaction_coef <- factorial_names[
    grepl("BCC", factorial_names) &
      grepl("EL", factorial_names) &
      !grepl("_vs_", factorial_names)
  ]

  if (length(bcc_coef) != 1L || length(el_coef) != 1L || length(interaction_coef) != 1L) {
    stop(
      "Could not identify factorial coefficients. resultsNames(dds_factorial): ",
      paste(factorial_names, collapse = ", ")
    )
  }

  get_factorial_result <- function(dds_obj, label, coef_name = NULL, contrast_spec = NULL) {
    if (!is.null(coef_name)) {
      res_raw <- DESeq2::results(dds_obj, name = coef_name, alpha = padj)
      res_shrunk <- DESeq2::lfcShrink(
        dds_obj,
        coef = coef_name,
        res = res_raw,
        type = "ashr"
      )
    } else {
      res_raw <- DESeq2::results(dds_obj, contrast = contrast_spec, alpha = padj)
      res_shrunk <- DESeq2::lfcShrink(
        dds_obj,
        contrast = contrast_spec,
        res = res_raw,
        type = "ashr"
      )
    }

    out <- as.data.frame(res_shrunk)
    out$stat_unshrunk <- res_raw$stat
    out$GeneID_X <- rownames(out)

    annotation_idx <- match(out$GeneID_X, tx2gene$GeneID_X)
    out$GeneID <- tx2gene$GeneID[annotation_idx]
    out$Name <- tx2gene$Name[annotation_idx]
    out$Description <- tx2gene$Description[annotation_idx]
    out$contrast <- label
    out$direction <- dplyr::case_when(
      !is.na(out$padj) & out$padj < padj & out$log2FoldChange >  log2foldchange ~ "UP",
      !is.na(out$padj) & out$padj < padj & out$log2FoldChange < -log2foldchange ~ "DOWN",
      TRUE ~ "not_significant"
    )

    out <- out[, c(
      "GeneID_X", "GeneID", "Name", "Description", "contrast",
      "baseMean", "log2FoldChange", "lfcSE", "stat_unshrunk",
      "pvalue", "padj", "direction"
    )]
    rownames(out) <- out$GeneID_X
    out
  }

  factorial_results <- list(
    BCC_at_noEL = get_factorial_result(
      dds_factorial,
      label = "BCC effect without EL: AtBCC - AtCTRL",
      coef_name = bcc_coef
    ),
    EL_at_noBCC = get_factorial_result(
      dds_factorial,
      label = "EL effect without BCC: AtHL - AtCTRL",
      coef_name = el_coef
    ),
    BCC_at_EL = get_factorial_result(
      dds_factorial,
      label = "BCC effect under EL: AtBCCHL - AtHL",
      contrast_spec = list(c(bcc_coef, interaction_coef))
    ),
    EL_at_BCC = get_factorial_result(
      dds_factorial,
      label = "EL effect after BCC: AtBCCHL - AtBCC",
      contrast_spec = list(c(el_coef, interaction_coef))
    ),
    BCC_x_EL_interaction = get_factorial_result(
      dds_factorial,
      label = "Interaction: (AtBCCHL - AtHL) - (AtBCC - AtCTRL)",
      coef_name = interaction_coef
    )
  )

  # Save complete result tables, not only thresholded DEGs.
  purrr::iwalk(factorial_results, function(tbl, nm) {
    readr::write_tsv(
      tibble::rownames_to_column(tbl, var = "row_id") %>%
        select(-row_id),
      file.path(factorial_dir, paste0(nm, ".tsv"))
    )
  })
  saveRDS(dds_factorial, file.path(factorial_dir, "dds_factorial.rds"))
  saveRDS(factorial_results, file.path(factorial_dir, "factorial_results.rds"))

  factorial_summary <- purrr::imap_dfr(factorial_results, function(tbl, nm) {
    data.frame(
      contrast = nm,
      significant_padj_only = sum(
        !is.na(tbl$padj) & tbl$padj < padj,
        na.rm = TRUE
      ),
      up = sum(tbl$direction == "UP", na.rm = TRUE),
      down = sum(tbl$direction == "DOWN", na.rm = TRUE),
      total = sum(tbl$direction %in% c("UP", "DOWN"), na.rm = TRUE)
    )
  })
  readr::write_tsv(
    factorial_summary,
    file.path(factorial_dir, "factorial_contrast_summary.tsv")
  )

  knitr::kable(
    factorial_summary,
    caption = paste0(
      "Factorial contrasts. significant_padj_only uses padj < ", padj,
      "; up, down and total additionally require |log2FC| > ", log2foldchange
    )
  )

  # The interaction volcano is the direct gene-wise test of whether the
  # BCC response differs between no-EL and EL backgrounds.
  interaction_df <- factorial_results$BCC_x_EL_interaction %>%
    mutate(
      log2FoldChange_capped = pmax(pmin(log2FoldChange, 10), -10),
      minus_log10_padj_capped = pmin(
        -log10(pmax(padj, .Machine$double.xmin)),
        100
      ),
      plot_group = dplyr::case_when(
        !is.na(.data$padj) & .data$padj < .env$padj &
          log2FoldChange > 0 ~ "positive",
        !is.na(.data$padj) & .data$padj < .env$padj &
          log2FoldChange < 0 ~ "negative",
        TRUE ~ "ns"
      ),
      plot_group = factor(plot_group, levels = c("positive", "negative", "ns"))
    )

  interaction_positive_n <- sum(interaction_df$plot_group == "positive", na.rm = TRUE)
  interaction_negative_n <- sum(interaction_df$plot_group == "negative", na.rm = TRUE)
  interaction_color_labels <- c(
    "positive" = paste0("POSITIVE (", interaction_positive_n, ")"),
    "negative" = paste0("NEGATIVE (", interaction_negative_n, ")")
  )

  p_interaction <- ggplot(
    interaction_df,
    aes(x = log2FoldChange_capped, y = minus_log10_padj_capped)
  ) +
    geom_point(aes(color = plot_group, size = plot_group), alpha = 0.7) +
    scale_size_manual(
      values = c("positive" = 1.8, "negative" = 1.8, "ns" = 0.9),
      guide = "none"
    ) +
    geom_hline(
      yintercept = -log10(padj),
      linetype = "dashed", color = "grey50", linewidth = 0.4
    ) +
    scale_color_manual(
      values = c("positive" = "orchid", "negative" = "skyblue", "ns" = "grey75"),
      breaks = c("positive", "negative"),
      labels = interaction_color_labels,
      name = "BCC × RT-EL interaction"
    ) +
    labs(
      x = "Interaction log2 Fold Change (capped ±10)",
      y = "-log10(adjusted p-value) (capped 100)"
    ) +
    theme_classic(base_size = 14) +
    theme(panel.grid.major = element_line(color = "grey90"))

  print(p_interaction)
  ggsave(
    file.path(factorial_dir, "volcano_BCC_x_RT-EL_interaction.svg"),
    p_interaction,
    width = 7,
    height = 5,
    dpi = 600
  )

  # Among genes regulated by BCC without EL, visualize whether the interaction
  # has the opposite sign. This directly tests the global counteraction of the
  # BCC-responsive transcriptional program by EL.
  bcc_opposite_df <- factorial_results$BCC_at_noEL %>%
    select(
      GeneID_X,
      BCC_log2FC = log2FoldChange,
      BCC_padj = padj,
      BCC_direction = direction
    ) %>%
    inner_join(
      factorial_results$BCC_x_EL_interaction %>%
        select(
          GeneID_X,
          interaction_log2FC = log2FoldChange,
          interaction_padj = padj
        ),
      by = "GeneID_X"
    ) %>%
    filter(BCC_direction %in% c("UP", "DOWN")) %>%
    mutate(
      opposite_sign = sign(BCC_log2FC) != sign(interaction_log2FC),
      interaction_significant = !is.na(interaction_padj) &
        interaction_padj < .env$padj,
      plot_group = dplyr::case_when(
        opposite_sign & interaction_significant & BCC_direction == "UP" ~
          "BCC_UP_significant",
        opposite_sign & interaction_significant & BCC_direction == "DOWN" ~
          "BCC_DOWN_significant",
        opposite_sign ~ "opposite_not_significant",
        TRUE ~ "same_sign"
      ),
      plot_group = factor(
        plot_group,
        levels = c(
          "opposite_not_significant",
          "BCC_UP_significant",
          "BCC_DOWN_significant",
          "same_sign"
        )
      )
    )

  bcc_total_n <- nrow(bcc_opposite_df)
  opposite_n <- sum(bcc_opposite_df$opposite_sign, na.rm = TRUE)
  same_sign_n <- bcc_total_n - opposite_n
  bcc_up_sig_n <- sum(
    bcc_opposite_df$plot_group == "BCC_UP_significant",
    na.rm = TRUE
  )
  bcc_down_sig_n <- sum(
    bcc_opposite_df$plot_group == "BCC_DOWN_significant",
    na.rm = TRUE
  )
  opposite_not_sig_n <- sum(
    bcc_opposite_df$plot_group == "opposite_not_significant",
    na.rm = TRUE
  )

  opposite_color_labels <- c(
    "BCC_UP_significant" = paste0(
      "BCC UP; significant interaction (", bcc_up_sig_n, ")"
    ),
    "BCC_DOWN_significant" = paste0(
      "BCC DOWN; significant interaction (", bcc_down_sig_n, ")"
    ),
    "opposite_not_significant" = paste0(
      "Opposite sign; FDR ≥ ", padj, " (", opposite_not_sig_n, ")"
    ),
    "same_sign" = paste0("Same sign (", same_sign_n, ")")
  )

  p_opposite_sign <- ggplot(
    bcc_opposite_df,
    aes(x = BCC_log2FC, y = interaction_log2FC)
  ) +
    geom_point(
      aes(color = plot_group, shape = plot_group, size = plot_group),
      alpha = 0.70
    ) +
    geom_abline(
      slope = -1, intercept = 0,
      linetype = "dashed", color = "grey35", linewidth = 0.5
    ) +
    geom_hline(yintercept = 0, color = "grey55", linewidth = 0.35) +
    geom_vline(xintercept = 0, color = "grey55", linewidth = 0.35) +
    scale_color_manual(
      values = c(
        "BCC_UP_significant" = "orchid",
        "BCC_DOWN_significant" = "skyblue",
        "opposite_not_significant" = "grey75",
        "same_sign" = "black"
      ),
      breaks = names(opposite_color_labels),
      labels = opposite_color_labels,
      name = NULL
    ) +
    scale_shape_manual(
      values = c(
        "BCC_UP_significant" = 16,
        "BCC_DOWN_significant" = 16,
        "opposite_not_significant" = 16,
        "same_sign" = 4
      ),
      breaks = names(opposite_color_labels),
      labels = opposite_color_labels,
      name = NULL
    ) +
    scale_size_manual(
      values = c(
        "BCC_UP_significant" = 1.8,
        "BCC_DOWN_significant" = 1.8,
        "opposite_not_significant" = 0.9,
        "same_sign" = 2.2
      ),
      guide = "none"
    ) +
    coord_cartesian(xlim = c(-7, 7), ylim = c(-7, 7)) +
    labs(
      title = "EL counteracts the BCC-responsive transcriptional program",
      subtitle = paste0(
        format(opposite_n, big.mark = ","), "/",
        format(bcc_total_n, big.mark = ","),
        " BCC-responsive genes have an interaction of the opposite sign (",
        round(100 * opposite_n / bcc_total_n, 1), "%)"
      ),
      x = "BCC effect without EL (shrunken log2 Fold Change)",
      y = "BCC × RT-EL interaction (shrunken log2 Fold Change)"
    ) +
    theme_classic(base_size = 14) +
    theme(panel.grid.major = element_line(color = "grey90"))

  print(p_opposite_sign)
  ggsave(
    file.path(factorial_dir, "scatter_BCC_effect_vs_interaction_opposite_sign.svg"),
    p_opposite_sign,
    width = 8,
    height = 6,
    dpi = 600
  )

  opposite_sign_summary <- data.frame(
    BCC_responsive_genes = bcc_total_n,
    opposite_interaction_sign = opposite_n,
    opposite_percent = 100 * opposite_n / bcc_total_n,
    significant_interaction_padj = bcc_up_sig_n + bcc_down_sig_n,
    significant_interaction_percent =
      100 * (bcc_up_sig_n + bcc_down_sig_n) / bcc_total_n,
    same_sign = same_sign_n
  )
  readr::write_tsv(
    opposite_sign_summary,
    file.path(factorial_dir, "BCC_opposite_interaction_sign_summary.tsv")
  )

} # end run_factorial_model
```

#### 7 Raw counts per sample

```
if (run_raw_counts) {
raw_counts <- counts(dds, normalized = FALSE)
total_counts <- colSums(raw_counts)
df_counts <- data.frame(sample = names(total_counts), total = total_counts/ 1e6)

# Improved: color by condition, add mean line
df_counts$condition <- samples[df_counts$sample, "condition"]

# Sort samples alphabetically
df_counts$sample <- factor(df_counts$sample, levels = sort(unique(df_counts$sample)))

p_counts <- ggplot(df_counts, aes(x = sample, y = total, fill = condition)) +
  geom_bar(stat = "identity") +
  geom_hline(yintercept = mean(df_counts$total), linetype = "dashed", color = "grey40") +
  annotate("text", x = 0.7, y = mean(df_counts$total) * 1.03,
           label = paste0("mean: ", round(mean(df_counts$total), 1), "M"),
           hjust = 0, size = 3.5, color = "grey40") +
  coord_flip() +
  theme_classic(base_size = 13) +
  labs(title = "Total Raw Counts per Sample", x = NULL, y = "Total Counts (millions)", fill = "Condition")

print(p_counts)
ggsave("Report/total_counts_per_sample.svg", p_counts, dpi = 300)
} # end run_raw_counts
```

#### 8 Total DEG vs Control

```
if (run_deg_plot) {

# --- Count DEGs per condition ---
res_ctrl <- res_list[grep(paste0("_vs_", control_cond, "$"), names(res_list))]

deg_counts <- lapply(names(res_ctrl), function(nm) {
  res_df <- res_ctrl[[nm]]
  res_df <- res_df %>% filter(complete.cases(log2FoldChange, padj))
  
  up_count <- sum(res_df$padj < padj & res_df$log2FoldChange > log2foldchange, na.rm = TRUE)
  down_count <- sum(res_df$padj < padj & res_df$log2FoldChange < -log2foldchange, na.rm = TRUE)
  
  cond_name <- sub(paste0("_vs_", control_cond), "", nm)
  
  data.frame(
    condition = cond_name,
    up = up_count,
    down = down_count,
    total = up_count + down_count
  )
}) %>% bind_rows()

# --- DEGs for PCA & Heatmap ---
geni_variabili <- max(deg_counts$total, na.rm = TRUE)

# --- long for ggplot --- 
deg_counts_long <- deg_counts %>%
  pivot_longer(cols = c("up", "down"), 
               names_to = "direction", 
               values_to = "count")

deg_counts_long <- deg_counts_long %>%
  mutate(direction = factor(direction, levels = c("up", "down")))

# Plot
p_deg <- ggplot(deg_counts_long, aes(x = condition, y = count, fill = direction)) +
  geom_bar(stat = "identity", position = position_dodge(width = 0.9)) +
  geom_text(aes(label = count),
            position = position_dodge(width = 0.9),
            vjust = -0.2, size = 5) +
  scale_fill_manual(
    values = c("up" = "orchid", "down" = "skyblue"),
    labels = c("Upregulated", "Downregulated")
  ) +
  scale_y_continuous(expand = expansion(mult = c(0, 0.1))) +  # lascia il 10% di spazio sopra
  theme_classic(base_size = 14) +
  labs(
    title = paste0("Number of DEGs vs Control (", control_cond, ")"),
    x = "Treatment",
    y = "Number of DEGs",
    fill = "Direction"
  ) +
  theme(
    axis.text.x = element_text(angle = 45, hjust = 1)
  )

print(p_deg)

ggsave("Report/DEGs_per_condition.svg", p_deg, width = 7, height = 5, dpi = 300)


} else {
  res_ctrl <- res_list[grep(paste0("_vs_", control_cond, "$"), names(res_list))]
  deg_counts_tmp <- lapply(names(res_ctrl), function(nm) {
    res_df <- res_ctrl[[nm]] %>% dplyr::filter(complete.cases(log2FoldChange, padj))
    data.frame(total = sum(res_df$padj < padj & abs(res_df$log2FoldChange) > log2foldchange, na.rm=TRUE))
  })
  geni_variabili <- max(do.call(rbind, deg_counts_tmp)$total, na.rm = TRUE)
  res_ctrl <- res_list[grep(paste0("_vs_", control_cond, "$"), names(res_list))]
}
```

#### 9 List of DEGs for table

```
if (run_deg_table) {
# ------------------------------
# 1. Mean TPM per group
# ------------------------------
tpm_tx <- txi$abundance

tpm_gene <- sapply(levels(dds$condition), function(cond) {
  rowMeans(tpm_tx[, dds$condition == cond, drop = FALSE])
})

tpm_sd <- sapply(levels(dds$condition), function(cond) {
  apply(tpm_tx[, dds$condition == cond, drop = FALSE], 1, sd)
})

# Data frame 
tpm_gene <- data.frame(
  GeneID_X = rownames(tpm_tx),
  tpm_gene,
  check.names = FALSE
)
colnames(tpm_gene)[-1] <- paste0("TPM_", colnames(tpm_gene)[-1])

# First column: control 
control_sample <- paste0("TPM_", control_cond)
other_samples  <- paste0("TPM_", other_conds)
tpm_gene <- tpm_gene[, c("GeneID_X", control_sample, other_samples), drop = FALSE]
tpm_sd <- tpm_sd[, c(control_cond, other_conds), drop = FALSE]

# ------------------------------
# 2. Adding DESeq2 result for each comparison vs control
# ------------------------------
df_final <- tpm_gene

for(nm in names(res_ctrl)) {
  res <- res_ctrl[[nm]]
  res$GeneID <- rownames(res)
  
  # Remove duplicates and incomplete rows
  res <- res[!duplicated(res$GeneID), ]
  res <- res[complete.cases(res[, c("log2FoldChange", "padj")]), ]
  
  # Extract names (without vs_<control>)
  cond_name <- sub(paste0("_vs_", control_cond), "", nm)
  
  # Rename columns
  colnames(res)[colnames(res) == "log2FoldChange"] <- paste0("log2FC_", cond_name)
  colnames(res)[colnames(res) == "padj"] <- paste0("padj_", cond_name)
  
  res_sub <- res[, c("GeneID", paste0("log2FC_", cond_name), paste0("padj_", cond_name))]
  colnames(res_sub) <- c("GeneID_X", paste0("log2FC_", cond_name), paste0("padj_", cond_name))
  
  # Merge  df_final
  df_final <- merge(df_final, res_sub, by = "GeneID_X", all.x = TRUE)
}

# ========================================
# 2b Adding Mapping to At
# ========================================
colnames(tx2gene) <- c("TranscriptID", "GeneID_X", "GeneID", "E", "Description", "Name")

tx2gene_filtrato <- tx2gene %>%
  filter(E < 0.01) %>%
  select(GeneID_X, GeneID)

df_final <- df_final %>%
  left_join(tx2gene_filtrato, by = "GeneID_X")

# ========================================
# 3 Adding SUBA localization
# ========================================

# Importing SUBA database
suba_loc <- read.csv(file.path(db_dir, "19_8_2022_suba5_summary.csv"), header = TRUE, stringsAsFactors = FALSE)

colnames(suba_loc)[colnames(suba_loc) == "locus"] <- "GeneID"
colnames(suba_loc)[colnames(suba_loc) == "location_consensus"] <- "SUBA_Localization"
suba_loc$GeneID <- substr(suba_loc$GeneID, 1, 9)

# Removing duplicates and other columns
suba_loc <- suba_loc[, c("GeneID", "SUBA_Localization")]
suba_loc <- suba_loc[!duplicated(suba_loc$GeneID), ]

# Merge 
df_final <- merge(df_final, suba_loc, by = "GeneID", all.x = TRUE)

# ========================================
# 4. TAIR Annotation (Arabidopsis)
# ========================================

tx2gene <- tx2gene[!duplicated(tx2gene$GeneID), ]

# Adding Descritpion and name to df_final
df_final <- merge(df_final, tx2gene[, c("GeneID", "Name", "Description")],
                  by = "GeneID", all.x = TRUE)

df_final <- df_final[!duplicated(df_final$GeneID_X), ]


# ========================================
# 5 output
# ========================================

# Save the table
write.table(df_final,
            file = "Report/DEGs_per_condition_vs_control_TPM.tsv",
            sep = "\t",
            quote = FALSE,
            row.names = FALSE
            )

write.table(tpm_sd,
            file = "Report/Standard_deviation_for_TPM.tsv",
            sep = "\t",
            quote = FALSE,
            row.names = FALSE
            )

} # end run_deg_table
```

#### 10 PCA Plot

```
if (run_pca) {
vst_obj <- vst(dds, blind = TRUE)
pca_data <- plotPCA(vst_obj, intgroup = "condition", returnData = TRUE, ntop = geni_variabili)
percentVar <- round(100 * attr(pca_data, "percentVar"))

# Use sample name (last 10 chars) as label
pca_data$label <- pca_data$condition

p_pca <- ggplot(pca_data, aes(x = PC1, y = PC2, color = condition, fill = condition)) +
  stat_ellipse(geom = "polygon", alpha = 0.08, level = 0.9, linetype = "dashed") +
  geom_point(size = 4, shape = 21, color = "white", stroke = 0.5,
             aes(fill = condition)) +
  ggrepel::geom_text_repel(aes(label = label), size = 3.5, show.legend = FALSE,
                            box.padding = 0.4, segment.color = "grey60") +
  labs(title = "PCA – variable DEGs",
       x = paste0("PC1: ", percentVar[1], "% variance"),
       y = paste0("PC2: ", percentVar[2], "% variance"),
       color = "Condition", fill = "Condition") +
  theme_classic(base_size = 14) +
  theme(panel.grid.major = element_line(color = "grey90"),
        legend.position = "right") +
  coord_equal()

print(p_pca)
ggsave("Report/PCA_plot.svg", p_pca, dpi = 300, width = 8, height = 7)

# --- Scree plot: variance explained by each PC (same ntop as PCA variable genes) ---
rv <- rowVars(assay(vst_obj))
ntop_scree <- min(geni_variabili, sum(is.finite(rv)))
pca_full <- prcomp(t(assay(vst_obj)[order(rv, decreasing = TRUE)[seq_len(ntop_scree)], ]))
pct_var  <- round(100 * pca_full$sdev^2 / sum(pca_full$sdev^2), 2)
n_show   <- min(15, length(pct_var))
scree_df <- data.frame(PC = paste0("PC", seq_len(n_show)), Variance = pct_var[seq_len(n_show)],
                       Cumulative = cumsum(pct_var)[seq_len(n_show)])
scree_df$PC <- factor(scree_df$PC, levels = scree_df$PC)

p_scree <- ggplot(scree_df, aes(x = PC, y = Variance)) +
  geom_col(fill = "steelblue", alpha = 0.8) +
  geom_line(aes(y = Cumulative, group = 1), color = "orchid", linewidth = 1) +
  geom_point(aes(y = Cumulative), color = "orchid", size = 2) +
  geom_text(aes(y = Variance, label = paste0(Variance, "%")), vjust = -0.4, size = 3) +
  scale_y_continuous(sec.axis = sec_axis(~ ., name = "Cumulative variance (%)")) +
  theme_classic(base_size = 13) +
  labs(title = sprintf("Scree plot – variance explained by PC (top %d variable genes)", ntop_scree),
       x = NULL, y = "Variance (%)")
print(p_scree)
ggsave("Report/PCA_scree.svg", p_scree, width = 8, height = 4, dpi = 300)
} # end run_pca
```

#### 11 PCA Plot (Expressed Genes)

```
if (run_pca) {
vst_obj <- vst(dds, blind = TRUE)

mat <- SummarizedExperiment::assay(vst_obj)
keep_genes <- rowSums(!is.finite(mat)) == 0
vst_clean <- vst_obj[keep_genes, ]

pca_data <- plotPCA(vst_clean, intgroup = "condition", returnData = TRUE, ntop = nrow(vst_clean))
percentVar <- round(100 * attr(pca_data, "percentVar"))

pca_data$label <- paste0(stringr::str_sub(rownames(pca_data), -10), " (", pca_data$condition, ")")

p_pca <- ggplot(pca_data, aes(x = PC1, y = PC2, color = condition, fill = condition)) +
  stat_ellipse(geom = "polygon", alpha = 0.08, level = 0.9, linetype = "dashed") +
  geom_point(size = 4, shape = 21, color = "white", stroke = 0.5,
             aes(fill = condition)) +
  ggrepel::geom_text_repel(aes(label = label), size = 3.2, show.legend = FALSE,
                            box.padding = 0.4, segment.color = "grey60") +
  labs(title = "PCA – all expressed genes",
       x = paste0("PC1: ", percentVar[1], "% variance"),
       y = paste0("PC2: ", percentVar[2], "% variance"),
       color = "Condition", fill = "Condition") +
  theme_classic(base_size = 14) +
  theme(panel.grid.major = element_line(color = "grey90")) +
  coord_equal()

print(p_pca)
ggsave("Report/PCA_plot_TOT.svg", p_pca, dpi = 300, width = 8, height = 7)

} # end run_pca
```

#### 12 Heatmaps

```
if (run_heatmap_variable && run_pca) {
vst_mat <- assay(vst_obj)
gene_vars <- matrixStats::rowVars(vst_mat)
topVarGenes <- head(order(gene_vars, decreasing = TRUE), geni_variabili)
mat_topVar <- vst_mat[topVarGenes, , drop = FALSE]

vst_dist <- dist(t(mat_topVar))
sample_clustering <- hclust(vst_dist)

# Cap z-scores for readability
mat_scaled <- t(scale(t(mat_topVar)))
mat_scaled <- pmax(pmin(mat_scaled, 3), -3)

annotation_col <- as.data.frame(colData(dds)[, "condition", drop = FALSE])

my_cols_heat <- grDevices::colorRampPalette(c("#2166AC", "white", "#B2182B"))(100)

p_heat <- pheatmap(mat_scaled,
         color = my_cols_heat,
         cluster_rows = TRUE,
         cluster_cols = sample_clustering,
         annotation_col = annotation_col,
         show_rownames = FALSE,
         show_colnames = TRUE,
         fontsize_col = 10,
         angle_col = 45,
         treeheight_row = 30,
         treeheight_col = 20,
         border_color = NA,
         main = sprintf("Heatmap of %d most variable genes (z-score, capped ±3)", geni_variabili),
         silent = TRUE)

# Save as SVG via svglite (more reliable than ggsave for pheatmap grobs)
h_in <- max(6, min(30, 0.03 * geni_variabili + 4))
svglite::svglite("Report/Heatmap_top.svg", width = 10, height = h_in)
grid::grid.newpage(); grid::grid.draw(p_heat$gtable)
grDevices::dev.off()
grid::grid.newpage(); grid::grid.draw(p_heat$gtable)
} # end run_heatmap_variable
```

#### 13 Sample Distance Heatmap

```
if (run_heatmap_variable) {
  # Euclidean distance between samples on VST-normalized data (QC)
  vst_mat <- assay(vst_obj)
  gene_vars <- matrixStats::rowVars(vst_mat)
 
  topVarGenes <- head(order(gene_vars, decreasing = TRUE), geni_variabili)

  vst_dist <- dist(t(vst_mat[topVarGenes, , drop = FALSE]))
  dist_mat  <- as.matrix(vst_dist)
  
  anno_row <- data.frame(condition = colData(dds)[, "condition"], row.names = rownames(dist_mat))
  
  sd_cols <- grDevices::colorRampPalette(c( "#B2182B", "white"))(100)
  
  ph_dist <- pheatmap::pheatmap(
    dist_mat,
    color           = rev(sd_cols),
    clustering_distance_rows = vst_dist,
    clustering_distance_cols = vst_dist,
    annotation_row  = anno_row,
    annotation_col  = anno_row,
    border_color    = NA,
    treeheight_row  = 25,
    treeheight_col  = 25,
    main            = "Sample-to-sample Euclidean distance (VST)",
    silent          = TRUE
  )
  
  svglite::svglite("Report/Sample_distance_heatmap.svg", width = 8, height = 7)
  grid::grid.newpage(); grid::grid.draw(ph_dist$gtable)
  grDevices::dev.off()
  grid::grid.newpage(); grid::grid.draw(ph_dist$gtable)
} # end run_heatmap_variable (sample distance)
```

#### 14 Specific Markers

```
if (run_heatmap_markers) {

dir_create("Report/Pathways", recurse = TRUE)


marker_genes <- list(
  H2O2 = c(
    "AT5G59820", # ZAT12
    "AT3G46090", # ZAT7
    "AT1G07890", # APX1
    "AT4G35090", # CAT2
    "AT5G01600", # FER1
    "At2G41730", # HRG1
    "AT5G24640", # HRG2
    "AT1G34190", # ANAC017
    "AT3G26744", # ICE1 (bHLH)
    "AT1G26260", # bHLH TF (H2O2 footprint)
    "AT2G42280", # bHLH TF
    "AT1G69570", # Dof zinc finger TF
    "AT5G15840", # CONSTANS Zn finger TF 
    "AT2G13960", # MYB TF
    "AT2G47190", # MYB2
    "AT3G55730", # MYB109
    "AT1G69490", # NAM TF
    "AT4G11660", # HsfB2B
    "AT2G30250", # WRKY25
    "AT3G25250", # OXI1
    "AT2G26150", # HSFA2
    "AT5G05410", # DREB2A
    "AT4G34410", # RRTF1 (redox responsive TF; ROS/redox inducibile)
    "AT4G18880", # HSFA4A (coinvolto in tolleranza a stress/ossidanti)
    "AT3G04670", # WRKY39 (ROS footprint; often increases under severe oxidative stress)
    "AT4G33950", # OST1/SnRK2.6 (crosstalk ABA–ROS; utile in stress con ROS elevati)
    "AT5G47910"  # RBOHD
  ),
  eSORG = c(
    "AT2G26530",
    "AT4G11280",
    "AT1G61340",
    "AT3G55980",
    "AT4G01250",
    "AT1G80840",
    "AT4G23810",
    "AT1G18570",
    "AT5G04340",
    "AT2G44840",
    "AT1G27730",
    "AT3G46090",
    "AT1G32640",
    "AT1G59590",
    "AT2G40140",
    "AT5G08790",
    "AT2G40000",
    "AT5G62020",
    "AT2G38230",
    "AT5G27420",
    "AT3G05200",
    "AT1G31290",
    "AT2G07681",
    "AT5G18670",
    "AT5G20230",
    "AT3G61190",
    "AT2G41010",
    "AT1G66400",
    "AT5G42380",
    "AT5G26920",
    "AT3G44260",
    "AT4G25470",
    "AT2G45080",
    "AT3G50930",
    "AT5G45340",
    "AT4G13395",
    "AT2G22500",
    "AT4G24570",
    "AT5G24530",
    "AT5G52250",
    "AT1G28370",
    "AT5G61600",
    "AT4G17500",
    "AT5G47220",
    "AT3G15210",
    "AT5G47230",
    "AT4G17490",
    "AT5G58430",
    "AT2G33580",
    "AT1G73500",
    "AT3G18690",
    "AT1G64660",
    "AT3G50060",
    "AT5G06320",
    "AT1G73540",
    "AT1G18300",
    "AT1G79360",
    "AT1G56250",
    "AT1G35140",
    "AT1G06570",
    "AT5G54490",
    "AT1G19770",
    "AT2G38210",
    "AT5G59820",
    "AT3G46620",
    "AT5G59550",
    "AT3G10985",
    "AT3G02040",
    "AT3G56710",
    "AT1G72930",
    "AT2G41100",
    "AT3G16720",
    "AT3G23230",
    "AT2G36800",
    "AT2G38470",
    "AT2G46400",
    "AT3G56400",
    "AT5G57560",
    "AT4G25810",
    "AT2G28400",
    "AT4G29780",
    "AT2G07708",
    "AT1G61470",
    "AT1G69890",
    "AT3G55840",
    "AT2G32210",
    "AT1G22470",
    "AT3G45730",
    "AT2G36220",
    "AT2G07687",
    "AT3G04640",
    "AT1G72920",
    "AT2G32200",
    "AT3G56880",
    "AT1G66090",
    "AT1G21326",
    "AT4G30370",
    "AT5G64870",
    "AT3G50560",
    "AT1G31310",
    "AT2G30020",
    "AT2G22880",
    "AT1G72950",
    "AT1G77640",
    "AT1G64065",
    "AT1G21680",
    "AT1G05575",
    "AT2G07671",
    "AT3G02840",
    "AT5G51190",
    "AT1G15010",
    "AT2G20142",
    "AT4G24380",
    "AT5G17350",
    "AT3G12910",
    "AT5G52750",
    "AT4G27657",
    "AT4G27280",
    "AT3G52800",
    "AT1G76600",
    "AT1G35210",
    "AT1G76590",
    "AT1G27100",
    "AT1G14480",
    "AT2G27080",
    "AT5G41750",
    "AT3G57450",
    "AT5G24640",
    "AT2G32190",
    "AT5G55970",
    "AT5G46710",
    "AT1G19380",
    "AT1G23710",
    "AT4G34150",
    "AT3G12830",
    "AT3G10930",
    "AT4G28703",
    "AT1G59860",
    "AT5G63130",
    "AT3G09870",
    "AT2G39650",
    "AT5G19240",
    "AT2G07698",
    "AT1G72900",
    "AT2G21820",
    "AT4G17250",
    "AT1G30190",
    "AT5G45630",
    "AT1G72940",
    "AT4G24110",
    "AT1G19020",
    "AT3G50800",
    "AT1G33760",
    "AT1G32920",
    "AT5G10695",
    "AT2G32030",
    "AT5G66070",
    "AT1G72910",
    "AT1G32928",
    "AT4G37290",
    "AT5G39670",
    "AT3G29000",
    "AT5G66650",
    "AT1G14200",
    "AT2G04795",
    "AT1G07135",
    "AT2G07727",
    "AT1G76650"
  ),
  ColdStress = c(
  "AT4G25490", # CBF1 / DREB1B
  "AT4G25470", # CBF2 / DREB1C
  "AT4G25480", # CBF3 / DREB1A
  "AT3G50970", # LTI30
  "AT5G52310", # COR15A
  "AT2G42540", # COR15B
  "AT5G15960", # KIN1
  "AT5G15970", # KIN2
  "AT1G20440", # COR47 / RD17
  "AT2G15970", # RD29A / COR78
  "AT5G52300", # RD29B
  "AT1G29395", # ERD10
  "AT1G11760", # MED2 required for CBF
  "AT1G12860", # ICE2 required for CBF in freezing
  "AT3G26744", # ICE1
  "AT4G33950", # OST1 / SnRK2.6 (cold/ABA crosstalk)
  "AT1G76180"  # ERD14
    ),
  HeatStress = c(
    "AT2G29500", # HSFA2
    "AT4G36990", # HSFA1A
    "AT5G16820", # HSFA1B
    "AT3G02990", # HSFA1D
    "AT3G24520", # HSFA1E
    "AT1G74310", # HSP101
    "AT5G59720", # HSP90.1
    "AT3G12580", # HSP70-1
    "AT1G16030", # HSP70-3
    "AT5G52640", # HSP70-4
    "AT3G46230" # HSP17.6A
  ),
  ABA = c(
    "AT5G52300", # RD29B
    "AT5G66400", # RAB18
    "AT5G25610", # RD22
    "AT2G36270", # ABI5 (TF chiave ABA)
    "AT4G26080", # ABI1 / PP2C
    "AT5G67030", # ABA1
    "AT2G42540", # COR15A
    "AT5G15960", # KIN1
    "AT4G33950", # OST1 / SnRK2.6
    "AT1G45249",  # ABF2/AREB1 (TF ABRE-binding; ABA-inducibile)
    "AT3G14440"  # NCED3
  ),
  Jasmonate = c(
    "AT5G44420", # PDF1.2
    "AT5G24770", # VSP2
    "AT3G45140", # LOX2
    "AT1G17420", # LOX3
    "AT5G42650", # AOS
    "AT2G06050", # OPR3
    "AT3G25760", # AOC1
    "AT3G25770", # AOC2
    "AT3G25780", # AOC3
    "AT1G13280", # AOC4
    "AT1G32640", # MYC2
    "AT5G13220", # JAZ10
    "AT2G39940", # COI1
    "AT2G46370"  # JAR1
  ),
  Detox = c(
    "AT3G28740", # CYP81D11 (marker classico della detox; sotto controllo TGAII–SCL14)
    "AT1G54870", # ChlADR (chloroplast aldehyde reductase; gene reporter della via)
    "AT5G16970", # AER (2-alkenal reductase; detox di reactive carbonyls)
    "AT2G37770", # AKR4C9 / ChlAKR (aldo-keto reductase; reporter SCL14-dipendente)
    "AT1G28480", # ROXY19 / GRX480 (CC-type glutaredoxin; legata al network TGA)
    "AT5G61820", # Nod19-like / stress up-regulated Nod19 protein (usato come marker in questo ramo)
    "AT2G47730", # GSTF8 (reporter classico; early stress/chemical response)
    "AT3G61220", # SDR1 (riportato come gene SCL14-dipendente in questo contesto)
    "AT5G63790", # ANAC102
    "AT5G08790", # ATAF2
    "AT1G01720", # ATAF1
     
    # ---- Phase II: GSTU (glutathione S-transferases, tau) ----
  "AT2G29420", # GSTU7
  "AT3G09270", # GSTU8
  "AT1G78380", # GSTU19
  "AT1G78360", # GSTU21
  "AT1G78340", # GSTU22
  "AT1G17170", # GSTU24
  "AT1G17180", # GSTU25
  "AT4G34138", # UGT73B1
  "AT4G34135", # UGT73B2
  "AT4G34131", # UGT73B3
  "AT2G15480", # UGT73B5
  "AT1G30400", # ABCC1 (MRP1)
  "AT2G34660", # ABCC2 (MRP2)
  "AT3G13080", # ABCC3 (MRP3)
  "AT1G59870"  # ABCG36 / PEN3 / PDR8
  ),
  Photosynthesis = c(
 # -------------------------
  # Antenna proteins / PhANGs (molto sensibili a retrograde)
  # -------------------------
  "AT1G29910", # CAB3 / Lhcb1-type (LHCII antenna)  
  "AT1G29920", # CAB2 / Lhcb1.1 (LHCII antenna)     
  "AT1G29930", # CAB1 / Lhcb1-type (LHCII antenna)  
  "AT2G34420", # Lhcb1.5 (cluster chr2)             
  "AT2G34430", # Lhcb1.4 (cluster chr2)             
  "AT2G05100", # LHCB2.1 (LHCII antenna)            
  "AT5G54270", # LHCB3 (LHCII antenna)              
  "AT3G54890", # LHCA1 (LHCI antenna, PSI)          

  # -------------------------
  # Core photosynthesis readouts (PhANGs “classici”)
  # -------------------------
  "AT1G67090", # RBCS1A (Rubisco small subunit)      

  # -------------------------
  # Chlorophyll / antenna biogenesis (often GLK targets)
  # -------------------------
  "AT1G58290", # HEMA1 (tetrapyrrole/chlorophyll start) 
  "AT5G54190", # PORA (protochlorophyllide reductase A) 
  "AT1G44446", # CAO/CH1 (chlorophyll a oxygenase -> Chl b, antenna) 

  # -------------------------
  # Retrograde / nuclei-chloroplast signaling nodes
  # -------------------------
  "AT2G20570", # GLK1 (Golden2-like 1; attiva antenna/clorofilla; retrograde-responsive) 
  "AT5G44190", # GLK2 (Golden2-like 2)                                         
  "AT2G31400", # GUN1 (plastid PPR; biogenic retrograde)                         
  "AT5G13630", # GUN5 / CHLH (Mg-chelatase subunit H; tetrapyrrole/retrograde)   
  "AT2G40220", # ABI4 (TF proposto downstream di GUN1 in retrograde)             

  # -------------------------
  # Light acclimation / state transitions (antenne “dinamiche”)
  # -------------------------
  "AT1G68830", # STN7 (kinase; fosforila LHCII per state transitions) 
  "AT5G01920", # STN8 (kinase; core PSII phosphorylation)            
  "AT4G27800"  # TAP38/PPH1 (fosfatasi LHCII; state transitions)      
  ),
  Salicylate = c(
    "AT2G14610", # PR1
    "AT3G57260", # PR2
    "AT1G75040", # PR5
    "AT1G64280", # NPR1
    "AT1G74710", # ICS1/SID2
    "AT4G39030", # EDS5
    "AT5G13320", # PBS3
    "AT5G67160", # EPS1 a valle di PBS3 nella sintesi
    "AT3G56400"  # WRKY70
  ),
 Mitochondrial_ribosomal_protein = c(
  "AT2G42710", # RPL1
  "AT2G07715", # RPL2A
  "AT2G44065", # RPL2B
  "ATMG00560", # RPL2C
  "ATMG00690", # RPL2D
  "AT3G17465", # RPL3
  "AT2G20060", # RPL4
  "ATMG00210", # RPL5
  "AT5G53070", # RPL9
  "AT3G12370", # RPL10
  "AT4G35490", # RPL11
  "AT1G70190", # RPL12A
  "AT4G36420", # RPL12B
  "AT4G37660", # RPL12C
  "AT3G01790", # RPL13
  "AT1G17560", # HLL
  "AT5G46160", # HLP
  "AT4G23620", # RPL15A
  "AT5G64670", # RPL15B
  "ATMG00080", # RPL16
  "AT5G64650", # RPL17A
  "AT5G09770", # RPL17B
  "AT3G22450", # RPL18A
  "AT3G45020", # RPL18B
  "AT1G08845", # RPL18C
  "AT1G24240", # RPL19A
  "AT4G11630", # RPL19B
  "AT4G16030", # RPL19D
  "AT1G16740", # RPL20
  "AT1G57860", # RPL21F
  "AT4G30930", # NFD1
  "AT1G09690", # RPL21C
  "AT1G52370", # RPL22A
  "AT4G28360", # RPL22B
  "AT1G04480", # RPL23B
  "AT3G04400", # RPL23C
  "AT4G39880", # RPL23A
  "AT5G66860", # RPL25
  "AT2G16930", # RPL27A
  "AT5G15220", # RPL27B
  "AT5G39800", # RPL27C
  "AT1G07830", # RPL29
  "AT5G55140", # RPL30
  "AT1G07070", # RPL35AA
  "AT3G55750", # RPL35AD
  "AT5G40080", # RPL41
  "AT3G59650", # RPL51
  "AT3G03600", # RPS2
  "ATMG00090", # RPS3
  "ATMG00290", # RPS4
  "AT1G64880", # RPS5
  "AT2G07696", # RPS7A
  "ATMG01270", # RPS7B
  "AT3G49080", # RPS9
  "AT3G22300", # RPS10
  "AT1G31817", # NFD3
  "ATMG00980", # RPS12A
  "AT2G07675", # RPS12B
  "AT1G77750", # RPS13
  "AT2G34520", # RPS14
  "AT1G15810", # RPS15A
  "AT1G80620", # RPS15B
  "AT5G56940", # RPS16
  "AT2G05220", # RPS17B
  "AT3G10610", # RPS17C
  "AT5G04800", # RPS17D
  "AT1G07210", # RPS18
  "AT5G47320", # RPS19
  "AT1G16870"  # RPS29
),

Prohibitin = c(
  "AT5G40770", # PHB3
  "AT3G27280", # PHB4
  "AT4G28510", # PHB1
  "AT1G03860", # PHB2
  "AT2G20530"  # PHB6
),

Chaperone = c(
  "AT1G14980", # CPN10
  "AT1G23100", # GroES-like
  "AT3G23990", # HSP60-3B
  "AT2G33210", # HSP60-2
  "AT3G13860", # HSP60-3A
  "AT4G37910", # mtHSC70-1
  "AT5G09590"  # mtHSC70-5
),

Protease_and_peptidase = c(
  "AT5G23140", # CLPP2
  "AT5G26860", # LON1
  "AT3G05790", # LON4
  "AT3G05780", # LON3
  "AT1G07510", # FTSH10
  "AT2G29080", # FTSH3
  "AT2G26140", # FTSH4
  "AT5G53170", # FTSH11
  "AT5G27660", # DegP14
  "AT1G65630", # DegP3
  "AT1G65640", # DegP4
  "AT5G36950", # DegP10
  "AT3G16540", # DegP11
  "AT3G16550", # DegP12
  "AT3G02090", # Mppb
  "AT1G51980", # Mppa-1
  "AT3G16480"  # Mppa-2
),

Protein_transport = c(
  "AT1G20350", # TIM17-1
  "AT2G37410", # TIM17-2
  "AT5G11690"  # TIM17-3
),

Alternative_respiratory_pathway = c(
  "AT3G22370", # AOX1a
  "AT3G22360", # AOX1b
  "AT3G27620", # AOX1c
  "AT1G32350", # AOX1d
  "AT5G64210", # AOX2
  "AT1G07180", # NDA1
  "AT2G29990", # NDA2
  "AT4G28220", # NDB1
  "AT4G05020", # NDB2
  "AT4G21490", # NDB3
  "AT2G20800", # NDB4
  "AT5G08740", # NDC1
  "AT3G54110", # UCP1
  "AT5G58970", # UCP2
  "AT1G14140", # UCP3
  "AT4G24570", # UCP4
  "AT2G22500", # UCP5
  "AT5G09470"  # UCP6
)
)

# ----------------------------
# Build log2FC matrix from df_final
# ----------------------------

# Determine control + treatment order from the sample sheet (same order used in the analysis)
# other_conds and control_cond are defined earlier in the report
logfc_cols <- paste0("log2FC_", other_conds)

# --- Option A: mask non-significant cells using padj columns ---
padj_cols <- paste0("padj_", other_conds)
missing_padj <- setdiff(padj_cols, colnames(df_final))
if (length(missing_padj) > 0) {
  stop("Missing padj columns in df_final: ", paste(missing_padj, collapse = ", "))
}

# ----------------------------
match_geneids <- function(gene_ids, universe) {
  # Recupera TUTTE le righe di df_final associate ai TAIR richiesti.
  # match() restituirebbe invece soltanto la prima riga per ogni TAIR,
  # eliminando di fatto gli omologhi multipli di Physcomitrium.
  requested <- unique(toupper(as.character(gene_ids)))
  universe_upper <- toupper(as.character(universe))

  idx <- unlist(
    lapply(requested, function(id) which(!is.na(universe_upper) & universe_upper == id)),
    use.names = FALSE
  )

  list(idx = idx, ids = universe_upper[idx])
}

plot_marker_heatmap_log2fc <- function(set_name, genes, df_final, logfc_cols,
                                       padj_cols = NULL, padj_cutoff = NULL,
                                       scale_max = 2,
                                       scale_mode = c("cap", "fixed"),
                                       show_in_r = TRUE,
                                       save_png = TRUE,
                                       label_col = "Name") {
  scale_mode <- match.arg(scale_mode)
  stopifnot(all(c("GeneID_X", "GeneID", label_col) %in% colnames(df_final)))

  need_padj <- !is.null(padj_cols) && !is.null(padj_cutoff)
  if (need_padj) stopifnot(all(padj_cols %in% colnames(df_final)))

  m <- match_geneids(genes, df_final$GeneID)
  if (length(m$idx) == 0) {
    message(sprintf("[SKIP] %s: nessun marker trovato in df_final.", set_name))
    return(invisible(NULL))
  }

  cols_take <- c("GeneID_X", "GeneID", label_col,
                 logfc_cols, if (need_padj) padj_cols)
  sub <- df_final[m$idx, cols_take, drop = FALSE]

  # Etichetta univoca: nome/TAIR + gene di Physcomitrium.
  # In questo modo tutti i geni Pp che condividono lo stesso TAIR
  # restano visibili e distinguibili nella heatmap.
  lab <- as.character(sub[[label_col]])
  bad <- is.na(lab) | lab == "" | toupper(lab) == "#N/D"
  lab[bad] <- as.character(sub$GeneID)[bad]
  lab <- paste0(lab, " | ", sub$GeneID_X)
  rownames(sub) <- make.unique(lab)

  # Salva esplicitamente tutte le associazioni marker TAIR - omologhi Pp.
  marker_out <- file.path(
    "Report/Pathways",
    paste0("Marker_genes_", set_name, "_all_Pp_homologs.tsv")
  )
  write.table(sub, file = marker_out, sep = "\t", quote = FALSE,
              row.names = FALSE, col.names = TRUE)

  mat <- as.matrix(sub[, logfc_cols, drop = FALSE])
  storage.mode(mat) <- "numeric"
  mat[!is.finite(mat)] <- NA_real_
  colnames(mat) <- other_conds

  # --- Option A: mask cells with padj >= cutoff (set to NA -> grey) ---
  if (need_padj) {
    padj <- as.matrix(sub[, padj_cols, drop = FALSE])
    storage.mode(padj) <- "numeric"
    padj[!is.finite(padj)] <- NA_real_

    mat[!is.na(padj) & padj >= padj_cutoff] <- NA_real_
  }

  # --- SORT GENES BY THE VALUE OF THE FIRST COLUMN (other_conds[1]) ---
  first_col <- other_conds[1]
  ord <- order(mat[, first_col], decreasing = TRUE, na.last = TRUE)
  mat <- mat[ord, , drop = FALSE]

  maxabs_data <- suppressWarnings(max(abs(mat), na.rm = TRUE))
  if (!is.finite(maxabs_data) || maxabs_data == 0) maxabs_data <- 1
  maxabs <- if (scale_mode == "fixed") scale_max else min(maxabs_data, scale_max)

  mat_clip <- pmax(pmin(mat,  maxabs), -maxabs)
  breaks <- seq(-maxabs, maxabs, length.out = 101)

  main_txt <- paste0(
    "Marker heatmap: ", set_name,
    " (log2FC vs ", control_cond,
    if (need_padj) paste0("; masked padj ≥ ", padj_cutoff) else "",
    "; scale ±", round(maxabs, 2), ")"
  )
  my_cols <- grDevices::colorRampPalette(c("#2166AC", "white", "#B2182B"))(100)

  # ---- CREA L'OGGETTO UNA SOLA VOLTA ----
 ph <- pheatmap::pheatmap(
    mat_clip,
    color = my_cols,
    cluster_rows = FALSE,
    cluster_cols = FALSE,
    show_rownames = TRUE,
    show_colnames = TRUE,
    angle_col = 45,
    fontsize_row = 8,
    fontsize_col = 10,
    cellheight = 14,
    cellwidth = 14,
    main = main_txt,
    breaks = breaks,
    na_col = "grey90",
    silent = FALSE
  )

  # ---- SAVE PNG (senza richiamare pheatmap) ----
  if (isTRUE(save_png)) {
    out_png <- file.path("Report/Pathways", paste0("Heatmap_markers_", set_name, "_log2FC.png"))
    h_in <- max(6, min(40, 0.25 * nrow(mat_clip) + 3))
    w_in <- 8

    grDevices::png(out_png, width = w_in, height = h_in, units = "in", res = 300)
    grid::grid.newpage()
    grid::grid.draw(ph$gtable)
    grDevices::dev.off()
  }

  return(out_png)
}

heatmap_files <- character(0)

# Run heatmaps
for (nm in names(marker_genes)) {
  out_png <- plot_marker_heatmap_log2fc(
    set_name = nm,
    genes = marker_genes[[nm]],
    df_final = df_final,
    logfc_cols = logfc_cols,
    padj_cols = padj_cols,
    padj_cutoff = padj,
    scale_max = 2,
    scale_mode = "cap",
    show_in_r = TRUE,
    save_png = TRUE,
    label_col = "Name"
  )
   heatmap_files <- c(heatmap_files, out_png)
}

knitr::include_graphics(heatmap_files)

marker_correlation_table <- function(set_name, genes, df_final,
                                     other_conds, logfc_cols,
                                     label_col = "Name") {

  stopifnot(all(c("GeneID_X", "GeneID", label_col) %in% colnames(df_final)))

  m <- match_geneids(genes, df_final$GeneID)
  if (length(m$idx) == 0) {
    message(sprintf("[SKIP] %s: nessun marker trovato in df_final.", set_name))
    return(NULL)
  }

  sub <- df_final[m$idx,
                  c("GeneID_X", "GeneID", label_col, logfc_cols),
                  drop = FALSE]

  # label: Name con fallback a GeneID se NA/vuoto/#N/D
  lab <- as.character(sub[[label_col]])
  bad <- is.na(lab) | lab == "" | toupper(lab) == "#N/D"
  lab[bad] <- as.character(sub$GeneID)[bad]
  lab <- paste0(lab, " | ", sub$GeneID_X)
  lab <- make.unique(lab)

  # log2FC matrix (rows=genes, columns=conditions)
  mat <- as.matrix(sub[, logfc_cols, drop = FALSE])
  storage.mode(mat) <- "numeric"
  mat[!is.finite(mat)] <- NA_real_
  colnames(mat) <- other_conds
  rownames(mat) <- lab

  pairs <- combn(other_conds, 2, simplify = FALSE)

  out <- purrr::map_dfr(pairs, function(pr) {
    xcond <- pr[1]; ycond <- pr[2]

    dfp <- data.frame(
      gene = rownames(mat),
      x = mat[, xcond],
      y = mat[, ycond],
      stringsAsFactors = FALSE
    )
    dfp <- dfp[is.finite(dfp$x) & is.finite(dfp$y), , drop = FALSE]

    n <- nrow(dfp)
    if (n < 3) {
      return(tibble::tibble(
        Set = set_name,
        X = xcond, Y = ycond,
        n = n,
        Pearson_r = NA_real_,
        R2 = NA_real_,
        p_value = NA_real_
      ))
    }

    ct <- stats::cor.test(dfp$x, dfp$y, method = "pearson")
    fit <- stats::lm(y ~ x, data = dfp)

    tibble::tibble(
      Set = set_name,
      X = xcond, Y = ycond,
      n = n,
      Pearson_r = unname(ct$estimate),
      R2 = summary(fit)$r.squared,
      p_value = ct$p.value
    )
  })

  message(sprintf("[OK] %s: calcolate correlazioni su %d geni trovati.", set_name, nrow(mat)))
  out
}

# ---- RUN correlations on all categories ----
if (length(other_conds) < 2) {
  message("[SKIP] Correlations: servono >=2 condizioni in 'other_conds'.")
} else {

  corr_all <- purrr::map_dfr(names(marker_genes), function(nm) {
    marker_correlation_table(
      set_name    = nm,
      genes       = marker_genes[[nm]],
      df_final    = df_final,
      other_conds = other_conds,
      logfc_cols  = logfc_cols,
      label_col   = "Name"
    )
  })

  # opzionale: ordina e aggiungi FDR globale per set (o globale totale)
  corr_all <- corr_all %>%
    dplyr::group_by(Set) %>%
    dplyr::mutate(p_adj_BH = p.adjust(p_value, method = "BH")) %>%
    dplyr::ungroup() %>%
    dplyr::arrange(Set, dplyr::desc(abs(Pearson_r)))

  write.table(corr_all,
              file = "Report/Pathways/Marker_correlations.tsv",
              sep = "\t", quote = FALSE, row.names = FALSE)

}

} # end run_heatmap_markers
```

#### 15 Organelles

```
if (run_organelles) {

# ------------------------------------------------------------
# Organelle enrichment (SUBA localization) with UP/DOWN direction
# Fisher exact test vs background and mirror plot
# ------------------------------------------------------------

dir_create("Report/Organelle", recurse = TRUE)

# --- Parameters ---
padj_cutoff <- padj             
logfc_cutoff <- log2foldchange   

# TPM columns used to define "expressed" background
tpm_cols <- paste0("TPM_", c(control_cond, other_conds))
stopifnot(all(tpm_cols %in% colnames(df_final)))

# Contrast-specific DE columns
padj_cols  <- grep("^padj_",  colnames(df_final), value = TRUE)
logfc_cols <- grep("^log2FC_", colnames(df_final), value = TRUE)

# Ensure padj_* and log2FC_* columns match by contrast name
contrast_names_p <- sub("^padj_", "", padj_cols)
contrast_names_l <- sub("^log2FC_", "", logfc_cols)
idx <- match(contrast_names_p, contrast_names_l)
if (anyNA(idx)) stop("padj_* and log2FC_* columns do not match by contrast name.")
logfc_cols <- logfc_cols[idx]

# --- Build localization master table (exploded) ---
loc_df_master <- df_final %>%
  dplyr::select(GeneID, SUBA_Localization, all_of(tpm_cols)) %>%
  dplyr::filter(!is.na(SUBA_Localization),
                dplyr::if_any(all_of(tpm_cols), ~ . > 0)) %>%
  tidyr::separate_rows(SUBA_Localization, sep = ";|,") %>%
  dplyr::mutate(SUBA_Localization = trimws(SUBA_Localization))

organelle_list <- sort(unique(loc_df_master$SUBA_Localization))

# ------------------------------------------------------------
# Enrichment function for one contrast (UP and DOWN)
# ------------------------------------------------------------
calc_enrichment_by_direction <- function(padj_col, logfc_col, df, loc_df_master, organelle_list,
                                         tpm_cols, padj_cutoff, logfc_cutoff) {

  # Define the tested/background universe: expressed genes with localization and non-NA stats
  df_expressed <- df %>%
    dplyr::filter(
      rowSums(df[, tpm_cols, drop = FALSE] > 0, na.rm = TRUE) > 0,
      !is.na(SUBA_Localization),
      !is.na(.data[[padj_col]]),
      !is.na(.data[[logfc_col]])
    )

  all_genes <- unique(df_expressed$GeneID)
  total_genes <- length(all_genes)

  # Define DEGs within the same universe
  deg_up <- unique(df_expressed$GeneID[
    df_expressed[[padj_col]] < padj_cutoff & df_expressed[[logfc_col]] >  logfc_cutoff
  ])
  deg_down <- unique(df_expressed$GeneID[
    df_expressed[[padj_col]] < padj_cutoff & df_expressed[[logfc_col]] < -logfc_cutoff
  ])

  n_up <- length(deg_up)
  n_down <- length(deg_down)

  fisher_test_safe <- function(a, b, c, d) {
    mat <- matrix(c(a, b, c, d), nrow = 2)
    out <- tryCatch(stats::fisher.test(mat), error = function(e) NULL)
    list(
      OR = if (!is.null(out)) unname(out$estimate) else NA_real_,
      pval = if (!is.null(out)) out$p.value else NA_real_
    )
  }

  purrr::map_dfr(organelle_list, function(org) {

    # Organelle gene set restricted to the same universe
    genes_in_org <- unique(loc_df_master$GeneID[loc_df_master$SUBA_Localization == org])
    genes_in_org <- intersect(genes_in_org, all_genes)
    total_in_org <- length(genes_in_org)

    # UP contingency table
    a_up <- sum(deg_up %in% genes_in_org)
    b_up <- n_up - a_up
    nondeg_up <- setdiff(all_genes, deg_up)
    c_up <- sum(nondeg_up %in% genes_in_org)
    d_up <- length(nondeg_up) - c_up
    f_up <- fisher_test_safe(a_up, b_up, c_up, d_up)

    # DOWN contingency table
    a_down <- sum(deg_down %in% genes_in_org)
    b_down <- n_down - a_down
    nondeg_down <- setdiff(all_genes, deg_down)
    c_down <- sum(nondeg_down %in% genes_in_org)
    d_down <- length(nondeg_down) - c_down
    f_down <- fisher_test_safe(a_down, b_down, c_down, d_down)

    tibble::tibble(
      Contrast = padj_col,
      Organelle = org,
      Tot_in_Org = total_in_org,
      Tot_Genes = total_genes,
      Tot_UP = n_up,
      Tot_DOWN = n_down,
      UP_in_Org = a_up,
      DOWN_in_Org = a_down,
      OR_UP = f_up$OR,
      OR_DOWN = f_down$OR,
      Fisher_p_UP = f_up$pval,
      Fisher_p_DOWN = f_down$pval,
      Prop_UP = ifelse(n_up > 0, a_up / n_up, NA_real_),
      Prop_DOWN = ifelse(n_down > 0, a_down / n_down, NA_real_),
      Prop_Background = ifelse(total_genes > 0, total_in_org / total_genes, NA_real_)
    )
  }) %>%
    dplyr::mutate(
      Fisher_p_UP_adj = p.adjust(Fisher_p_UP, method = "BH"),
      Fisher_p_DOWN_adj = p.adjust(Fisher_p_DOWN, method = "BH")
    )
}

# ------------------------------------------------------------
# Run enrichment for all contrasts
# ------------------------------------------------------------
all_results <- purrr::map2(
  padj_cols, logfc_cols,
  ~ calc_enrichment_by_direction(.x, .y, df_final, loc_df_master, organelle_list,
                                 tpm_cols, padj_cutoff, logfc_cutoff)
)
all_results_df <- dplyr::bind_rows(all_results)

# Save table
write.csv(all_results_df,
          "Report/Organelle/Organelle_enrichment_by_direction.csv",
          row.names = FALSE)

# ------------------------------------------------------------
# Plotting (mirror plot)
# ------------------------------------------------------------
org_order <- all_results_df %>%
  dplyr::group_by(Organelle) %>%
  dplyr::summarise(bg_mean = mean(Prop_Background, na.rm = TRUE), .groups = "drop") %>%
  dplyr::arrange(desc(bg_mean)) %>%
  dplyr::pull(Organelle)

for (cond in unique(all_results_df$Contrast)) {

  df_c <- subset(all_results_df, Contrast == cond)
  cond_name <- sub("^padj_", "", cond)

  # Keep only the renaming we actually need for pivot_longer pattern
  df_c2 <- df_c %>%
    dplyr::rename(
      Fisher_UP = Fisher_p_UP_adj,
      Fisher_DOWN = Fisher_p_DOWN_adj
    )

  # Long format (Prop / OR / Fisher with Direction UP/DOWN)
  df_long <- df_c2 %>%
    tidyr::pivot_longer(
      cols = c(Prop_UP, Prop_DOWN, OR_UP, OR_DOWN, Fisher_UP, Fisher_DOWN),
      names_to = c(".value", "Direction"),
      names_pattern = "(.*)_(UP|DOWN)"
    )

  # Prepare coordinates and labels
  df_long <- df_long %>%
    dplyr::mutate(
      Organelle = factor(Organelle, levels = org_order),
      ynum = as.numeric(Organelle),

      Prop = as.numeric(Prop),
      Prop_Background = as.numeric(Prop_Background),

      delta = Prop - Prop_Background,
      dir_color = dplyr::case_when(
        delta > 0 ~ "enriched",
        delta < 0 ~ "depleted",
        TRUE ~ "equal"
      ),

      xmin_obs = ifelse(Direction == "DOWN", -Prop_Background,  Prop_Background),
      xmax_obs = ifelse(Direction == "DOWN", -Prop,            Prop),
      xmin_exp = ifelse(Direction == "DOWN", -Prop_Background, 0),
      xmax_exp = ifelse(Direction == "DOWN", 0,                Prop_Background),

      signif_symbol = ifelse(!is.na(Fisher) & Fisher < padj_cutoff, "*", ""),
      signif_flag = ifelse(!is.na(Fisher) & Fisher < padj_cutoff, "signif", "nonsignif"),
      alpha_val = ifelse(signif_flag == "signif", 1, 0.5),

      offset = 0.02 * max(abs(c(Prop, Prop_Background)), na.rm = TRUE),
      label_x = dplyr::case_when(
        dir_color == "enriched" ~ xmax_obs + offset,
        dir_color == "depleted" ~ xmin_obs - offset,
        TRUE ~ xmax_obs + offset
      ),
      hjust_val = ifelse(label_x >= 0, 0, 1)
    )

  # X limits
  xlim_val <- max(abs(df_long$Prop), abs(df_long$Prop_Background), na.rm = TRUE) * 1.4

  # Plot
  p <- ggplot(df_long, aes(y = Organelle)) +

    # Expected background (grey)
    geom_rect(aes(
      xmin = xmin_exp, xmax = xmax_exp,
      ymin = ynum - 0.3, ymax = ynum + 0.3,
      fill = factor("Expected")
    ), alpha = 0.1, color = NA) +

    # Observed (enriched/depleted/equal)
    geom_rect(aes(
      xmin = xmin_obs, xmax = xmax_obs,
      ymin = ynum - 0.3, ymax = ynum + 0.3,
      fill = dir_color,
      alpha = alpha_val
    ), color = NA) +

    # Central zero line
    geom_vline(xintercept = 0, color = "black", linetype = "dashed", linewidth = 0.4) +

    # Significance mark
    geom_text(aes(x = label_x, label = signif_symbol, hjust = hjust_val),
              size = 5, fontface = "bold", na.rm = TRUE) +

    # Fill palette (include 'equal')
    scale_fill_manual(
      values = c(
        "enriched" = "orchid",
        "depleted" = "skyblue",
        "equal" = "grey80",
        "Expected" = "grey50"
      ),
      name = "Type / Direction"
    ) +
    scale_alpha_identity() +

    scale_x_continuous(
      labels = function(x) scales::percent(abs(x), accuracy = 1),
      limits = c(-xlim_val, xlim_val)
    ) +

    labs(
      title = paste0("Organelle enrichment (DOWN & UP regulated): ", cond_name),
      x = "(DOWN) Proportion of DEGs vs Background (UP)",
      y = "Organelle"
    ) +
    theme_minimal() +
    theme(
      panel.grid.minor = element_blank(),
      axis.text.y = element_text(size = 9)
    )

  print(p)
  ggsave(
    filename = file.path("Report/Organelle", paste0("Enrichment_Mirror_", cond_name, ".svg")),
    plot = p, width = 10, height = 6, device = "svg"
  )
}


} # end run_organelles
```

#### 16 MA Plots

```
if (run_volcano) {
  dir_create("Report/MA", recurse = TRUE)
  for (pair_id in names(res_list)) {
    res_df <- res_list[[pair_id]]
    res_df$gene_id <- rownames(res_df)
    res_df_ma <- res_df[!is.na(res_df$baseMean) & !is.na(res_df$log2FoldChange) & res_df$baseMean > 0, ]
    res_df_ma$log2FoldChange_capped <- pmax(pmin(res_df_ma$log2FoldChange, 10), -10)
    res_df_ma$sig <- ifelse(!is.na(res_df_ma$padj) & res_df_ma$padj < padj &
                              abs(res_df_ma$log2FoldChange) > log2foldchange, "sig", "ns")

    up_n   <- sum(res_df_ma$sig == "sig" & res_df_ma$log2FoldChange > log2foldchange, na.rm = TRUE)
    down_n <- sum(res_df_ma$sig == "sig" & res_df_ma$log2FoldChange < -log2foldchange, na.rm = TRUE)

    res_df_ma$color <- "ns"
    res_df_ma$color[res_df_ma$sig == "sig" & res_df_ma$log2FoldChange >  0] <- "up"
    res_df_ma$color[res_df_ma$sig == "sig" & res_df_ma$log2FoldChange <  0] <- "down"

    p_ma <- ggplot(res_df_ma, aes(x = log10(baseMean + 1), y = log2FoldChange_capped, color = color)) +
      geom_point(alpha = 0.5, size = 0.8) +
      geom_hline(yintercept = c(-log2foldchange, log2foldchange),
                 linetype = "dashed", color = "grey50", linewidth = 0.4) +
      geom_hline(yintercept = 0, color = "black", linewidth = 0.3) +
      scale_color_manual(
        values = c("up" = "orchid", "down" = "skyblue", "ns" = "grey70"),
        breaks = c("up", "down"),
        labels = c("up" = paste0("Up (", up_n, ")"), "down" = paste0("Down (", down_n, ")")),
        name = NULL
      ) +
      labs(title = paste("MA plot:", pair_id),
           x = "log10(mean normalised count + 1)",
           y = "log2 Fold Change (capped ±10)") +
      theme_classic(base_size = 13) +
      theme(panel.grid.major = element_line(color = "grey92"))
    print(p_ma)
    ggsave(paste0("Report/MA/MA_", pair_id, ".svg"), p_ma, width = 7, height = 5, dpi = 300)
  }
} # end MA plots
```

#### 17 Volcano Plot

```
if (run_volcano) {
dir_create("Report/volcano")

# Volcano Plot function
for (pair_id in names(res_list)) {
  
  res_df <- res_list[[pair_id]]
  
  # ASS ID and symbols
  res_df$gene_id <- rownames(res_df)
  
  # Cap values
  res_df$log2FoldChange_capped <- pmax(pmin(res_df$log2FoldChange, 10), -10)
  res_df$neglog10pval_capped <- pmin(-log10(res_df$padj), 100)
  
  # Filter NA 
  res_df_plot <- subset(res_df, 
                        !is.na(log2FoldChange_capped) & 
                        !is.na(neglog10pval_capped) &
                        !is.na(padj))
  
  # Set colors
  res_df_plot$color <- "ns"
  res_df_plot$color[res_df_plot$padj < padj & res_df_plot$log2FoldChange > log2foldchange]  <- "up"
  res_df_plot$color[res_df_plot$padj < padj & res_df_plot$log2FoldChange < -log2foldchange] <- "down"
  
  # Add DEG number to legend
  up_count   <- sum(res_df_plot$color == "up", na.rm = TRUE)
  down_count <- sum(res_df_plot$color == "down", na.rm = TRUE)
  color_labels <- c(
    up   = paste0("Up (", up_count, ")"),
    down = paste0("Down (", down_count, ")")
  )
  
  padj_line <- -log10(padj)

  # Volcano plot
  volcano_plot <- ggplot(res_df_plot, aes(x = log2FoldChange_capped,
                                          y = neglog10pval_capped)) +
    geom_point(aes(color = color, size = color), alpha = 0.7) +
    scale_size_manual(values = c("up" = 1.8, "down" = 1.8, "ns" = 0.9), guide = "none") +
    # Threshold lines
    geom_hline(yintercept = padj_line, linetype = "dashed", color = "grey50", linewidth = 0.4) +
    geom_vline(xintercept = c(-log2foldchange, log2foldchange),
               linetype = "dashed", color = "grey50", linewidth = 0.4) +
    scale_color_manual(
      values = c("up" = "orchid", "down" = "skyblue", "ns" = "grey75"),
      breaks = c("up", "down"),
      labels = color_labels,
      name = "Differential Expression"
    ) +
    labs(title = paste("Volcano:", pair_id),
         x = "log2 Fold Change (capped ±10)",
         y = "-log10(p-value) (capped 100)") +
    theme_classic(base_size = 14) +
    theme(panel.grid.major = element_line(color = "grey90"))
  
  print(volcano_plot)
  
  # Save plot
  ggsave(paste0("Report/volcano/Volcano_", pair_id, ".svg"), volcano_plot, dpi = 300, width = 7, height = 5)
}
} # end run_volcano
```

#### 18 Venn Diagram: look only at gene directions

```
if (run_venn) {

# Select only "vs_control"
control_comparisons <- grep(paste0("_vs_", control_cond, "$"), names(res_list), value = TRUE)

if (length(control_comparisons) < 2) {
  message("Not enough comparisons to generate Venn diagrams. Skipping this section.")
} else {

  dir_create("Report/venn", recurse = TRUE)

  deg_lists <- list()

  # Extract UP/DOWN lists (use GeneID, never symbols)
  for (comp in control_comparisons) {

    res <- res_list[[comp]]

    # Robust gene id
    #if ("GeneID" %in% colnames(res)) {
    #  res$gene_id <- as.character(res$GeneID)
    #} else {
    #  res$gene_id <- rownames(res)
    #}
    res$gene_id <- rownames(res)

    # Keep only rows with valid stats
    res <- res[!is.na(res$padj) & !is.na(res$log2FoldChange), , drop = FALSE]

    cond <- sub(paste0("_vs_", control_cond, "$"), "", comp)

    deg_lists[[paste0(cond, "_UP")]] <- unique(res$gene_id[
      res$padj < padj & res$log2FoldChange >  log2foldchange
    ])

    deg_lists[[paste0(cond, "_DOWN")]] <- unique(res$gene_id[
      res$padj < padj & res$log2FoldChange < -log2foldchange
    ])
  }

  # Conditions involved (unique cond names)
  cond_names <- unique(sub(paste0("_vs_", control_cond, "$"), "", control_comparisons))

  # All pairwise combinations of conditions
  comb_pairs <- combn(cond_names, 2, simplify = FALSE)

  for (pair in comb_pairs) {

    cond1 <- pair[1]
    cond2 <- pair[2]

    set_names <- c(
      paste0(cond1, "_UP"),
      paste0(cond1, "_DOWN"),
      paste0(cond2, "_UP"),
      paste0(cond2, "_DOWN")
    )

    venn_sets <- list(
      deg_lists[[set_names[1]]],
      deg_lists[[set_names[2]]],
      deg_lists[[set_names[3]]],
      deg_lists[[set_names[4]]]
    )
    names(venn_sets) <- set_names

    # Overlaps (explicit namespace)
    overlaps <- VennDiagram::calculate.overlap(venn_sets)

    # Safer region names: include region index + size
    region_sizes <- vapply(overlaps, length, integer(1))
    region_names <- paste0("Region", seq_along(overlaps), "_nGenes_", region_sizes)
    names(overlaps) <- region_names

    # Make a rectangular table of genes per region
    max_len <- max(region_sizes)
    overlaps_df <- as.data.frame(lapply(overlaps, function(v) {
      c(v, rep("", max_len - length(v)))
    }), stringsAsFactors = FALSE)
    colnames(overlaps_df) <- names(overlaps)

    # Save TSV of region genes
    out_file_genes <- file.path("Report/venn", paste0("genes_", cond1, "_vs_", cond2, ".tsv"))
    write.table(overlaps_df, file = out_file_genes, sep = "\t",
                quote = FALSE, row.names = FALSE, col.names = TRUE)

    # Draw Venn diagram
    venn_plot <- VennDiagram::venn.diagram(
      x = venn_sets,
      filename = NULL,
      fill = c("#9ecae1", "#08519c", "#fcbba1", "#a50f15"),  # fixed order = set_names
      alpha = 0.5,
      cex = 1.2,
      cat.cex = 1.2,
      cat.pos = 0,
      cat.dist = 0.1,
      margin = 0.1,
      main = paste0("Venn Diagram: ", cond1, " vs ", cond2)
    )
    
    # Show in RStudio
    grid::grid.newpage()
    grid::grid.draw(venn_plot)
   
     # Save as SVG
    svg(file.path("Report/venn", paste0("Venn_", cond1, "_vs_", cond2, ".svg")), width = 8, height = 8)
    grid::grid.newpage()
    grid::grid.draw(venn_plot)
    dev.off()
  }

  # Remove VennDiagram log files if present
  suppressWarnings(file.remove(list.files(pattern = "Venn.*\\.log$")))
}


} # end run_venn
```

```
## [1] TRUE TRUE TRUE
```

#### 19 GO Enrichment

```
if (run_go) {
dir_create("Report/GO/GO_results", recurse = TRUE)
dir_create("Report/GO/GO_results_simplified", recurse = TRUE)
dir_create("Report/GO/gene_lists", recurse = TRUE)

min_genes <- 10

for (cond in names(res_list)) {

  res_df <- res_list[[cond]]

  # Tieni solo righe con GeneID, log2FC, padj non-NA
  res_df <- res_df[complete.cases(res_df[, c("GeneID", "log2FoldChange", "padj")]), ]

  # Gene universe (TAIR IDs) + unique
  expressed_genes <- unique(res_df$GeneID)
  expressed_genes <- expressed_genes[!is.na(expressed_genes)]

  # UP/DOWN (TAIR IDs) + unique
  up_genes   <- unique(res_df$GeneID[res_df$padj < padj_cutoff & res_df$log2FoldChange > 0])
  down_genes <- unique(res_df$GeneID[res_df$padj < padj_cutoff & res_df$log2FoldChange < 0])

  # Se entrambi sotto soglia, salta
  if (length(up_genes) < min_genes && length(down_genes) < min_genes) next

  # Save gene lists
  if (length(up_genes) >= min_genes)
    write.table(data.frame(GeneID = up_genes),
                file = file.path("Report/GO/gene_lists", paste0("genes_", cond, "_UP.tsv")),
                sep = "\t", quote = FALSE, row.names = FALSE)

  if (length(down_genes) >= min_genes)
    write.table(data.frame(GeneID = down_genes),
                file = file.path("Report/GO/gene_lists", paste0("genes_", cond, "_DOWN.tsv")),
                sep = "\t", quote = FALSE, row.names = FALSE)

  # Enrichment GO
  ego_up <- if (length(up_genes) >= min_genes)
    enrichGO(gene = up_genes, OrgDb = org.At.tair.db,
             keyType = "TAIR", ont = "BP",
             pAdjustMethod = "BH", qvalueCutoff = 0.05,
             universe = expressed_genes,
             readable = TRUE) else NULL

  ego_down <- if (length(down_genes) >= min_genes)
    enrichGO(gene = down_genes, OrgDb = org.At.tair.db,
             keyType = "TAIR", ont = "BP",
             pAdjustMethod = "BH", qvalueCutoff = 0.05,
             universe = expressed_genes,
             readable = TRUE) else NULL

  # Save every significant term before semantic redundancy reduction.
  ego_up_full <- ego_up
  ego_down_full <- ego_down

  if (!is.null(ego_up_full))
    write.table(as.data.frame(ego_up_full),
                file = file.path("Report/GO/GO_results", paste0("GO_", cond, "_UP.tsv")),
                sep = "\t", quote = FALSE, row.names = FALSE)

  if (!is.null(ego_down_full))
    write.table(as.data.frame(ego_down_full),
                file = file.path("Report/GO/GO_results", paste0("GO_", cond, "_DOWN.tsv")),
                sep = "\t", quote = FALSE, row.names = FALSE)

  # Simplify only the copies used for plots and reduced result tables.
  if (!is.null(ego_up))   ego_up   <- clusterProfiler::simplify(ego_up, cutoff = cutoff_val, by = "p.adjust",
                                                                select_fun = min, measure = "Wang")
  if (!is.null(ego_down)) ego_down <- clusterProfiler::simplify(ego_down, cutoff = cutoff_val, by = "p.adjust",
                                                                select_fun = min, measure = "Wang")

  # Save redundancy-reduced terms separately.
  if (!is.null(ego_up))
    write.table(as.data.frame(ego_up),
                file = file.path("Report/GO/GO_results_simplified", paste0("GO_", cond, "_UP_simplified.tsv")),
                sep = "\t", quote = FALSE, row.names = FALSE)

  if (!is.null(ego_down))
    write.table(as.data.frame(ego_down),
                file = file.path("Report/GO/GO_results_simplified", paste0("GO_", cond, "_DOWN_simplified.tsv")),
                sep = "\t", quote = FALSE, row.names = FALSE)

  # Combine UP/DOWN
  top_up   <- if (!is.null(ego_up))   as.data.frame(ego_up)   else NULL
  top_down <- if (!is.null(ego_down)) as.data.frame(ego_down) else NULL

  top_up   <- if (!is.null(top_up)   && nrow(top_up) > 0)   { top_up$Regulation <- "UP";   top_up }   else NULL
  top_down <- if (!is.null(top_down) && nrow(top_down) > 0) { top_down$Regulation <- "DOWN"; top_down } else NULL

  go_all <- do.call(rbind, Filter(Negate(is.null), list(top_up, top_down)))
  if (is.null(go_all) || nrow(go_all) == 0) next

  # Top 10 most significant (by adjusted p)
  go_top10 <- go_all %>%
    dplyr::mutate(logp = -log10(p.adjust)) %>%
    dplyr::arrange(dplyr::desc(logp)) %>%
    utils::head(10)

  cap <- stats::quantile(go_top10$logp, 0.9, na.rm = TRUE)
  logp_capped <- pmin(go_top10$logp, cap)
  go_top10$Zscore <- ifelse(go_top10$Regulation == "UP", logp_capped, -logp_capped)

  go_plot <- ggplot(go_top10, aes(
    x = Zscore,
    y = reorder(Description, Zscore),
    size = Count,
    color = logp
  )) +
    geom_vline(xintercept = 0, linetype = "dashed", color = "grey60", linewidth = 0.6) +
    geom_point() +
    labs(title = paste("Top 10 GO terms -", cond),
         x = "GO enrichment Z-score",
         y = "GO term",
         size = "Gene Count",
         color = "-log10(adj. p)") +
    scale_color_gradient(low = "skyblue", high = "orchid", limits = c(0, cap), oob = scales::squish) +
    theme_classic(base_size = 14)

  print(go_plot)
  ggsave(filename = paste0("Report/GO/GO_top10_", cond, ".svg"),
         plot = go_plot, width = 10, height = 6, dpi = 300)
}
} # end run_go
```

#### 20 GO Enrichment — *Physcomitrium patens*

This module performs over-representation analysis directly on
*Physcomitrium* gene identifiers (for example,
`Pp3c10_24280V3`), without converting them to TAIR
identifiers. The annotation file must contain the columns
`term`, `gene`, `name`, and
`domain`, and may be tab-, comma-, or
semicolon-delimited.

```
if (run_go_physco) {

  # Also look in db_dir and accept common extensions when the default name is used.
  
  physcomitrium_go_file <- file.path(db_dir, "physcomitrium_GO.tsv")

    # read_delim(delim = NULL) detects tab, comma, or semicolon delimiters.
    physco_go <- readr::read_delim(
      physcomitrium_go_file,
      delim = NULL,
      trim_ws = TRUE,
      show_col_types = FALSE,
      progress = FALSE
    )
    names(physco_go) <- tolower(trimws(names(physco_go)))

    normalise_physco_id <- function(x) {
      toupper(trimws(as.character(x)))
    }

    physco_go <- physco_go %>%
      dplyr::transmute(
        term = trimws(as.character(term)),
        gene = normalise_physco_id(gene),
        name = trimws(as.character(name)),
        domain = tolower(trimws(as.character(domain)))
      ) %>%
      dplyr::filter(
        !is.na(term), nzchar(term), grepl("^GO:[0-9]{7}$", term),
        !is.na(gene), nzchar(gene),
        !is.na(name), nzchar(name)
      ) %>%
      dplyr::distinct(term, gene, .keep_all = TRUE)

    if (nrow(physco_go) == 0) {
      stop("No valid GO-gene associations found in ", physcomitrium_go_file)
    }
    # Do not rely on the domain column supplied by the custom file. Recover the
    # official ontology (BP/MF/CC) for every GO ID from GO.db. This prevents CC
    # terms such as chloroplast/plastid from entering a BP enrichment.
    physco_go_ontology <- suppressMessages(
      AnnotationDbi::select(
        GO.db::GO.db,
        keys = unique(physco_go$term),
        columns = "ONTOLOGY",
        keytype = "GOID"
      )
    ) %>%
      dplyr::distinct(GOID, .keep_all = TRUE)

    physco_go <- physco_go %>%
      dplyr::rename(domain_from_file = domain) %>%
      dplyr::left_join(
        physco_go_ontology,
        by = c("term" = "GOID")
      ) %>%
      dplyr::filter(ONTOLOGY == "BP") %>%
      dplyr::mutate(domain = "biological_process") %>%
      dplyr::select(term, gene, name, domain)

    if (nrow(physco_go) == 0) {
      stop(
        "No Biological Process GO IDs found in ", physcomitrium_go_file,
        " after ontology validation with GO.db."
      )
    }
    
    # One term name/domain per GO ID; duplicated gene-term associations are removed.
    physco_term_meta <- physco_go %>%
      dplyr::group_by(term) %>%
      dplyr::summarise(
        name = dplyr::first(name),
        domain = paste(sort(unique(domain[nzchar(domain)])), collapse = ";"),
        .groups = "drop"
      )

    physco_term2gene <- physco_go %>%
      dplyr::select(term, gene) %>%
      dplyr::distinct()

    physco_term2name <- physco_term_meta %>%
      dplyr::select(term, name)

    annotated_physco_genes <- unique(physco_term2gene$gene)

    fs::dir_create("Report/GO_Physcomitrium/GO_results", recurse = TRUE)
    fs::dir_create("Report/GO_Physcomitrium/GO_results_simplified", recurse = TRUE)
    fs::dir_create("Report/GO_Physcomitrium/gene_lists", recurse = TRUE)

    # Check identifier compatibility before running enrichment. This catches, for
    # example, gene IDs ending in V3 in one file and transcript IDs ending in
    # V3.1 in the other.
    result_physco_genes <- unique(unlist(lapply(
      res_list,
      function(x) normalise_physco_id(rownames(x))
    )))
    matched_physco_genes <- intersect(result_physco_genes, annotated_physco_genes)

    id_matching_summary <- data.frame(
      result_ids = length(result_physco_genes),
      annotation_ids = length(annotated_physco_genes),
      matched_ids = length(matched_physco_genes),
      result_match_percent = round(
        100 * length(matched_physco_genes) / max(1, length(result_physco_genes)),
        2
      ),
      annotation_match_percent = round(
        100 * length(matched_physco_genes) / max(1, length(annotated_physco_genes)),
        2
      )
    )
    readr::write_tsv(
      id_matching_summary,
      "Report/GO_Physcomitrium/id_matching_summary.tsv"
    )

    if (length(matched_physco_genes) == 0) {
      result_examples <- paste(utils::head(result_physco_genes, 5), collapse = ", ")
      annotation_examples <- paste(utils::head(annotated_physco_genes, 5), collapse = ", ")
      stop(
        "No Physcomitrium IDs match between DESeq2 results and GO annotation. ",
        "Examples from DESeq2: ", result_examples, ". ",
        "Examples from GO file: ", annotation_examples, ". ",
        "Check genome release and identifier type (gene versus transcript)."
      )
    }

    # Record annotation size and per-contrast coverage for quality control.
    physco_go_coverage <- list()
    min_deg_genes_physco <- 10

    add_physco_domain <- function(enrichment) {
      if (is.null(enrichment)) return(NULL)
      out <- as.data.frame(enrichment)
      if (nrow(out) == 0) return(NULL)
      out %>%
        dplyr::left_join(
          physco_term_meta %>% dplyr::select(term, domain),
          by = c("ID" = "term")
        )
    }

    for (cond in names(res_list)) {
      res_df <- res_list[[cond]]
      res_df$PhyscoGene <- normalise_physco_id(rownames(res_df))
      res_df <- res_df[
        complete.cases(res_df[, c("PhyscoGene", "log2FoldChange", "padj")]),
        ,
        drop = FALSE
      ]

      expressed_physco <- intersect(
        unique(res_df$PhyscoGene),
        annotated_physco_genes
      )
      up_physco <- intersect(
        unique(res_df$PhyscoGene[
          res_df$padj < padj_cutoff & res_df$log2FoldChange > 0
        ]),
        annotated_physco_genes
      )
      down_physco <- intersect(
        unique(res_df$PhyscoGene[
          res_df$padj < padj_cutoff & res_df$log2FoldChange < 0
        ]),
        annotated_physco_genes
      )

      physco_go_coverage[[cond]] <- data.frame(
        contrast = cond,
        tested_genes = nrow(res_df),
        annotated_tested_genes = length(expressed_physco),
        significant_up = sum(res_df$padj < padj_cutoff & res_df$log2FoldChange > 0),
        annotated_significant_up = length(up_physco),
        significant_down = sum(res_df$padj < padj_cutoff & res_df$log2FoldChange < 0),
        annotated_significant_down = length(down_physco)
      )

      if (length(up_physco) >= min_deg_genes_physco) {
        readr::write_tsv(
          data.frame(GeneID = up_physco),
          file.path("Report/GO_Physcomitrium/gene_lists", paste0("genes_", cond, "_UP.tsv"))
        )
      }
      if (length(down_physco) >= min_deg_genes_physco) {
        readr::write_tsv(
          data.frame(GeneID = down_physco),
          file.path("Report/GO_Physcomitrium/gene_lists", paste0("genes_", cond, "_DOWN.tsv"))
        )
      }

      if (length(expressed_physco) < min_deg_genes_physco ||
          (length(up_physco) < min_deg_genes_physco &&
           length(down_physco) < min_deg_genes_physco)) next

      run_physco_enricher <- function(genes) {
        if (length(genes) < min_deg_genes_physco) return(NULL)
        enrichment <- clusterProfiler::enricher(
          gene = genes,
          universe = expressed_physco,
          TERM2GENE = physco_term2gene,
          TERM2NAME = physco_term2name,
          pAdjustMethod = "BH",
          pvalueCutoff = 0.05,
          qvalueCutoff = 0.05,
          minGSSize = 5,
          maxGSSize = 5000
        )

        if (is.null(enrichment) || nrow(as.data.frame(enrichment)) == 0) {
          return(NULL)
        }

        # enricher() marks custom annotations as UNKNOWN. These are genuine BP
        # GO IDs, so set the ontology explicitly before semantic simplification.
        enrichment@ontology <- "BP"
        enrichment_simplified <- tryCatch(
          clusterProfiler::simplify(
            enrichment,
            cutoff = cutoff_val,
            by = "p.adjust",
            select_fun = min,
            measure = "Wang"
          ),
          error = function(e) {
            warning(
              "Physcomitrium GO simplify failed; using unsimplified terms: ",
              conditionMessage(e)
            )
            enrichment
          }
        )

        # Keep both objects: the complete significant result is written to the
        # result tables, while the simplified result is used only for plotting.
        list(
          full = enrichment,
          simplified = enrichment_simplified
        )
      }

      ego_physco_up <- run_physco_enricher(up_physco)
      ego_physco_down <- run_physco_enricher(down_physco)

      # Complete significant GO terms: these are the main result tables.
      physco_up_df <- add_physco_domain(
        if (is.null(ego_physco_up)) NULL else ego_physco_up$full
      )
      physco_down_df <- add_physco_domain(
        if (is.null(ego_physco_down)) NULL else ego_physco_down$full
      )

      # Redundancy-reduced GO terms: saved separately and used for the plot only.
      physco_up_simplified_df <- add_physco_domain(
        if (is.null(ego_physco_up)) NULL else ego_physco_up$simplified
      )
      physco_down_simplified_df <- add_physco_domain(
        if (is.null(ego_physco_down)) NULL else ego_physco_down$simplified
      )

      if (!is.null(physco_up_df)) {
        readr::write_tsv(
          physco_up_df,
          file.path("Report/GO_Physcomitrium/GO_results", paste0("GO_Physco_", cond, "_UP.tsv"))
        )
      }
      if (!is.null(physco_down_df)) {
        readr::write_tsv(
          physco_down_df,
          file.path("Report/GO_Physcomitrium/GO_results", paste0("GO_Physco_", cond, "_DOWN.tsv"))
        )
      }

      if (!is.null(physco_up_simplified_df)) {
        readr::write_tsv(
          physco_up_simplified_df,
          file.path(
            "Report/GO_Physcomitrium/GO_results_simplified",
            paste0("GO_Physco_", cond, "_UP_simplified.tsv")
          )
        )
        physco_up_simplified_df$Regulation <- "UP"
      }
      if (!is.null(physco_down_simplified_df)) {
        readr::write_tsv(
          physco_down_simplified_df,
          file.path(
            "Report/GO_Physcomitrium/GO_results_simplified",
            paste0("GO_Physco_", cond, "_DOWN_simplified.tsv")
          )
        )
        physco_down_simplified_df$Regulation <- "DOWN"
      }

      physco_all <- dplyr::bind_rows(
        physco_up_simplified_df,
        physco_down_simplified_df
      )
      if (nrow(physco_all) == 0) next

      physco_top <- physco_all %>%
        dplyr::mutate(
          plot_logp = -log10(pmax(p.adjust, .Machine$double.xmin)),
          Description_plot = ifelse(
            nchar(Description) > 60,
            paste0(substr(Description, 1, 57), "..."),
            Description
          )
        ) %>%
        dplyr::arrange(p.adjust) %>%
        utils::head(10)

      # Use the same capped significance scale as the TAIR GO plot.
      cap <- stats::quantile(physco_top$plot_logp, 0.9, na.rm = TRUE)
      physco_top$plot_logp_capped <- pmin(physco_top$plot_logp, cap)
      physco_top$signed_logp <- ifelse(
        physco_top$Regulation == "UP",
        physco_top$plot_logp_capped,
        -physco_top$plot_logp_capped
      )

      physco_plot <- ggplot2::ggplot(
        physco_top,
        ggplot2::aes(
          x = signed_logp,
          y = stats::reorder(Description_plot, signed_logp),
          size = Count,
          color = plot_logp
        )
      ) +
        ggplot2::geom_vline(
          xintercept = 0,
          linetype = "dashed",
          color = "grey60",
          linewidth = 0.6
        ) +
        ggplot2::geom_point() +
        ggplot2::labs(
          title = paste("Top 10 Physcomitrium GO terms -", cond),
          x = "signed -log10(adjusted p-value)",
          y = "GO term",
          size = "Gene count",
          color = "-log10(adj. p)"
        ) +
        ggplot2::scale_color_gradient(
          low = "skyblue",
          high = "orchid",
          limits = c(0, cap),
          oob = scales::squish
        ) +
        ggplot2::theme_classic(base_size = 14)

      print(physco_plot)
      ggplot2::ggsave(
        file.path("Report/GO_Physcomitrium", paste0("GO_Physco_top10_", cond, ".svg")),
        physco_plot,
        width = 11,
        height = 7,
        dpi = 300
      )
    }

    physco_go_coverage <- dplyr::bind_rows(physco_go_coverage)
    readr::write_tsv(
      physco_go_coverage,
      "Report/GO_Physcomitrium/annotation_coverage.tsv"
    )
    readr::write_tsv(
      data.frame(
        annotation_file = normalizePath(physcomitrium_go_file),
        go_terms = dplyr::n_distinct(physco_go$term),
        annotated_genes = dplyr::n_distinct(physco_go$gene),
        gene_term_associations = nrow(physco_term2gene)
      ),
      "Report/GO_Physcomitrium/annotation_summary.tsv"
    )
  }
```

#### 21 KEGG ENRICHMENT

```
if (run_go) {
dir_create("Report/KEGG/KEGG_results", recurse = TRUE)
dir_create("Report/KEGG/gene_lists", recurse = TRUE)

min_genes <- 10

for (cond in names(res_list)) {

  res_df <- res_list[[cond]]

  # Tieni solo righe con GeneID, log2FC, padj non-NA
  res_df <- res_df[complete.cases(res_df[, c("GeneID", "log2FoldChange", "padj")]), ]

  # Gene universe (TAIR IDs) + unique
  expressed_genes <- unique(res_df$GeneID)
  expressed_genes <- expressed_genes[!is.na(expressed_genes)]

  # UP/DOWN (TAIR IDs) + unique
  up_genes   <- unique(res_df$GeneID[res_df$padj < padj_cutoff & res_df$log2FoldChange > 0])
  down_genes <- unique(res_df$GeneID[res_df$padj < padj_cutoff & res_df$log2FoldChange < 0])

  if (length(up_genes) < min_genes && length(down_genes) < min_genes) next

  # Salva liste TAIR (come nel GO)
  if (length(up_genes) >= min_genes)
    write.table(data.frame(TAIR = up_genes),
                file = file.path("Report/KEGG/gene_lists", paste0("genes_", cond, "_UP_TAIR.tsv")),
                sep = "\t", quote = FALSE, row.names = FALSE)

  if (length(down_genes) >= min_genes)
    write.table(data.frame(TAIR = down_genes),
                file = file.path("Report/KEGG/gene_lists", paste0("genes_", cond, "_DOWN_TAIR.tsv")),
                sep = "\t", quote = FALSE, row.names = FALSE)

  # KEGG enrichment
  ek_up <- if (length(up_genes) >= min_genes) enrichKEGG(
    gene          = up_genes,
    organism      = "ath",
    keyType       = "kegg",
    universe      = expressed_genes,
    pAdjustMethod = "BH",
    qvalueCutoff  = 0.05
  ) else NULL

  ek_down <- if (length(down_genes) >= min_genes) enrichKEGG(
    gene          = down_genes,
    organism      = "ath",
    keyType       = "kegg",
    universe      = expressed_genes,
    pAdjustMethod = "BH",
    qvalueCutoff  = 0.05
  ) else NULL

  # Salva risultati
  if (!is.null(ek_up) && nrow(as.data.frame(ek_up)) > 0)
    write.table(as.data.frame(ek_up),
                file = file.path("Report/KEGG/KEGG_results", paste0("KEGG_", cond, "_UP.tsv")),
                sep = "\t", quote = FALSE, row.names = FALSE)

  if (!is.null(ek_down) && nrow(as.data.frame(ek_down)) > 0)
    write.table(as.data.frame(ek_down),
                file = file.path("Report/KEGG/KEGG_results", paste0("KEGG_", cond, "_DOWN.tsv")),
                sep = "\t", quote = FALSE, row.names = FALSE)

  # -----------------------------
  # GRAFICO TOP10 COME NEL GO
  # -----------------------------
  top_up <- if (!is.null(ek_up)) as.data.frame(ek_up) else NULL
  top_down <- if (!is.null(ek_down)) as.data.frame(ek_down) else NULL

  top_up <- if (!is.null(top_up) && nrow(top_up) > 0) { top_up$Regulation <- "UP"; top_up } else NULL
  top_down <- if (!is.null(top_down) && nrow(top_down) > 0) { top_down$Regulation <- "DOWN"; top_down } else NULL

  kegg_all <- do.call(rbind, Filter(Negate(is.null), list(top_up, top_down)))
  if (is.null(kegg_all) || nrow(kegg_all) == 0) next

  # Top 10 most significant
  kegg_top10 <- kegg_all %>%
    dplyr::mutate(logp = -log10(qvalue)) %>%
    dplyr::arrange(dplyr::desc(logp)) %>%
    head(10)

  # cap at the 90th percentile and compute a signed Z-score
  cap <- stats::quantile(kegg_top10$logp, 0.9, na.rm = TRUE)
  logp_capped <- pmin(kegg_top10$logp, cap)
  kegg_top10$Zscore <- ifelse(kegg_top10$Regulation == "UP", logp_capped, -logp_capped)

  # Plot combined
  kegg_plot <- ggplot2::ggplot(kegg_top10, ggplot2::aes(
    x = Zscore,
    y = stats::reorder(Description, Zscore),
    size = Count,
    color = logp
  )) +
    ggplot2::geom_vline(xintercept = 0, linetype = "dashed", color = "grey60", linewidth = 0.6) +
    ggplot2::geom_point() +
    ggplot2::labs(
      title = paste("Top 10 KEGG pathways -", cond),
      x = "KEGG enrichment Z-score",
      y = "KEGG pathway",
      size = "Gene Count",
      color = "-log10(p.adjust)"
    ) +
    ggplot2::scale_color_gradient(
      low = "skyblue",
      high = "orchid",
      limits = c(0, cap),
      oob = scales::squish
    ) +
    ggplot2::theme_classic(base_size = 14)

  print(kegg_plot)

  ggplot2::ggsave(
    filename = paste0("Report/KEGG/KEGG_top10_", cond, ".svg"),
    plot = kegg_plot, width = 10, height = 6, dpi = 300
  )
}
  
}
```

##Enrichment all sources

```
if (run_go) {
dir_create("Report/GOalternative/GOalternative_results", recurse = TRUE)
dir_create("Report/GOalternative/gene_lists", recurse = TRUE)

# --- Funzione dotplot GOalternative combinato UP/DOWN con Z-score ---
plot_reactome_updown_z <- function(res_up = NULL, res_down = NULL,
                                   title, out_path, top_n = 10, max_chars = 45) {
  df_up   <- if(!is.null(res_up)   && nrow(res_up)   > 0) res_up   else data.frame()
  df_down <- if(!is.null(res_down) && nrow(res_down) > 0) res_down else data.frame()

  if(nrow(df_up) == 0 && nrow(df_down) == 0) return(NULL)

  # gprofiler2 output: term_name, p_value, intersection_size, term_id, source, ...
  # Nota: p_value è già "significativo" secondo correzione g:SCS (default).
  if(nrow(df_up) > 0) {
    df_up$Zscore <- -log10(df_up$p_value)
    df_up$Regulation <- "UP"
  }
  if(nrow(df_down) > 0) {
    df_down$Zscore <-  log10(df_down$p_value)  # negativo
    df_down$Regulation <- "DOWN"
  }

  df <- rbind(df_up, df_down)

  # Top terms by |Z|
  df <- df %>% arrange(desc(abs(Zscore))) %>% head(top_n)

  # Accorcia label
  df$term_name_short <- sapply(df$term_name, function(x) {
    x <- trimws(x)
    if(nchar(x) > max_chars) paste0(substr(x, 1, max_chars - 3), "…") else x
  })

  gg <- ggplot(df, aes(x = Zscore,
                       y = reorder(term_name_short, Zscore),
                       size = intersection_size,
                       color = -log10(p_value))) +
    geom_point() +
    scale_color_gradient(low = "skyblue", high = "orchid") +
    labs(title = title,
         x = " Z-score",
         y = " Pathway",
         size = "N° geni (overlap)",
         color = "-log10(p-value)") +
    theme_bw(base_size = 14)

  print(gg)
  ggsave(out_path, gg, width = 11, height = 7, dpi = 300)

  return(gg)
}

# --- Prepara liste UP/DOWN da res_list ---
deg_lists <- list()

for(pair_id in names(res_list)) {

  res_df <- res_list[[pair_id]]
  res_df <- res_df[complete.cases(res_df[, c("GeneID", "log2FoldChange", "padj")]), ]

  deg_lists[[paste0(pair_id, "_UP")]]       <- unique(res_df$GeneID[res_df$padj < padj_cutoff & res_df$log2FoldChange > 0])
  deg_lists[[paste0(pair_id, "_DOWN")]]     <- unique(res_df$GeneID[res_df$padj < padj_cutoff & res_df$log2FoldChange < 0])
  deg_lists[[paste0(pair_id, "_universe")]] <- unique(res_df$GeneID)
}

# --- Enrichment Reactome + plot ---
reactome_results_list <- list()

for(pair_id in unique(sub("_UP|_DOWN|_universe", "", names(deg_lists)))) {

  up_genes        <- toupper(deg_lists[[paste0(pair_id, "_UP")]])
  down_genes      <- toupper(deg_lists[[paste0(pair_id, "_DOWN")]])
  expressed_genes <- toupper(deg_lists[[paste0(pair_id, "_universe")]])

  if(length(up_genes) < 5 && length(down_genes) < 5) next

  # Save gene lists
  if(length(up_genes) >= 5) write.table(data.frame(GeneID = up_genes),
                                        file = file.path("Report/GOalternative/gene_lists", paste0("genes_", pair_id, "_UP.tsv")),
                                        sep = "\t", quote = FALSE, row.names = FALSE)

  if(length(down_genes) >= 5) write.table(data.frame(GeneID = down_genes),
                                          file = file.path("Report/GOalternative/gene_lists", paste0("genes_", pair_id, "_DOWN.tsv")),
                                          sep = "\t", quote = FALSE, row.names = FALSE)

  # --- Enrichment con gprofiler2 ---
  # organism for Arabidopsis: "athaliana"
  # background custom: expressed_genes
  res_up <- NULL
  res_down <- NULL

    gost_up <- gost(query = up_genes,
           organism = "athaliana",
           custom_bg = expressed_genes,
           correction_method = "g_SCS")
     
    if(!is.null(gost_up) && !is.null(gost_up$result) && nrow(gost_up$result) > 0) {
      res_up <- gost_up$result
      parents_go <- unique(unlist(res_up$parents[grepl("^GO:", res_up$source)]))
      res_up <- dplyr::filter(res_up, !term_id %in% parents_go)
      res_up <- dplyr::select(res_up, dplyr::where(~ !is.list(.)))
      
      
    }

    gost_down <- gost(query = down_genes,
           organism = "athaliana",
           custom_bg = expressed_genes,
           correction_method = "g_SCS")
    if(!is.null(gost_down) && !is.null(gost_down$result) && nrow(gost_down$result) > 0) {
      res_down <- gost_down$result
      parents_go <- unique(unlist(res_down$parents[grepl("^GO:", res_down$source)]))
      res_down <- dplyr::filter(res_down, !term_id %in% parents_go)
      res_down <- dplyr::select(res_down, dplyr::where(~ !is.list(.)))
    }
    
    
  # Salva risultati
  if(!is.null(res_up)) write.table(res_up,
                                  file = file.path("Report/GOalternative/GOalternative_results", paste0("Reactome_", pair_id, "_UP.tsv")),
                                  sep = "\t", quote = FALSE, row.names = FALSE)

  if(!is.null(res_down)) write.table(res_down,
                                     file = file.path("Report/GOalternative/GOalternative_results", paste0("Reactome_", pair_id, "_DOWN.tsv")),
                                     sep = "\t", quote = FALSE, row.names = FALSE)

  # Plot combinato UP/DOWN con Z-score
  reactome_results_list[[pair_id]] <- plot_reactome_updown_z(
    res_up, res_down,
    title = paste("Enrichment -", pair_id),
    out_path = file.path("Report/GOalternative", paste0("GOalternative_", pair_id, "_UPDOWN.svg")),
    top_n = 10,
    max_chars = 55
  )
}
}
```

#### 22 GSEA

```
if (run_gsea) {

  fs::dir_create("Report/GSEA", recurse = TRUE)

  for (cond in names(res_list)) {

    res_df <- res_list[[cond]]

    # --------------------------------------------------
    # Preparazione della ranked list
    # --------------------------------------------------

    res_df <- res_df[
      !is.na(res_df$GeneID) &
      nzchar(res_df$GeneID) &
      is.finite(res_df$stat_unshrunk),
      ,
      drop = FALSE
    ]

    if (nrow(res_df) == 0) {
      message("Nessun gene TAIR valido per ", cond)
      next
    }

    res_df$GeneID <- toupper(as.character(res_df$GeneID))

    # Più geni di P. patens possono avere lo stesso omologo TAIR.
    # Per ogni TAIR mantieni il gene con |stat| maggiore.
    res_df <- res_df %>%
      dplyr::arrange(dplyr::desc(abs(stat_unshrunk))) %>%
      dplyr::distinct(GeneID, .keep_all = TRUE)

    gene_list <- res_df$stat_unshrunk
    names(gene_list) <- res_df$GeneID

    # gseGO richiede un vettore ordinato in ordine decrescente
    gene_list <- sort(gene_list, decreasing = TRUE)

    if (length(gene_list) < 10) {
      message("Troppi pochi geni TAIR per la GSEA di ", cond)
      next
    }

    stopifnot(
      !anyNA(gene_list),
      !anyNA(names(gene_list)),
      !anyDuplicated(names(gene_list)),
      all(is.finite(gene_list))
    )

    # --------------------------------------------------
    # GSEA
    # --------------------------------------------------

    gsea_res <- tryCatch(
      {
        clusterProfiler::gseGO(
          geneList = gene_list,
          OrgDb = org.At.tair.db,
          ont = "BP",
          keyType = "TAIR",
          minGSSize = 10,
          maxGSSize = 500,
          pvalueCutoff = 0.05,
          pAdjustMethod = "BH",
          verbose = FALSE,
          eps = 0,
          nPermSimple = 50000
        )
      },
      error = function(e) {
        message("Errore GSEA per ", cond, ": ", conditionMessage(e))
        NULL
      }
    )

    if (is.null(gsea_res)) next

    gsea_df_full <- as.data.frame(gsea_res)

    if (nrow(gsea_df_full) == 0) {
      message("Nessun termine arricchito per ", cond)
      next
    }

    # --------------------------------------------------
    # Salva core-enrichment non semplificato
    # --------------------------------------------------

    required_columns <- c("ID", "Description", "core_enrichment")

    if (all(required_columns %in% colnames(gsea_df_full))) {

      gene_lists_df <- gsea_df_full[, required_columns, drop = FALSE]

      gene_lists_df$core_enrichment <- gsub(
        "/",
        ",",
        gene_lists_df$core_enrichment,
        fixed = TRUE
      )

      write.table(
        gene_lists_df,
        file = file.path(
          "Report/GSEA",
          paste0("GSEA_genes_", cond, ".tsv")
        ),
        sep = "\t",
        quote = FALSE,
        row.names = FALSE
      )
    }

    # --------------------------------------------------
    # Rimozione dei termini GO ridondanti
    # --------------------------------------------------

    gsea_res_simplified <- tryCatch(
      {
        clusterProfiler::simplify(
          gsea_res,
          cutoff = cutoff_val,
          by = "p.adjust",
          select_fun = min,
          measure = "Wang"
        )
      },
      error = function(e) {
        message(
          "Simplify non riuscito per ", cond,
          "; uso risultati non semplificati: ",
          conditionMessage(e)
        )
        gsea_res
      }
    )

    gsea_df <- as.data.frame(gsea_res_simplified)

    gsea_df <- gsea_df[
      is.finite(gsea_df$NES) &
      is.finite(gsea_df$p.adjust),
      ,
      drop = FALSE
    ]

    if (nrow(gsea_df) == 0) {
      message("Nessun NES valido per ", cond)
      next
    }

    # Coerenza: ordina per p.adjust, usato anche nel grafico
    top <- head(
      gsea_df[order(gsea_df$p.adjust), , drop = FALSE],
      20
    )

    # Evita Inf quando p.adjust è numericamente zero
    top$plot_logp <- -log10(
      pmax(top$p.adjust, .Machine$double.xmin)
    )

    # --------------------------------------------------
    # Grafico
    # --------------------------------------------------

    gsea_plot <- ggplot2::ggplot(
      top,
      ggplot2::aes(
        x = stats::reorder(Description, NES),
        y = NES,
        size = setSize,
        color = plot_logp
      )
    ) +
      ggplot2::geom_point() +
      ggplot2::coord_flip() +
      ggplot2::labs(
        title = paste("Top 20 GO terms -", cond),
        x = "GO term",
        y = "Normalized Enrichment Score (NES)",
        size = "Gene count",
        color = "-log10(p.adjust)"
      ) +
      ggplot2::scale_color_gradient(
        low = "skyblue",
        high = "orchid"
      ) +
      ggplot2::theme_classic(base_size = 14)

    print(gsea_plot)

    ggplot2::ggsave(
      filename = file.path(
        "Report/GSEA",
        paste0("GSEA_top20_", cond, ".svg")
      ),
      plot = gsea_plot,
      width = 9,
      height = 6,
      dpi = 300
    )
  }
}
```

#### 23 GSEA GO — *Physcomitrium patens*

```
if (run_gsea_physco) {

  fs::dir_create("Report/GSEA_Physcomitrium/GSEA_results", recurse = TRUE)
  fs::dir_create("Report/GSEA_Physcomitrium/core_enrichment", recurse = TRUE)

  # Reuse the cleaned annotation produced by the Physcomitrium GO module.
  # If that module was disabled, load and clean the annotation here.
  if (!exists("normalise_physco_id")) {
    normalise_physco_id <- function(x) {
      x <- toupper(trimws(as.character(x)))
      x <- gsub("^[\"']|[\"']$", "", x)
      x <- sub("\\.[0-9]+$", "", x)
      x
    }
  }

  if (!exists("physco_term2gene") ||
      !exists("physco_term2name") ||
      !exists("physco_term_meta")) {

    physco_gsea_candidates <- unique(c(
      physcomitrium_go_file,
      if (exists("db_dir")) file.path(db_dir, physcomitrium_go_file),
      "physcomitrium_GO",
      "physcomitrium_GO.tsv",
      "physcomitrium_GO.txt",
      "physcomitrium_GO.csv",
      if (exists("db_dir")) file.path(db_dir, c(
        "physcomitrium_GO", "physcomitrium_GO.tsv",
        "physcomitrium_GO.txt", "physcomitrium_GO.csv"
      ))
    ))
    physco_gsea_candidates <- physco_gsea_candidates[
      file.exists(physco_gsea_candidates)
    ]

    if (length(physco_gsea_candidates) == 0) {
      stop(
        "Physcomitrium GSEA cannot start: GO annotation file not found. ",
        "Set physcomitrium_go_file in the launcher."
      )
    }

    physcomitrium_go_file <- physco_gsea_candidates[[1]]
    physco_go_gsea <- readr::read_delim(
      physcomitrium_go_file,
      delim = NULL,
      trim_ws = TRUE,
      show_col_types = FALSE,
      progress = FALSE
    )
    names(physco_go_gsea) <- tolower(trimws(names(physco_go_gsea)))

    required_gsea_go_cols <- c("term", "gene", "name", "domain")
    missing_gsea_go_cols <- setdiff(
      required_gsea_go_cols,
      names(physco_go_gsea)
    )
    if (length(missing_gsea_go_cols) > 0) {
      stop(
        "Missing columns in ", physcomitrium_go_file, ": ",
        paste(missing_gsea_go_cols, collapse = ", ")
      )
    }

    physco_go_gsea <- physco_go_gsea %>%
      dplyr::transmute(
        term = trimws(as.character(term)),
        gene = normalise_physco_id(gene),
        name = trimws(as.character(name)),
        domain = tolower(trimws(as.character(domain)))
      ) %>%
      dplyr::filter(
        !is.na(term), grepl("^GO:[0-9]{7}$", term),
        !is.na(gene), nzchar(gene),
        !is.na(name), nzchar(name)
      ) %>%
      dplyr::distinct(term, gene, .keep_all = TRUE)

    if (nrow(physco_go_gsea) == 0) {
      stop("No valid GO-gene associations found in ", physcomitrium_go_file)
    }

    physco_term_meta <- physco_go_gsea %>%
      dplyr::group_by(term) %>%
      dplyr::summarise(
        name = dplyr::first(name),
        domain = paste(sort(unique(domain[nzchar(domain)])), collapse = ";"),
        .groups = "drop"
      )
    physco_term2gene <- physco_go_gsea %>%
      dplyr::select(term, gene) %>%
      dplyr::distinct()
    physco_term2name <- physco_term_meta %>%
      dplyr::select(term, name)
  }

  annotated_physco_genes_gsea <- unique(physco_term2gene$gene)
  physco_gsea_coverage <- list()

  for (cond in names(res_list)) {
    res_df <- res_list[[cond]]
    res_df$PhyscoGene <- normalise_physco_id(rownames(res_df))

    ranked_df <- res_df[
      !is.na(res_df$PhyscoGene) &
      nzchar(res_df$PhyscoGene) &
      is.finite(res_df$stat_unshrunk),
      ,
      drop = FALSE
    ]

    # One ranking value per gene. When several rows collapse to the same gene ID,
    # retain the one with the strongest absolute Wald statistic.
    ranked_df <- ranked_df %>%
      dplyr::arrange(dplyr::desc(abs(stat_unshrunk))) %>%
      dplyr::distinct(PhyscoGene, .keep_all = TRUE)

    matched_ranked_genes <- intersect(
      ranked_df$PhyscoGene,
      annotated_physco_genes_gsea
    )
    physco_gsea_coverage[[cond]] <- data.frame(
      contrast = cond,
      ranked_genes = nrow(ranked_df),
      annotated_ranked_genes = length(matched_ranked_genes),
      annotation_coverage_percent = round(
        100 * length(matched_ranked_genes) / max(1, nrow(ranked_df)),
        2
      )
    )

    if (length(matched_ranked_genes) < 10) {
      message(
        "Too few annotated Physcomitrium genes for GSEA of ", cond,
        ": ", length(matched_ranked_genes)
      )
      next
    }

    gene_list_physco <- ranked_df$stat_unshrunk
    names(gene_list_physco) <- ranked_df$PhyscoGene
    gene_list_physco <- sort(gene_list_physco, decreasing = TRUE)

    stopifnot(
      !anyNA(gene_list_physco),
      !anyNA(names(gene_list_physco)),
      !anyDuplicated(names(gene_list_physco)),
      all(is.finite(gene_list_physco))
    )

    gsea_physco <- tryCatch(
      clusterProfiler::GSEA(
        geneList = gene_list_physco,
        TERM2GENE = physco_term2gene,
        TERM2NAME = physco_term2name,
        minGSSize = 10,
        maxGSSize = 500,
        pvalueCutoff = 0.05,
        pAdjustMethod = "BH",
        verbose = FALSE,
        seed = TRUE,
        eps = 0,
        by = "fgsea"
      ),
      error = function(e) {
        message(
          "Physcomitrium GSEA failed for ", cond, ": ",
          conditionMessage(e)
        )
        NULL
      }
    )

    if (is.null(gsea_physco)) next
    gsea_physco_df <- as.data.frame(gsea_physco)
    if (nrow(gsea_physco_df) == 0) {
      message("No significant Physcomitrium GO terms for ", cond)
      next
    }

    gsea_physco_df <- gsea_physco_df %>%
      dplyr::left_join(
        physco_term_meta %>% dplyr::select(term, domain),
        by = c("ID" = "term")
      )

    readr::write_tsv(
      gsea_physco_df,
      file.path(
        "Report/GSEA_Physcomitrium/GSEA_results",
        paste0("GSEA_Physco_", cond, ".tsv")
      )
    )

    if (all(c("ID", "Description", "NES", "core_enrichment") %in%
            names(gsea_physco_df))) {
      core_physco <- gsea_physco_df %>%
        dplyr::select(ID, Description, domain, NES, core_enrichment) %>%
        dplyr::mutate(
          core_enrichment = gsub("/", ",", core_enrichment, fixed = TRUE)
        )
      readr::write_tsv(
        core_physco,
        file.path(
          "Report/GSEA_Physcomitrium/core_enrichment",
          paste0("GSEA_Physco_core_genes_", cond, ".tsv")
        )
      )
    }

    top_physco_gsea <- gsea_physco_df %>%
      dplyr::filter(is.finite(NES), is.finite(p.adjust)) %>%
      dplyr::arrange(p.adjust) %>%
      utils::head(20) %>%
      dplyr::mutate(
        plot_logp = -log10(pmax(p.adjust, .Machine$double.xmin)),
        domain_label = dplyr::case_when(
          domain == "biological_process" ~ "BP",
          domain == "molecular_function" ~ "MF",
          domain == "cellular_component" ~ "CC",
          TRUE ~ domain
        ),
        Description_plot = paste0("[", domain_label, "] ", Description),
        Description_plot = ifelse(
          nchar(Description_plot) > 70,
          paste0(substr(Description_plot, 1, 67), "..."),
          Description_plot
        )
      )

    if (nrow(top_physco_gsea) == 0) next

    gsea_physco_plot <- ggplot2::ggplot(
      top_physco_gsea,
      ggplot2::aes(
        x = stats::reorder(Description_plot, NES),
        y = NES,
        size = setSize,
        color = plot_logp
      )
    ) +
      ggplot2::geom_hline(
        yintercept = 0,
        linetype = "dashed",
        color = "grey60",
        linewidth = 0.6
      ) +
      ggplot2::geom_point() +
      ggplot2::coord_flip() +
      ggplot2::scale_color_gradient(low = "skyblue", high = "orchid") +
      ggplot2::labs(
        title = paste("Top 20 Physcomitrium GO GSEA terms -", cond),
        x = "GO term",
        y = "Normalized Enrichment Score (NES)",
        size = "Gene-set size",
        color = "-log10(p.adjust)"
      ) +
      ggplot2::theme_classic(base_size = 14)

    print(gsea_physco_plot)
    ggplot2::ggsave(
      file.path(
        "Report/GSEA_Physcomitrium",
        paste0("GSEA_Physco_top20_", cond, ".svg")
      ),
      gsea_physco_plot,
      width = 11,
      height = 8,
      dpi = 300
    )
  }

  readr::write_tsv(
    dplyr::bind_rows(physco_gsea_coverage),
    "Report/GSEA_Physcomitrium/annotation_coverage.tsv"
  )
}
```

#### 24 MAPMAN GO Enrichment

```
if (run_mapman_go) {

# Crea cartelle
dir_create("Report/MapMan/MapMan_results", recurse = TRUE)
dir_create("Report/MapMan/gene_lists", recurse = TRUE)

# --- Carica file di mappatura MapMan ---
mapman_x4 <- read_excel(file.path(db_dir, "X4_Araport11_R1.0.xls"))
mapman_mapping <- mapman_x4 %>%
  dplyr::select(GeneID = "IDENTIFIER", BinCode = "BINCODE", Description = "NAME")
mapman_mapping$GeneID <- toupper(mapman_mapping$GeneID)

# --- Funzione dotplot MapMan combinato UP/DOWN con Z-score ---
plot_mapman_updown_z <- function(ego_up = NULL, ego_down = NULL,
                                 title, out_path, top_n = 10, max_chars = 20) {
  # Converti in data frame
  df_up <- if(!is.null(ego_up)) as.data.frame(ego_up) else data.frame()
  df_down <- if(!is.null(ego_down)) as.data.frame(ego_down) else data.frame()
  
  if(nrow(df_up) == 0 & nrow(df_down) == 0) return(NULL)
  
  # Z-score basato su -log10(qvalie)
  if(nrow(df_up) > 0) {
    df_up$Zscore <- -log10(df_up$qvalue)  # UP = positivo
    df_up$Regulation <- "UP"
  }
  if(nrow(df_down) > 0) {
    df_down$Zscore <- log10(df_down$qvalue)  # DOWN = negativo
    df_down$Regulation <- "DOWN"
  }
  
  # Combine UP e DOWN
  df <- rbind(df_up, df_down)
  
  # Top termini in base al valore assoluto di Z-score
  df <- df %>% arrange(desc(abs(Zscore))) %>% head(top_n)
  
 # Prendi ultimi due livelli della descrizione (separati da ".") e tronca se troppo lunghi
  df$Description_short <- sapply(df$Description, function(x) {
    # Split solo sul punto
    levels <- unlist(strsplit(x, "\\.", fixed = FALSE))
    last2 <- tail(levels, 2)
    last2 <- sapply(last2, function(lvl) {
      lvl <- trimws(lvl)  # rimuove eventuali spazi iniziali/finali
      if(nchar(lvl) > max_chars) paste0(substr(lvl,1,max_chars-3),"…") else lvl
    })
    paste(last2, collapse = "\n")  # a capo per leggibilità
  })
  
  # Plot combinato
  gg <- ggplot(df, aes(x = Zscore, y = reorder(Description_short, Zscore),
                       size = Count, color = -log10(p.adjust))) +
    geom_point() +
    scale_color_gradient(low = "skyblue", high = "orchid") +
    labs(title = title, x = "MapMan Z-score",
         y = "MapMan Bin", size = "N° geni", color = "-log10(p.adjust)") +
    theme_bw(base_size = 14)
  
  print(gg)
  ggsave(out_path, gg, width = 10, height = 7, dpi = 300)
  
  return(gg)
}

# --- Prepara liste UP/DOWN da res_list ---
deg_lists <- list()

for(pair_id in names(res_list)) {
  
  res_df <- res_list[[pair_id]]
  res_df <- res_df[complete.cases(res_df[, c("GeneID", "log2FoldChange", "padj")]), ]

  deg_lists[[paste0(pair_id, "_UP")]]       <- unique(res_df$GeneID[res_df$padj < padj_cutoff & res_df$log2FoldChange > 0])
  deg_lists[[paste0(pair_id, "_DOWN")]]     <- unique(res_df$GeneID[res_df$padj < padj_cutoff & res_df$log2FoldChange < 0])
  deg_lists[[paste0(pair_id, "_universe")]] <- unique(res_df$GeneID)
}

# --- Enrichment MapMan e plot ---
mapman_results_list <- list()

for(pair_id in unique(sub("_UP|_DOWN", "", names(deg_lists)))) {
  up_genes   <- toupper(deg_lists[[paste0(pair_id, "_UP")]])
  down_genes <- toupper(deg_lists[[paste0(pair_id, "_DOWN")]])
  expressed_genes <- toupper(deg_lists[[paste0(pair_id, "_universe")]])
  
  if(length(up_genes) < 5 & length(down_genes) < 5) next
  
  # Save gene lists
  if(length(up_genes) >= 5) write.table(data.frame(GeneID = up_genes),
                                        file = file.path("Report/MapMan/gene_lists", paste0("genes_", pair_id, "_UP.tsv")),
                                        sep="\t", quote=FALSE, row.names=FALSE)
  if(length(down_genes) >= 5) write.table(data.frame(GeneID = down_genes),
                                          file = file.path("Report/MapMan/gene_lists", paste0("genes_", pair_id, "_DOWN.tsv")),
                                          sep="\t", quote=FALSE, row.names=FALSE)
  
  # Enrichment MapMan
  ego_up   <- if(length(up_genes) >= 5) enricher(up_genes, universe = expressed_genes,
                                                 TERM2GENE = mapman_mapping[, c("BinCode","GeneID")],
                                                 TERM2NAME = mapman_mapping[, c("BinCode","Description")],
                                                 pAdjustMethod="BH", qvalueCutoff=0.05) else NULL
  ego_down <- if(length(down_genes) >= 5) enricher(down_genes, universe = expressed_genes,
                                                   TERM2GENE = mapman_mapping[, c("BinCode","GeneID")],
                                                   TERM2NAME = mapman_mapping[, c("BinCode","Description")],
                                                   pAdjustMethod="BH", qvalueCutoff=0.05) else NULL
  
  # Salva risultati MapMan
  if(!is.null(ego_up)) write.table(as.data.frame(ego_up),
                                   file=file.path("Report/MapMan/MapMan_results", paste0("MapMan_", pair_id, "_UP.tsv")),
                                   sep="\t", quote=FALSE, row.names=FALSE)
  if(!is.null(ego_down)) write.table(as.data.frame(ego_down),
                                     file=file.path("Report/MapMan/MapMan_results", paste0("MapMan_", pair_id, "_DOWN.tsv")),
                                     sep="\t", quote=FALSE, row.names=FALSE)
  
  # Plot combinato UP/DOWN con Z-score
  mapman_results_list[[pair_id]] <- plot_mapman_updown_z(
    ego_up, ego_down,
    title = paste("MapMan enrichment -", pair_id),
    out_path = file.path("Report/MapMan", paste0("MapMan_", pair_id, "_UPDOWN.svg")),
    top_n = 10,
    max_chars = 40
  )
}

} # end run_mapman_go
```

#### 25 MAPMAP HEAT

```
if (run_mapman_heat) {
# ------------------------------
# Step 0: Preparazione
# ------------------------------
dir_create("Report/MapManBin", recurse = TRUE)

# Prepara mapman con Class e ClassLabel robusti
mapman <- mapman_x4 %>%
  mutate(
    BINCODE = as.character(BINCODE),
    NAME = as.character(NAME),
    Class = sapply(strsplit(BINCODE, "\\."), function(x) paste(head(x, 3), collapse = ".")),
    ClassLabel = sapply(strsplit(NAME, "\\."), function(x) {
      y <- x[2:min(3, length(x))]
      if (length(y) == 0) x[1] else paste(y, collapse = ".")
    }),
    IDENTIFIER = toupper(as.character(IDENTIFIER))
  )

original_order <- unique(mapman$Class[!is.na(mapman$Class)])

for (pair in names(res_list)) {

  res_df <- res_list[[pair]]
  if (is.null(res_df) || nrow(res_df) == 0) next
  if (!all(c("log2FoldChange","padj") %in% colnames(res_df))) next

  res_df <- res_df[complete.cases(res_df[, c("GeneID","log2FoldChange","padj")]), ]

  # (opzionale) se vuoi solo significativi:
  # res_df <- res_df[res_df$padj < padj & abs(res_df$log2FoldChange) >= log2foldchange, ]
  # if (nrow(res_df) == 0) next

  res_class <- merge(
    res_df[, c("GeneID", "log2FoldChange")],
    mapman[, c("IDENTIFIER", "Class", "ClassLabel")],
    by.x = "GeneID", by.y = "IDENTIFIER",
    all.x = TRUE
  )

  res_class <- res_class[!is.na(res_class$Class), ]
  if (nrow(res_class) == 0) next

  res_class$Class <- factor(res_class$Class, levels = original_order)

  log2_by_class <- res_class %>%
    dplyr::group_by(Class, ClassLabel) %>%
    dplyr::summarise(mean_log2FC = mean(log2FoldChange, na.rm = TRUE), .groups = "drop") %>%
    dplyr::filter(mean_log2FC > 1 | mean_log2FC < -1)

  if (nrow(log2_by_class) == 0) next

  write.table(log2_by_class,
              file = paste0("Report/MapManBin/log2_by_class_", pair, ".tsv"),
              sep = "\t", row.names = FALSE, quote = FALSE)

  top_n <- min(10, nrow(log2_by_class))
  log2_by_class_extremes <- dplyr::bind_rows(
    dplyr::slice_max(log2_by_class, order_by = mean_log2FC, n = top_n),
    dplyr::slice_min(log2_by_class, order_by = mean_log2FC, n = top_n)
  ) %>%
    dplyr::distinct(Class, ClassLabel, .keep_all = TRUE) %>%
    dplyr::arrange(dplyr::desc(mean_log2FC))

  log2_by_class_extremes$x <- 1

  log2_by_class_extremes$ClassLabel_plot <- vapply(log2_by_class_extremes$ClassLabel, function(lbl) {
    lbl <- gsub("\\.", "\n", lbl)
    parts <- strsplit(lbl, "\n")[[1]]
    parts <- vapply(parts, function(line) {
      if (nchar(line) > 60) paste0(substr(line, 1, 57), "…") else line
    }, character(1))
    paste(parts, collapse = "\n")
  }, character(1))

  binplot <- ggplot(log2_by_class_extremes,
                    aes(x = x, y = reorder(ClassLabel_plot, mean_log2FC), fill = mean_log2FC)) +
    geom_tile(width = 0.8, height = 0.8, color = "white") +
    scale_fill_gradient2(low = "skyblue", mid = "white", high = "orchid", midpoint = 0) +
    theme_minimal() +
    theme(
      axis.text.x = element_blank(),
      axis.ticks.x = element_blank(),
      axis.title = element_blank(),
      panel.grid = element_blank(),
      axis.text.y = element_text(size = 10, hjust = 1)
    ) +
    coord_fixed(ratio = 0.5) +
    ggtitle(paste0("Top MAPMAN classes ", pair))

  print(binplot)
  ggsave(
    filename = paste0("Report/MapManBin/BINMAP_", pair, ".svg"),
    plot = binplot, width = 9, height = 6, dpi = 300, limitsize = FALSE
  )
}


} # end run_mapman_heat
```

#### 26 TF enrichment based on PlantTFDB

```
if (run_tf_enrichment) {

dir_create("Report/TFEnrichment", recurse = TRUE)

atreg <- read.csv(file.path(db_dir, "AtRegNet.csv"),
                  header = TRUE, row.names = NULL, check.names = TRUE)

names(atreg)[1:(ncol(atreg)-1)] <- names(atreg)[2:ncol(atreg)]
atreg <- atreg[, c("TFName", "TargetLocus")]
colnames(atreg) <- c("TF", "target.gene")

arab_cat <- na.omit(atreg)
arab_cat$target.gene <- toupper(arab_cat$target.gene)

for (pair in names(res_list)) {

  res_df <- res_list[[pair]]
  if (is.null(res_df) || nrow(res_df) == 0) next

  # serve anche baseMean
  if (!all(c("log2FoldChange","padj") %in% colnames(res_df))) next

  res_df <- res_df[complete.cases(res_df[, c("GeneID","log2FoldChange","padj")]), ]

  # Expressed universe (consistent and uppercase)
  universe_genes <- toupper(unique(res_df$GeneID))
  if (length(universe_genes) == 0) next

  # DEGs significativi
  sig_genes <- toupper(res_df$GeneID[res_df$padj < padj & abs(res_df$log2FoldChange) > log2foldchange])
  sig_genes <- intersect(sig_genes, universe_genes)
  if (length(sig_genes) == 0) next

  # Restrict the catalog to genes in the universe
  arab_cat_red <- arab_cat[arab_cat$target.gene %in% universe_genes, ]
  if (nrow(arab_cat_red) == 0) next

  tf_results <- list()

  for (tf in unique(arab_cat_red$TF)) {

    targets <- unique(arab_cat_red$target.gene[arab_cat_red$TF == tf])
    targets_in_universe <- intersect(targets, universe_genes)
    if (length(targets_in_universe) < 3) next

    a <- sum(targets_in_universe %in% sig_genes)
    b <- length(targets_in_universe) - a
    non_targets <- setdiff(universe_genes, targets_in_universe)
    c <- sum(non_targets %in% sig_genes)
    d <- length(non_targets) - c

    mat <- matrix(c(a, b, c, d), nrow = 2)
    if (any(mat < 0) || sum(mat) == 0) next

    ft <- fisher.test(mat, alternative = "greater")

    tf_results[[tf]] <- data.frame(
      TF = tf,
      Targets = length(targets_in_universe),
      DEGs_in_targets = a,
      pvalue = ft$p.value,
      odds_ratio = unname(ft$estimate)
    )
  }

  tf_res <- do.call(rbind, tf_results)
  if (is.null(tf_res) || nrow(tf_res) == 0) {
    message("Nessun TF arricchito per ", pair)
    next
  }

  tf_res <- tf_res %>%
    mutate(FDR.BH = p.adjust(pvalue, method = "BH")) %>%
    arrange(FDR.BH)

  # pulizia OR non finiti
  tf_res$odds_ratio[!is.finite(tf_res$odds_ratio)] <- NA

  write.table(tf_res,
              file = paste0("Report/TFEnrichment/TFactSR_results_", pair, ".tsv"),
              sep = "\t", row.names = FALSE, quote = FALSE)

  tf_signif <- tf_res %>% filter(FDR.BH < 0.05 & !is.na(odds_ratio))
  if (nrow(tf_signif) == 0) {
    message("Nessun TF significativamente arricchito per ", pair)
    next
  }

  top_n <- min(10, nrow(tf_signif))
  tf_top <- tf_signif %>%
    arrange(FDR.BH) %>%
    slice(1:top_n) %>%
    mutate(
      TF_plot = ifelse(nchar(TF) > 20, paste0(substr(TF, 1, 17), "…"), TF),
      negLogFDR = -log10(FDR.BH)
    )

  xmax <- max(tf_top$odds_ratio, na.rm = TRUE)
  if (!is.finite(xmax)) xmax <- 1

  tf_plot <- ggplot(tf_top, aes(x = odds_ratio, y = reorder(TF_plot, odds_ratio), fill = negLogFDR)) +
    geom_col(width = 0.7) +
    geom_text(aes(label = sprintf("%.2f", odds_ratio)),
              hjust = -0.1, size = 3.2, color = "black") +
    scale_fill_gradient(low = "skyblue", high = "orchid") +
    theme_minimal() +
    theme(
      axis.title.y = element_blank(),
      axis.title.x = element_text(size = 10),
      axis.text.y = element_text(size = 10, hjust = 1),
      panel.grid.major.y = element_blank()
    ) +
    labs(x = "Odds Ratio", fill = "-log10(FDR)") +
    ggtitle(paste0("Top enriched TFs ", pair)) +
    coord_cartesian(xlim = c(0, xmax * 1.2))

  print(tf_plot)

  ggsave(
    filename = paste0("Report/TFEnrichment/TFactSR_topTF_", pair, ".svg"),
    plot = tf_plot,
    width = 8, height = 6, dpi = 300, limitsize = FALSE
  )
}


} # end run_tf_enrichment
```

#### 27 Reproducibility

```
if (requireNamespace("sessioninfo", quietly = TRUE)) {
  sessioninfo::session_info()
} else {
  sessionInfo()
}
```

```
## ─ Session info ───────────────────────────────────────────────────────────────────────────────────────────────────────────────────────────────────────────────────────────────────
##  setting  value
##  version  R version 4.4.3 (2025-02-28 ucrt)
##  os       Windows 11 x64 (build 26200)
##  system   x86_64, mingw32
##  ui       RStudio
##  language (EN)
##  collate  Italian_Italy.utf8
##  ctype    Italian_Italy.utf8
##  tz       Europe/Rome
##  date     2026-09-29
##  rstudio  2026.09.0+174 Autumn Hawkbit (desktop)
##  pandoc   3.10 @ C:/Program Files/RStudio/resources/app/bin/quarto/bin/tools/ (via rmarkdown)
##  quarto   1.10.18 @ C:\\PROGRA~1\\RStudio\\RESOUR~1\\app\\bin\\quarto\\bin\\quarto.exe
## 
## ─ Packages ───────────────────────────────────────────────────────────────────────────────────────────────────────────────────────────────────────────────────────────────────────
##  package              * version   date (UTC) lib source
##  abind                  1.4-8     2024-09-12 [1] CRAN (R 4.4.1)
##  AnnotationDbi        * 1.68.0    2024-10-29 [1] Bioconductor 3.20 (R 4.4.1)
##  ape                    5.8-1     2024-12-16 [1] CRAN (R 4.4.3)
##  aplot                  0.3.2     2026-09-22 [1] CRAN (R 4.4.3)
##  ashr                   2.2-63    2023-08-21 [1] CRAN (R 4.4.3)
##  Biobase              * 2.66.0    2024-10-29 [1] Bioconductor 3.20 (R 4.4.1)
##  BiocGenerics         * 0.52.0    2024-10-29 [1] Bioconductor 3.20 (R 4.4.1)
##  BiocManager            1.30.27   2025-11-14 [1] CRAN (R 4.4.3)
##  BiocParallel         * 1.40.2    2025-04-16 [1] Bioconductor
##  Biostrings             2.74.1    2024-12-16 [1] Bioconductor 3.20 (R 4.4.2)
##  bit                    4.6.0     2025-03-06 [1] CRAN (R 4.4.3)
##  bit64                  4.8.6     2026-09-01 [1] CRAN (R 4.4.3)
##  bitops                 1.1-0     2026-07-30 [1] CRAN (R 4.4.3)
##  blob                   1.3.0     2026-01-14 [1] CRAN (R 4.4.3)
##  bslib                  0.12.0    2026-08-04 [1] CRAN (R 4.4.3)
##  cachem                 1.1.0     2024-05-16 [1] CRAN (R 4.4.3)
##  cellranger             1.1.0     2016-07-27 [1] CRAN (R 4.4.3)
##  cli                    3.6.6     2026-04-09 [1] CRAN (R 4.4.3)
##  clusterProfiler      * 4.14.6    2025-02-27 [1] Bioconductor 3.20 (R 4.4.2)
##  codetools              0.2-20    2024-03-31 [2] CRAN (R 4.4.3)
##  colorspace             2.1-3     2026-07-12 [1] CRAN (R 4.4.3)
##  cowplot                1.2.0     2025-07-07 [1] CRAN (R 4.4.3)
##  crayon                 1.5.3     2024-06-20 [1] CRAN (R 4.4.3)
##  curl                   8.0.0     2026-08-25 [1] CRAN (R 4.4.3)
##  data.table             1.18.6.1  2026-08-24 [1] CRAN (R 4.4.3)
##  DBI                    1.3.0     2026-02-25 [1] CRAN (R 4.4.3)
##  DelayedArray           0.32.0    2024-10-29 [1] Bioconductor 3.20 (R 4.4.1)
##  DESeq2               * 1.46.0    2024-10-29 [1] Bioconductor 3.20 (R 4.4.1)
##  digest                 0.6.39    2025-11-19 [1] CRAN (R 4.4.3)
##  DOSE                   4.0.1     2025-03-27 [1] Bioconductor 3.20 (R 4.4.3)
##  dplyr                * 1.2.1     2026-04-03 [1] CRAN (R 4.4.3)
##  enrichplot             1.26.6    2025-01-09 [1] Bioconductor 3.20 (R 4.4.2)
##  evaluate               1.0.5     2025-08-27 [1] CRAN (R 4.4.3)
##  farver                 2.1.2     2024-05-13 [1] CRAN (R 4.4.3)
##  fastmap                1.2.0     2024-05-15 [1] CRAN (R 4.4.3)
##  fastmatch              1.1-8     2026-01-17 [1] CRAN (R 4.4.3)
##  fgsea                  1.32.4    2025-03-20 [1] Bioconductor 3.20 (R 4.4.3)
##  formatR                1.14      2023-01-17 [1] CRAN (R 4.4.3)
##  fs                   * 2.1.0     2026-04-18 [1] CRAN (R 4.4.3)
##  futile.logger        * 1.4.9     2025-12-29 [1] CRAN (R 4.4.3)
##  futile.options         1.0.1     2018-04-20 [1] CRAN (R 4.4.0)
##  generics               0.1.4     2025-05-09 [1] CRAN (R 4.4.3)
##  GenomeInfoDb         * 1.42.3    2025-01-27 [1] Bioconductor 3.20 (R 4.4.2)
##  GenomeInfoDbData       1.2.13    2025-04-02 [1] Bioconductor
##  GenomicRanges        * 1.58.0    2024-10-29 [1] Bioconductor 3.20 (R 4.4.1)
##  ggfun                  0.2.1     2026-07-02 [1] CRAN (R 4.4.3)
##  ggplot2              * 4.0.3     2026-04-22 [1] CRAN (R 4.4.3)
##  ggplotify              0.1.3     2025-09-20 [1] CRAN (R 4.4.3)
##  ggrepel              * 0.9.8     2026-03-17 [1] CRAN (R 4.4.3)
##  ggtangle               0.1.3     2026-09-24 [1] CRAN (R 4.4.3)
##  ggtree                 3.14.0    2024-10-29 [1] Bioconductor 3.20 (R 4.4.1)
##  glue                   1.8.1     2026-04-17 [1] CRAN (R 4.4.3)
##  GO.db                  3.20.0    2025-07-21 [1] Bioconductor
##  GOSemSim               2.32.0    2024-10-29 [1] Bioconductor 3.20 (R 4.4.1)
##  gprofiler2           * 0.2.4     2025-11-11 [1] CRAN (R 4.4.3)
##  gridGraphics           0.5-1     2020-12-13 [1] CRAN (R 4.4.3)
##  gson                   0.2.1     2026-08-04 [1] CRAN (R 4.4.3)
##  gtable                 0.3.6     2024-10-25 [1] CRAN (R 4.4.3)
##  hms                    1.1.4     2025-10-17 [1] CRAN (R 4.4.3)
##  htmltools              0.5.9     2025-12-04 [1] CRAN (R 4.4.3)
##  htmlwidgets            1.6.4     2023-12-06 [1] CRAN (R 4.4.3)
##  httr                   1.4.9     2026-09-01 [1] CRAN (R 4.4.3)
##  igraph                 2.3.3     2026-06-26 [1] CRAN (R 4.4.3)
##  invgamma               1.2       2025-07-02 [1] CRAN (R 4.4.3)
##  IRanges              * 2.40.1    2024-12-05 [1] Bioconductor 3.20 (R 4.4.2)
##  irlba                  2.3.7     2026-01-30 [1] CRAN (R 4.4.3)
##  jquerylib              0.1.4     2021-04-26 [1] CRAN (R 4.4.3)
##  jsonlite               2.0.0     2025-03-27 [1] CRAN (R 4.4.3)
##  KEGGREST               1.46.0    2024-10-29 [1] Bioconductor 3.20 (R 4.4.1)
##  knitr                  1.52      2026-09-06 [1] CRAN (R 4.4.3)
##  labeling               0.4.3     2023-08-29 [1] CRAN (R 4.4.0)
##  lambda.r               1.2.4     2019-09-18 [1] CRAN (R 4.4.3)
##  lattice                0.23-1    2026-08-12 [2] CRAN (R 4.4.3)
##  lazyeval               0.2.3     2026-04-04 [1] CRAN (R 4.4.3)
##  lifecycle              1.0.5     2026-01-08 [1] CRAN (R 4.4.3)
##  locfit                 1.5-9.12  2025-03-05 [1] CRAN (R 4.4.3)
##  magrittr               2.0.5     2026-04-04 [1] CRAN (R 4.4.3)
##  MASS                   7.3-66    2026-07-15 [2] CRAN (R 4.4.3)
##  Matrix                 1.7-6     2026-07-25 [2] CRAN (R 4.4.3)
##  MatrixGenerics       * 1.18.1    2025-01-09 [1] Bioconductor 3.20 (R 4.4.2)
##  matrixStats          * 1.5.0     2025-01-07 [1] CRAN (R 4.4.3)
##  memoise                2.0.1     2021-11-26 [1] CRAN (R 4.4.3)
##  mixsqp                 0.3-54    2023-12-20 [1] CRAN (R 4.4.3)
##  nlme                   3.1-171   2026-09-01 [2] CRAN (R 4.4.3)
##  org.At.tair.db       * 3.20.0    2025-07-21 [1] Bioconductor
##  otel                   0.2.0     2025-08-29 [1] CRAN (R 4.4.3)
##  patchwork              1.3.2     2025-08-25 [1] CRAN (R 4.4.3)
##  pheatmap             * 1.0.13    2025-06-05 [1] CRAN (R 4.4.3)
##  pillar                 1.11.1    2025-09-17 [1] CRAN (R 4.4.3)
##  pkgconfig              2.0.3     2019-09-22 [1] CRAN (R 4.4.3)
##  plotly                 4.12.1    2026-07-22 [1] CRAN (R 4.4.3)
##  plyr                   1.8.9     2023-10-02 [1] CRAN (R 4.4.3)
##  png                    0.1-9     2026-03-15 [1] CRAN (R 4.4.3)
##  purrr                * 1.2.2     2026-04-10 [1] CRAN (R 4.4.3)
##  qvalue                 2.38.0    2024-10-29 [1] Bioconductor 3.20 (R 4.4.1)
##  R.methodsS3            1.8.2     2022-06-13 [1] CRAN (R 4.4.0)
##  R.oo                   1.27.1    2025-05-02 [1] CRAN (R 4.4.3)
##  R.utils                2.13.0    2025-02-24 [1] CRAN (R 4.4.3)
##  R6                     2.6.1     2025-02-15 [1] CRAN (R 4.4.3)
##  ragg                   1.5.2     2026-03-23 [1] CRAN (R 4.4.3)
##  rappdirs               0.3.4     2026-01-17 [1] CRAN (R 4.4.3)
##  RColorBrewer           1.1-3     2022-04-03 [1] CRAN (R 4.4.0)
##  Rcpp                   1.1.2     2026-07-05 [1] CRAN (R 4.4.3)
##  RCurl                  1.98-1.20 2026-08-21 [1] CRAN (R 4.4.3)
##  readr                * 2.2.0     2026-02-19 [1] CRAN (R 4.4.3)
##  readxl               * 1.5.0.1   2026-09-16 [1] CRAN (R 4.4.3)
##  reshape2               1.4.5     2025-11-12 [1] CRAN (R 4.4.3)
##  rlang                  1.3.0     2026-07-05 [1] CRAN (R 4.4.3)
##  rmarkdown              2.32      2026-09-01 [1] CRAN (R 4.4.3)
##  RSQLite                3.53.3    2026-06-30 [1] CRAN (R 4.4.3)
##  rstudioapi             0.19.0    2026-06-11 [1] CRAN (R 4.4.3)
##  S4Arrays               1.6.0     2024-10-29 [1] Bioconductor 3.20 (R 4.4.1)
##  S4Vectors            * 0.44.0    2024-10-29 [1] Bioconductor 3.20 (R 4.4.1)
##  S7                     0.2.2     2026-04-22 [1] CRAN (R 4.4.3)
##  sass                   0.4.10    2025-04-11 [1] CRAN (R 4.4.3)
##  scales               * 1.4.0     2025-04-24 [1] CRAN (R 4.4.3)
##  sessioninfo            1.2.4     2026-06-04 [1] CRAN (R 4.4.3)
##  snow                   0.4-4     2021-10-27 [1] CRAN (R 4.4.0)
##  SparseArray            1.6.2     2025-02-20 [1] Bioconductor 3.20 (R 4.4.2)
##  SQUAREM                2026.1    2026-03-12 [1] CRAN (R 4.4.3)
##  stringi                1.8.9     2026-08-04 [1] CRAN (R 4.4.3)
##  stringr                1.6.0     2025-11-04 [1] CRAN (R 4.4.3)
##  SummarizedExperiment * 1.36.0    2024-10-29 [1] Bioconductor 3.20 (R 4.4.1)
##  svglite              * 2.2.2     2025-10-21 [1] CRAN (R 4.4.3)
##  systemfonts            1.3.2     2026-03-05 [1] CRAN (R 4.4.3)
##  textshaping            1.0.5     2026-03-06 [1] CRAN (R 4.4.3)
##  tibble                 3.3.1     2026-01-11 [1] CRAN (R 4.4.3)
##  tidyr                * 1.3.2     2025-12-19 [1] CRAN (R 4.4.3)
##  tidyselect             1.2.1     2024-03-11 [1] CRAN (R 4.4.3)
##  tidytree               0.4.8     2026-07-02 [1] CRAN (R 4.4.3)
##  treeio                 1.30.0    2024-10-29 [1] Bioconductor 3.20 (R 4.4.1)
##  truncnorm              1.0-9     2023-03-20 [1] CRAN (R 4.4.3)
##  tximport             * 1.34.0    2024-10-29 [1] Bioconductor 3.20 (R 4.4.1)
##  tzdb                   0.5.0     2025-03-15 [1] CRAN (R 4.4.3)
##  UCSC.utils             1.2.0     2024-10-29 [1] Bioconductor 3.20 (R 4.4.1)
##  vctrs                  0.7.3     2026-04-11 [1] CRAN (R 4.4.3)
##  VennDiagram          * 1.8.2     2026-01-11 [1] CRAN (R 4.4.3)
##  viridisLite            0.4.3     2026-02-04 [1] CRAN (R 4.4.3)
##  vroom                  1.7.1     2026-03-31 [1] CRAN (R 4.4.3)
##  withr                  3.0.3     2026-06-19 [1] CRAN (R 4.4.3)
##  xfun                   0.61      2026-09-16 [1] CRAN (R 4.4.3)
##  XVector                0.46.0    2024-10-29 [1] Bioconductor 3.20 (R 4.4.1)
##  yaml                   2.3.12    2025-12-10 [1] CRAN (R 4.4.3)
##  yulab.utils            0.2.5     2026-08-29 [1] CRAN (R 4.4.3)
##  zlibbioc               1.52.0    2024-10-29 [1] Bioconductor 3.20 (R 4.4.1)
## 
##  [1] C:/Users/dalens/AppData/Local/R/win-library/4.4
##  [2] C:/Program Files/R/R-4.4.3/library
##  * ── Packages attached to the search path.
## 
## ──────────────────────────────────────────────────────────────────────────────────────────────────────────────────────────────────────────────────────────────────────────────────
```
